# Mega-scale experimental analysis of protein folding stability in biology and protein design

**DOI:** 10.1101/2022.12.06.519132

**Authors:** Kotaro Tsuboyama, Justas Dauparas, Jonathan Chen, Elodie Laine, Yasser Mohseni Behbahani, Jonathan J. Weinstein, Niall M. Mangan, Sergey Ovchinnikov, Gabriel J. Rocklin

## Abstract

Advances in DNA sequencing and machine learning are illuminating protein sequences and structures on an enormous scale. However, the energetics driving folding are invisible in these structures and remain largely unknown. The hidden thermodynamics of folding can drive disease, shape protein evolution, and guide protein engineering, and new approaches are needed to reveal these thermodynamics for every sequence and structure. We present cDNA display proteolysis, a new method for measuring thermodynamic folding stability for up to 900,000 protein domains in a one-week experiment. From 1.8 million measurements in total, we curated a set of ~850,000 high-quality folding stabilities covering all single amino acid variants and selected double mutants of 354 natural and 188 de novo designed protein domains 40-72 amino acids in length. Using this immense dataset, we quantified (1) environmental factors influencing amino acid fitness, (2) thermodynamic couplings (including unexpected interactions) between protein sites, and (3) the global divergence between evolutionary amino acid usage and protein folding stability. We also examined how our approach could identify stability determinants in designed proteins and evaluate design methods. The cDNA display proteolysis method is fast, accurate, and uniquely scalable, and promises to reveal the quantitative rules for how amino acid sequences encode folding stability.

**One-Sentence Summary:** Massively parallel measurement of protein folding stability by cDNA display proteolysis

## Main Text

Protein sequences vary by more than ten orders of magnitude in thermodynamic folding stability (the ratio of unfolded to folded molecules at equilibrium) (*1, 2*). Even single point mutations that alter stability can have profound effects on health and disease (*3–5*), pharmaceutical development (*6–8*), and protein evolution (*9–13*). Thousands of point mutants have been individually studied over decades to quantify the determinants of stability (*14, 15*), but these studies highlight a challenge: similar mutations can have widely varying effects in different protein contexts, and these subtleties remain difficult to predict despite substantial effort (*16, 17*). In fact, even as deep learning models have achieved transformative accuracy at protein structure prediction (*18–21*) progress in modeling folding stability has arguably stalled (*22–24*). New high-throughput experiments have the potential to transform our understanding of stability by quantifying the effects of mutations across a vast number of protein contexts, revealing new biophysical insights and empowering modern machine learning methods.

Here, we present a powerful new high-throughput stability assay along with a uniquely massive dataset of 851,552 folding stability measurements. Our new method - cDNA display proteolysis - combines the strengths of cell-free molecular biology and next-generation sequencing and requires no on-site equipment larger than a qPCR machine. Assaying one library (up to 900,000 sequences in our experiments) requires one week and only ~$2,000 in reagents, excluding the cost of DNA synthesis and sequencing. Compared to mass spectrometry-based high-throughput stability assays (*25–28*), cDNA display proteolysis achieves a 100-fold larger scale and can easily be applied to study mutational libraries that pose difficulties for proteomics. Compared to our earlier yeast display proteolysis method (*29*), cDNA display proteolysis resolves a wider dynamic range of stability and is more reproducible even at a 50-fold larger experimental scale. Large-scale proteolysis data have already played a key role in the development of machine learning methods for protein design and protein biophysics (*30–36*). The cDNA display proteolysis method massively expands this capability and has the potential to expand our knowledge of stability to the scale of all known small domains.

Our new dataset of 851,552 absolute folding stabilities is unique in size and character. Current thermodynamic databases contain a skewed assortment of mutations measured under many varied conditions (*14*). In contrast, our new dataset comprehensively measures all single mutants for 354 natural domains and 188 designed proteins - including single deletions and two insertions at each position - all under identical conditions. Our dataset also includes comprehensive double mutations at 595 site pairs spread across 208 domains (a total of 222,265 double mutants). By maintaining uniform experimental conditions, our data can be used to examine the determinants of *absolute* folding stability in addition to the effects of mutations. Using our unique dataset, we investigated how individual amino acids and pairs of amino acids contribute to folding stability (Figs. 3 and 4) as well as how selection for stability interacts with other selective pressures in natural protein domains (Figs. 5 and 6). We also explored how our unique scale of data can be applied in protein design (Fig. 7).

### Massively parallel measurement of folding stability by cDNA display proteolysis

Proteases typically cleave unfolded proteins more quickly than folded ones, and proteolysis assays have been used for decades to measure folding stability (*37*) and select for high stability proteins (*38, 39*). In 2017, we introduced the high-throughput yeast display proteolysis method for measuring folding stability using next generation sequencing (*29, 40–46*). To improve the scale, precision, speed, and cost of stability measurements, we developed cDNA display proteolysis. Each experiment begins with a DNA library. Here, we employ synthetic DNA oligo pools where each oligo encodes one test protein. The DNA library is transcribed and translated using cell-free cDNA display (*47*), based on mRNA display (*48, 49*), resulting in proteins that are attached at the C-terminus to their cDNA. We then incubate the protein-cDNA complexes with different concentrations of protease, quench the reactions, and pull down the proteins using an N-terminal PA tag (Fig. 1A). Intact (protease-resistant) proteins will also carry their C-terminal cDNA. Finally, we determine the relative amounts of all proteins in the surviving pool at each protease concentration by deep sequencing (Fig. 1D). To control for any effects of protease specificity, we perform separate experiments with two orthogonal proteases: trypsin (targeting basic amino acids) and chymotrypsin (targeting aromatic amino acids).

**Fig. 1.**
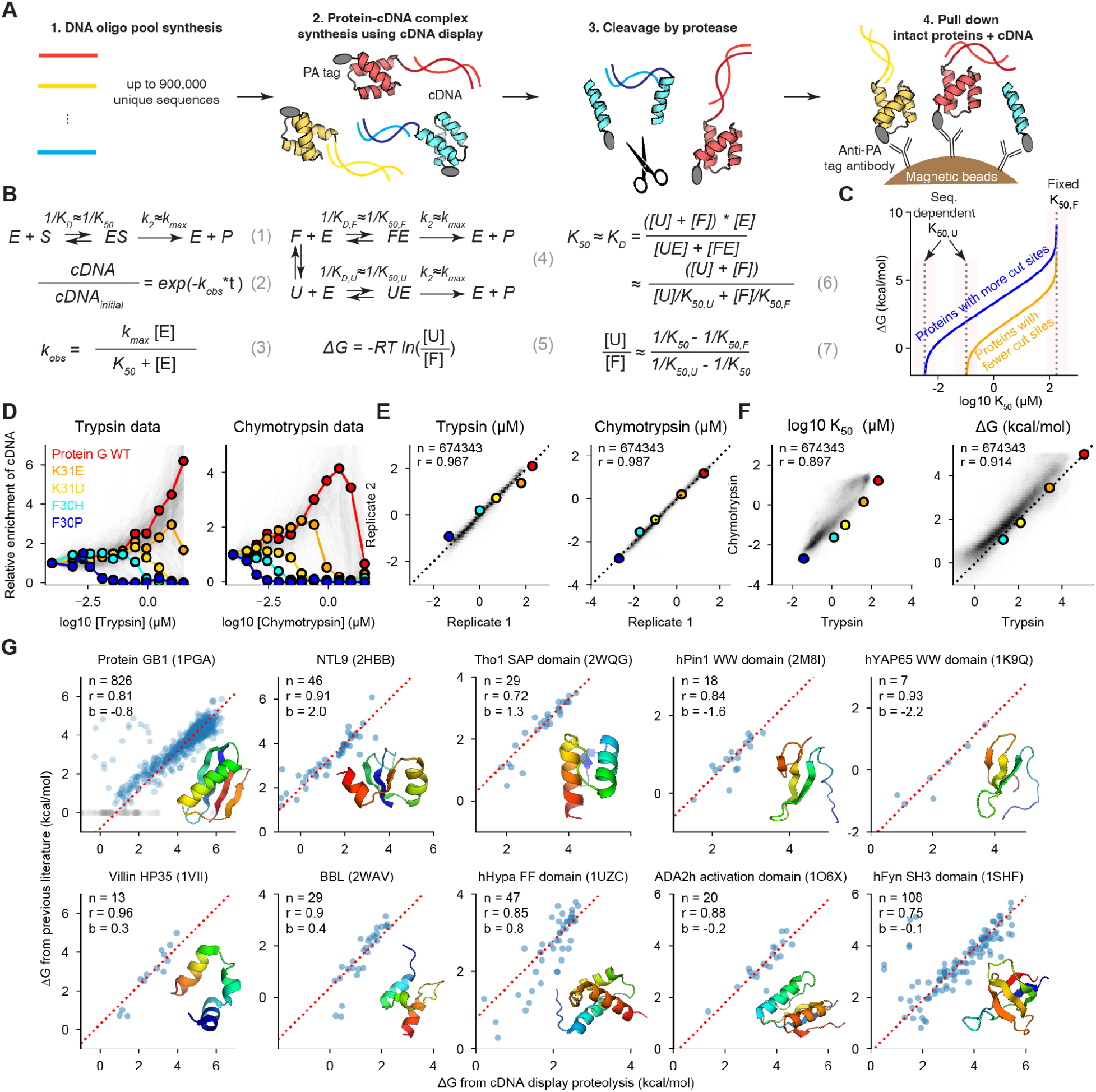
cDNA display enables massively parallel measurement of protein folding stability. **(A)** A DNA oligo library is expressed using cell-free cDNA display, producing proteins with an N-terminal PA tag and C-terminal covalent attachment to cDNA. Protease cleavage separates the cDNA from the PA tag. After protease challenge, magnetic beads with anti-PA antibodies pull down protein N-termini and intact proteins carry along their cDNA. cDNA is then amplified and sequenced to quantify the intact fraction of each protein. **(B)** Thermodynamic model of proteolysis based on single turnover kinetics. Protease enzymes (E) and protein substrates (S) form an ES complex to produce cleaved protein products (P) (1). We model the cleavage as a first-order reaction (2) according to single turnover kinetics (3). We use an identical k_max_ for all sequences and fit each sequence’s K_50_ concentration to our data. Proteins are normally cleaved in the unfolded (U) state but can also be cleaved in the folded (F) state (e.g. by cleaving the PA tag) (4). We determine the folding equilibrium using each sequence’s measured K_50_, a predicted sequence-specific K_50_ for the unfolded state (K_50,U_), and a universal K_50_ for the folded state (K_50,F_). **(C)** Relationship between K_50_ and ΔG for a protein with fewer cut sites (yellow) and a protein with more cut sites (blue). When K_50_ approaches K_50,U_ or K_50,F_ (red shaded regions), ΔG becomes very sensitive to K_50_ and its uncertainty increases relative to the uncertainty in K_50_. **(D)** PA tag pulldown at increasing protease concentrations separates proteins by stability. Each sequence of Protein GB1 variants in a library is shown as a gray line tracking its change in population fraction relative to that in the pre-selection library (enrichment). Enrichment traces for the wild-type and four mutants are highlighted in color. **(E)** Reproducibility of K_50_ from two replicates of the proteolysis procedure, after filtering for data quality and range (see Methods). The K_50_ density is shown in gray with the proteins from (D) highlighted in color. **(F)** Consistency of K_50_ (left) and ΔG (right) between trypsin and chymotrypsin for one library (black), highlighting the five proteins shown in (D). **(G)** Our high-throughput ΔG measurements are consistent with previously published stability data from purified protein samples for wild types and mutants of 10 domains. The red dashed line represents the Y = X+b (intercept) line. Gray points (Protein GB1) indicate ‘no data’ in the previous paper. See Table. S2 and Fig. S3 for analysis of the intercepts

We inferred the protease stability of all sequences from our sequencing counts using a Bayesian model of the experimental procedure. We modeled protease cleavage using single turnover kinetics (*50, 51*) (Fig. 1B eqs. 1 to 3, Fig. S1, and Supplementary Text for the derivation) because we assume the enzyme is in excess over all substrates (up to ~20 pM of substrate based on previous estimates (*47*) versus 141 pM for the lowest concentration of protease). To parameterize the model, we used a universal k_max_ cleavage rate for all sequences (Fig. S1) and used our sequencing data to infer a unique K_50_ for each sequence (the protease concentration at which the cleavage rate is one-half k_max_, see Methods). The K_50_ values inferred by the model were consistent between two replicates of the proteolysis procedure (R = 0.97 for trypsin and 0.99 for chymotrypsin for ~84% of sequences in a pool of 806,640 sequences after filtering based on confidence and dynamic range; Fig. 1E).

To infer each sequence’s thermodynamic folding stability (ΔG for unfolding), we used a kinetic model that separately considers idealized folded (F) and unfolded (U) states (Fig. 1B eq. 4). We model both states using the same single-turnover equations as before (Fig. 1B eq. 3), with separate K_50_ protease concentrations for each state (K_50,F_ and K_50,U_) and a shared k_max_. We assume that cleavage in the folded state exclusively occurs outside the folded domain (e.g. in the N-terminal PA tag added to all sequences), so we use an identical K_50,F_ for all sequences. In contrast, K_50,U_ reflects an individual sequence’s unique protease susceptibility in the unfolded state, which depends on its potential cleavage sites. We inferred K_50,U_ for each sequence using a position-specific scoring matrix (PSSM) model of protease cleavage parameterized using measurements of 64,238 scrambled sequences (sequences with a high probability of being fully unfolded, Fig. S2; see also Methods). Finally, we assume that folding, unfolding, and enzyme binding are all in rapid equilibrium relative to cleavage, implying that K_50,U_, K_50,F_, and the overall K_50_ can be approximated by the enzyme-substrate equilibrium dissociation constants for each state (Fig. 1B eq. 6). Although these approximations will not be universally accurate, they are appropriate for the small domains examined here and facilitate consistent analysis of all test sequences. With these approximations, we can express a sequence’s ΔG in terms of the universal K_50,F_, its inferred K_50,U_, and its experimentally measured K_50_ (Fig. 1B eq. 5 and 7, and Supplementary Text for the derivation). For most analysis we combine our independent trypsin and chymotrypsin data into a single overall ΔG estimate (See Methods). Based on our kinetic model, (1) stability (ΔG) will be underestimated if significant cleavage occurs inside the test domain in the folded state, (2) stability can be over- or under-estimated depending on the accuracy of K_50,U_ (independent measurements with trypsin and chymotrypsin help correct this), and (3) ΔG values become unreliable if K_50_ approaches K_50,F_ or K_50,U_ (Fig. 1C).

### Folding stabilities from cDNA display proteolysis are consistent with traditional experiments on purified proteins

In Fig. 1G, we compare stabilities measured by cDNA display proteolysis to previous results from experiments on purified protein samples for 1,143 variants of ten proteins (*52–65*). All Pearson correlations are above 0.7. Our stability measurements for these 1,143 sequences were all performed in libraries of 244,000–900,000 total sequences. Although several sets of mutants show systematic offsets (y-intercept values) between literature values and our measurements, these offsets correlate with temperature differences between experimental conditions (with the exception of the N-terminal domain of Ribosomal Protein L9 (2HBB), Fig. S3, see Table S2 for all experimental conditions). We also noticed several variants of Protein GB1 appear unstable in our data but stable in the previous experiments (*52*). Our structural analysis of these mutations suggests that our measurements are more likely to be correct (Fig. S4). Overall, the consistency between our cDNA display proteolysis data and traditional biophysical measurements establishes that (1) small domains are cleaved mainly in the globally unfolded state, (2) our method can reliably measure these cleavage rates on a massive scale, and (3) our unfolded state model can remove protease-specific effects to attain accurate quantitative folding stability measurements.

### Comprehensive mutational analysis across designed and natural protein domains

To systematically examine how individual residues influence folding stability, we used cDNA display proteolysis to measure stability for all single substitutions, deletions, and Gly and Ala insertions in 983 natural and designed domains. We chose our natural domains to cover almost all of the small monomeric domains in the Protein Data Bank (30-72 amino acids in length). Our designed domains included (1) previous Rosetta designs with ααα, αββα, βαββ, and ββαββ topologies (40-43 a.a.) (*29, 66*), (2) new ββαα proteins designed using Rosetta (47 a.a.), and (3) new domains designed by trRosetta hallucination (46 to 69 a.a.) (*42, 67*). We collected these data using four giant synthetic DNA oligonucleotide libraries and obtained K_50_ values for 2,520,337 sequences; 1,844,548 of these measurements are included here. K_50_ values were reproducible across libraries (Fig. S5). Oligo pools were synthesized by Agilent Technologies (one 244,000-sequence library, length 170 nt) and Twist Bioscience (three libraries of 696,000 - 900,000 sequences, length 250-300 nt).

Deep mutational scanning of hundreds of domains revealed several overall patterns. The largest fraction of these domains showed clear, biophysically reasonable sequence-stability relationships that were consistent between separate experiments with trypsin and chymotrypsin. However, other domains were completely unfolded, too stable to resolve, insensitive to mutation, or inconsistent between the proteases. For 42 domains that were too stable to resolve, we introduced single mutations to destabilize the wild-type sequence, then performed new mutational scanning experiments in these 121 new “wild-type” backgrounds (Fig. S6). In four domains, mutational scanning revealed trypsin-sensitive loops that could be cleaved in the folded state, leading to inconsistent stabilities between trypsin and chymotrypsin (Fig. S7). In these cases, we introduced one to two substitutions into the wild-type sequences to remove trypsin-sensitive sites, then performed new mutational scanning experiments in these alternative backgrounds. This led to consistent results between the two proteases. In total, we performed deep mutational scanning for 983 domain sequences, including both original and revised wild-type backgrounds.

Our overall categorization of all domains is shown in Fig. 2B (see Fig. S8 for inclusion criteria). Based on these categories, we assembled three curated datasets for machine learning (Fig. 2A). Our ΔΔG dataset (Dataset #1) includes 586,938 sequences (single and double mutants) from 251 natural domains and 145 designs. In this dataset, the wild-type sequence is 1.25-4.5 kcal/mol in stability so that most ΔΔG values (including for stabilizing mutants) are correctly resolved. Our ΔG dataset (Dataset #2) includes all 851,552 single and double mutants from 354 natural domains and 188 designs. In this dataset, the large majority of mutant ΔGs are accurately resolved, but the wild-type ΔG may lie outside the dynamic range, preventing accurate ΔΔG calculations. Finally, Dataset #3 includes all ~1.8 million confidently estimated K_50_ values, even when trypsin and chymotrypsin measurements produced inconsistent ΔG estimates. The main domain classes in Dataset #1 are shown in Fig. 2C; all natural domains included in Dataset #1 are listed by category in Fig. S9 (see Supplementary Materials for all wild-type sequences).

**Fig. 2.**
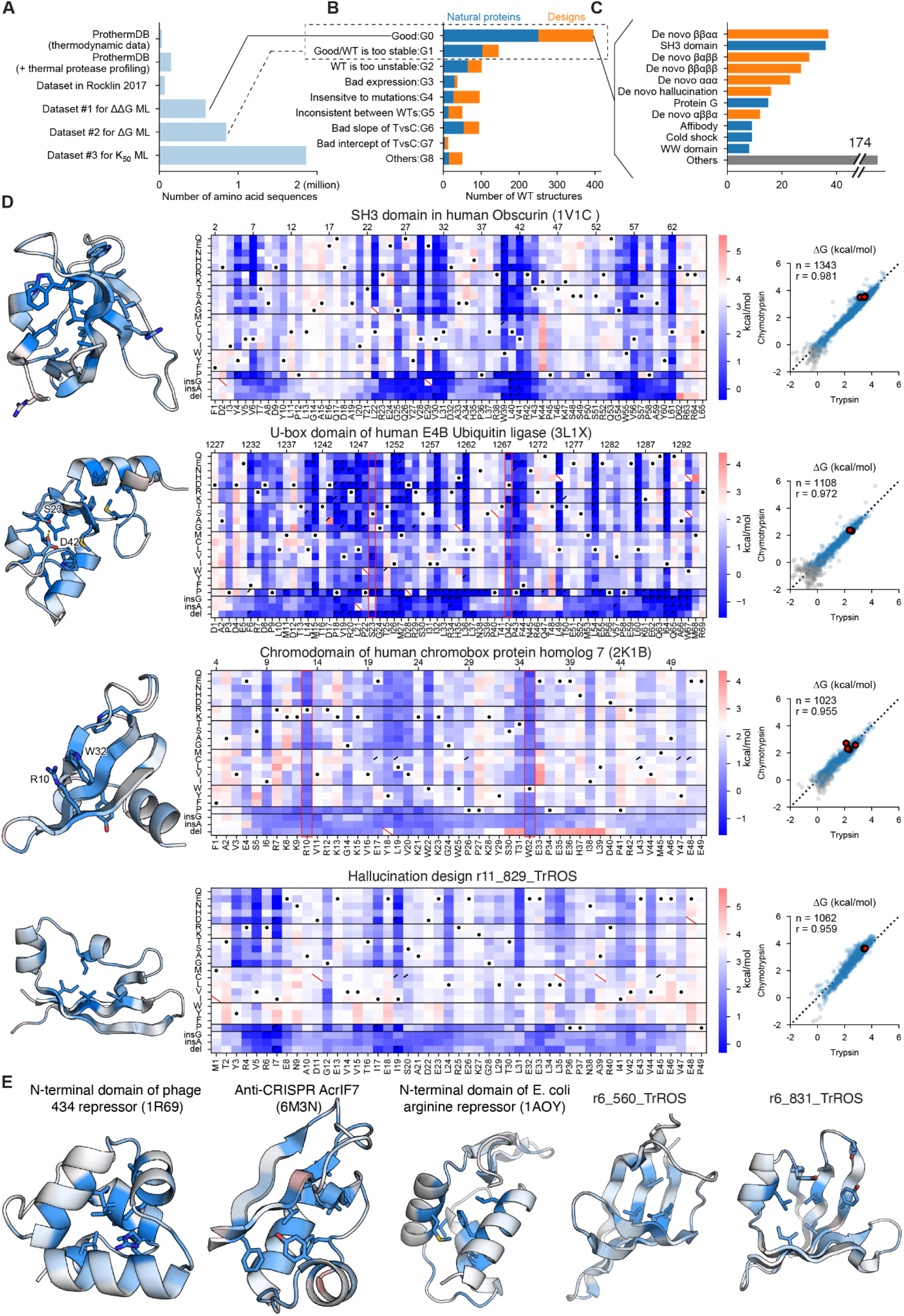
Comprehensive mutational analysis of stability in designed and natural proteins. **(A)** Comparison of the size of existing datasets and the datasets from this paper. The data of this paper are divided into three groups: datasets #1, #2, and #3, according to the quality of the data (see Table S1 and Fig. S8 for details). ML: machine learning **(B)** Classification of mutational scanning results for each wild-type sequence. The G0 group corresponds to Dataset #1, and G0 and G1 groups combined correspond to Dataset #2 in (A). (G1: Good but WT may be outside the dynamic range) **(C)** Wild-type structures classified as G0 in (B) grouped into domain families. The 11 most common domain types are shown; the remaining 174 domains are classified as “Other” (see Fig. S9). **(D)** Mutational scanning results for four domains. Left: domain structures colored by the average ΔΔG at each position; darker blue indicates mutants are more destabilizing. The structure of the design r11_829_TrROS is an AlphaFold model. Middle: Heat maps show ΔG for substitutions, deletions, and Gly and Ala insertions at each residue, with the PDB numbering shown at top and our one-indexed numbering at bottom. White represents the wild-type stability and red/blue indicate stabilizing/destabilizing mutations. Black dots indicate the wild-type amino acid, red slashes indicate missing data, and black corner slashes indicate lower confidence ΔG estimates, (95% confidence interval > 0.5 kcal/mol), including ΔG estimates near the edges of the dynamic range. Red boxes highlight the S23-D42 hydrogen bond in 3L1X and the R10-W32 cation-π interaction in 2K1B. ΔG values were fit to trypsin and chymotrypsin data together; see Methods. Right: Agreement between mutant ΔG values independently determined using assays with trypsin (x-axis) and chymotrypsin (y-axis). Multiple codon variants of the wild-type sequence are shown in red, reliable ΔG values in blue, and less reliable ΔG estimates (same as above) in gray. The black dashed line represents Y=X. Each plot shows the number of reliable points and the Pearson r-value for the blue (reliable) points. **(E)** As in (D) left, structures of five domains are shown colored by the average ΔΔG at each position; darker blue indicates mutants are more destabilizing. The two designed structures are AlphaFold models.

**Fig. 3.**
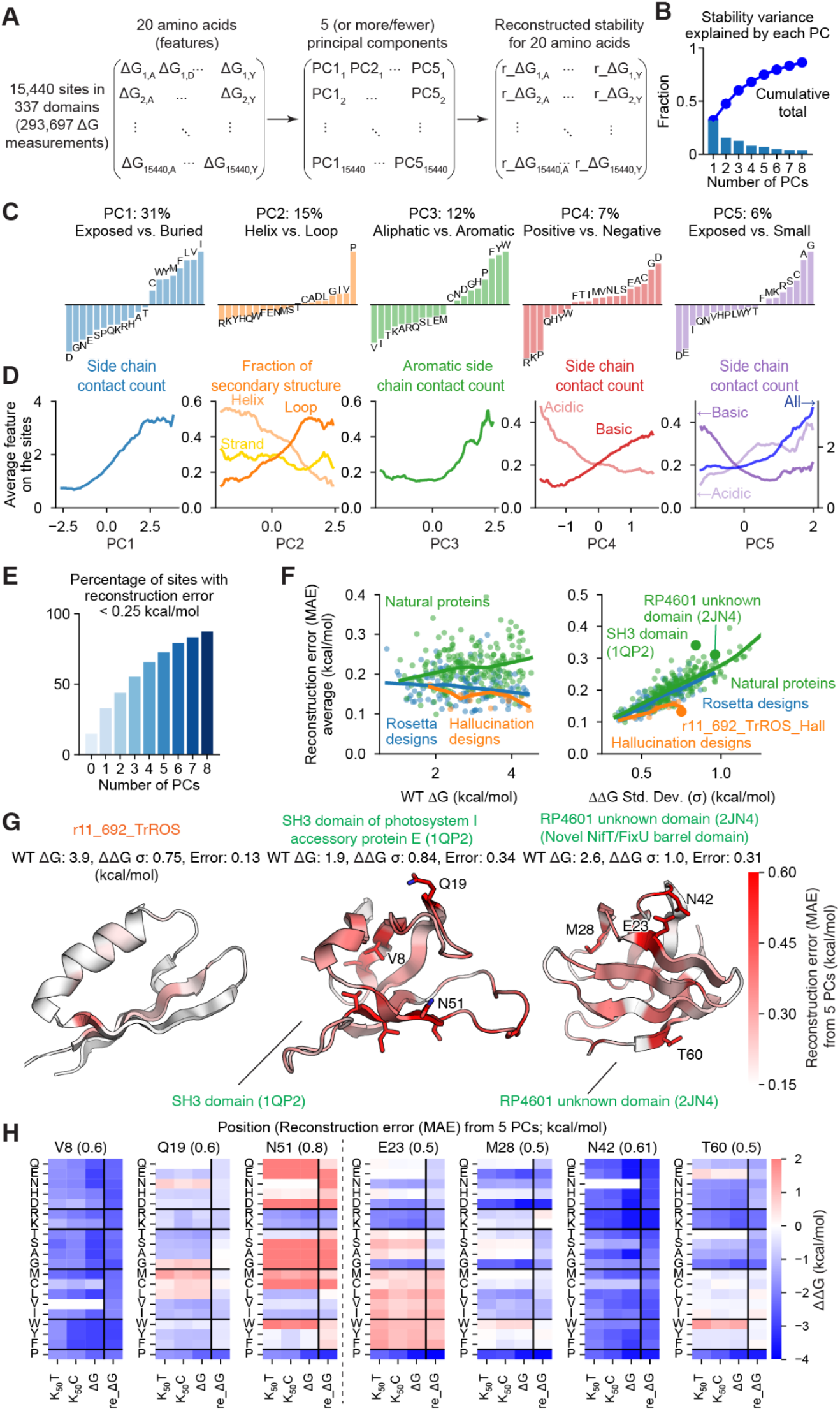
Environmental factors that determine amino acid stabilities at a position. **(A)** Principal component (PC) analysis on a matrix consisting of 15,440 observations (sites in proteins) x 20 amino acids (features) to determine the factors influencing stabilities of different amino acids. **(B)** Fraction of the total variance explained by each PC (bars) and the cumulative total (upper line). **(C)** Principal components of the stability data indicating the dominant trends for which amino acids would be stable or unstable at a site. We label each component with a biophysical interpretation and show the percent of the total variance explained by that component. **(D)** Relationships between the PC values (x-axis) for all 15,440 positions and environmental properties of the position from the three-dimensional structure (y-axis). Colored lines show each environmental feature averaged over a window of 0.25 in the units of each PC. **(E)** Fraction of sites (observations) whose stability landscapes can be reconstructed with MAE <0.25 kcal/mol using the first 1-8 principal components. **(F)** Relationship between reconstruction error using five PCs (MAE, y-axis) and wild-type stability (left, x-axis) or variance in the ΔΔG data (right, x-axis). Colors represent protein structures grouped into natural proteins (green), Rosetta designs (blue), and hallucination designs (orange). Three example proteins shown in (G) are shown as large dots. Lines show LOWESS fits. **(G)** Structures of three example proteins with each position colored by the error (MAE) between the reconstructed ΔΔG values (using 5 PCs) and the observed ΔΔG values. The left protein was designed by hallucination and each position is accurately reconstructed using only 5 PCs, whereas the middle and right natural proteins have positions with more unusual ΔΔG patterns and larger MAEs. The r11_692_TrROS structure is an AlphaFold model. **(H)** For seven positions with large MAE in the center (1QP2) and right domains (2JN4) from (G), we show the experimental trypsin and chymotrypsin K_50_ values, the ΔG values, and the reconstructed ΔG values based on the top five PCs.

**Fig. 4.**
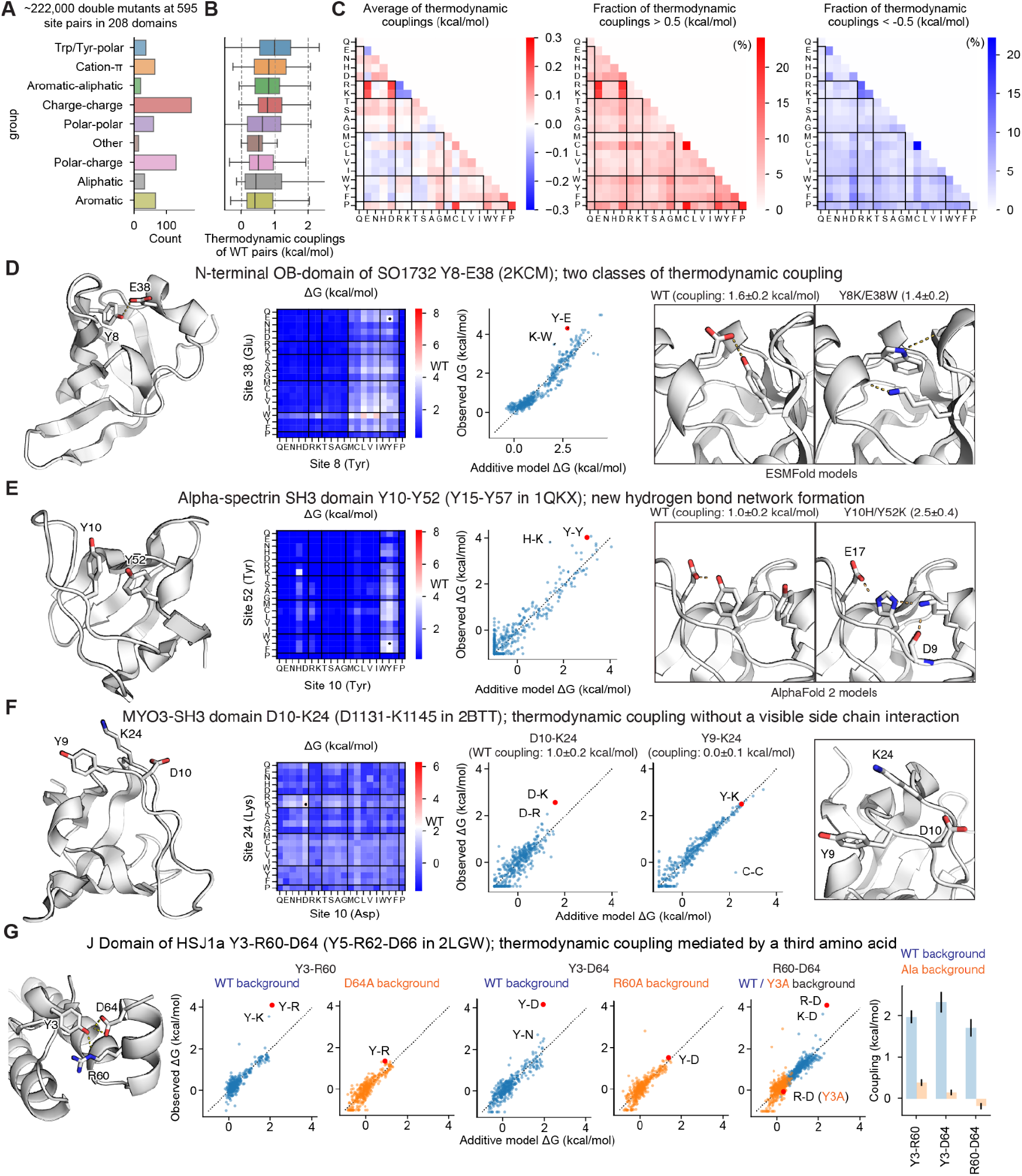
Quantifying thermodynamic coupling between amino acid pairs. **(A)** Categorization of 595 pairs of amino acids selected for exhaustive double mutant analysis. **(B)** Thermodynamic couplings of the wild-type amino acid pairs according to our additive model broken down by category. **(C)** The average thermodynamic couplings (left) and the fraction of amino acid pairs with thermodynamic coupling > 0.5 kcal/mol (middle) and < −0.5 kcal/mol (right) for all amino acid combinations (wild-type and mutant). **(D and E)** Analysis of thermodynamic coupling for two notable amino acid pairs. From left to right, we show the structure of each domain and the two positions that were mutated, the stabilities (ΔG) of all pairs of amino acids at those positions, the agreement between the stabilities from the additive model (x-axis) and the observed stabilities (y-axis) with the wild-type pair shown as a red dot, and AlphaFold or ESMFold models of amino acid pairs with strong thermodynamic couplings. Thermodynamic couplings show the observed stability minus the expected stability from the additive model; the uncertainties show the standard deviations from computing the couplings using bootstrapped samples of the 400 double mutants. **(F)** Thermodynamic coupling without a visible side chain interaction. From left to right, the structure of the MYO3 SH3 domain and notable residues; the stabilities (ΔG) of all pairs of amino acids at D10 and K24; the stabilities of double mutants in the additive model (x-axis) and experimental data (y-axis); and the zoomed structure for D10, K23, and K24. **(G)** Thermodynamic coupling mediated by a third amino acid. Exhaustive amino acid substitutions were performed for each pair of two out of the three amino acids. The same amino acid substitutions were also performed for the mutant background with the third amino acid replaced by Ala. From left to right, the AlphaFold-modeled structure of the J domain of HSJ1a with three interacting amino acids, the stabilities of double mutants in the additive model (x-axis) and experimental data (y-axis) in the wild-type background (blue) and Ala-replaced backgrounds (orange), and the thermodynamic coupling for each pair of wild-type amino acids in the wild-type background (blue) and the Ala-replaced backgrounds (orange). Substituting any of the three amino acids for Ala eliminates the thermodynamic coupling between the other two amino acids. Error bars show standard deviations from bootstrap resampling as before.

**Fig. 5.**
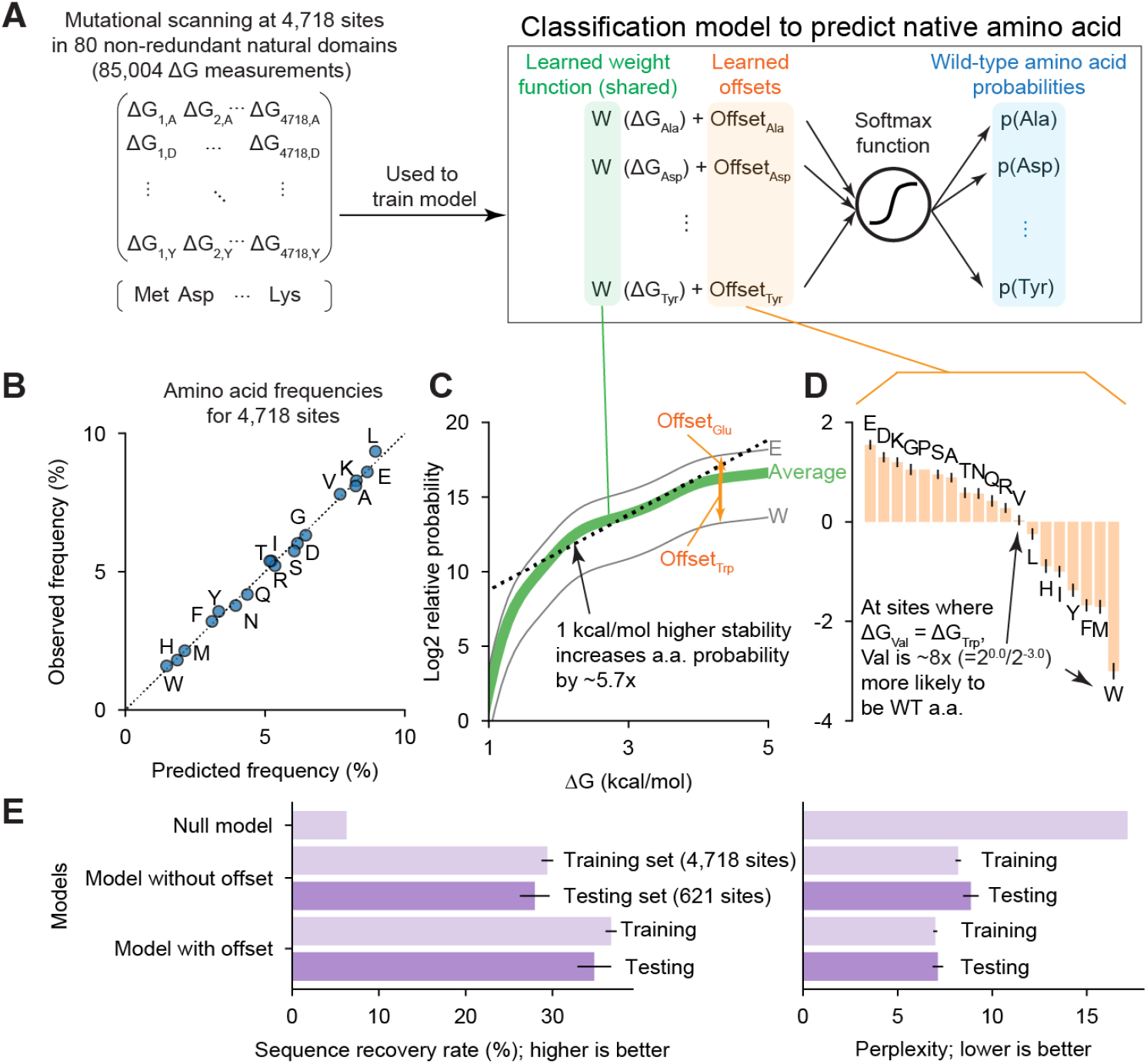
Amino acid usage in natural proteins systematically deviates from maximizing stability. **(A)** Classifier model for predicting wild-type amino acids based on the folding stabilities (ΔG) of each possible protein variant. A shared weighting function converts the stabilities of protein variants containing each amino acid into relative probabilities of those amino acids (green). The relative probability of each amino acid is further modified by a constant offset that is unique for each amino acid (orange). **(B)** Predicted and observed amino acid frequencies according to the classifier model after fitting. **(C)** The weighting function from the classifier model after fitting (green). Gray lines show the weight function after amino acid-specific offsets for Glu and Trp. In the region between 1.5 and 4 kcal/mol, the function has an approximately constant slope where a 1 kcal/mol increase in stability leads to a 5.7-fold increase in amino acid probability. **(D)** Relative offsets for 19 amino acids from the classifier model after fitting. Error bars show the standard deviation of the model posterior. **(E)** The sequence recovery rate (left) and perplexity (right) for predicting the wild-type amino acid using several models: an null model that ignores stability and always predicts amino acids at their observed frequencies, our classifier model without amino acid-specific offsets, and our full classifier model. Similar performance of the classifier model on a training set of 4,718 positions (light purple) and a testing set of 621 positions (dark purple) indicates that the model is not overfit. Error bars show standard deviations from bootstrap resampling of the sites in the training set and the testing set.

**Fig. 6.**
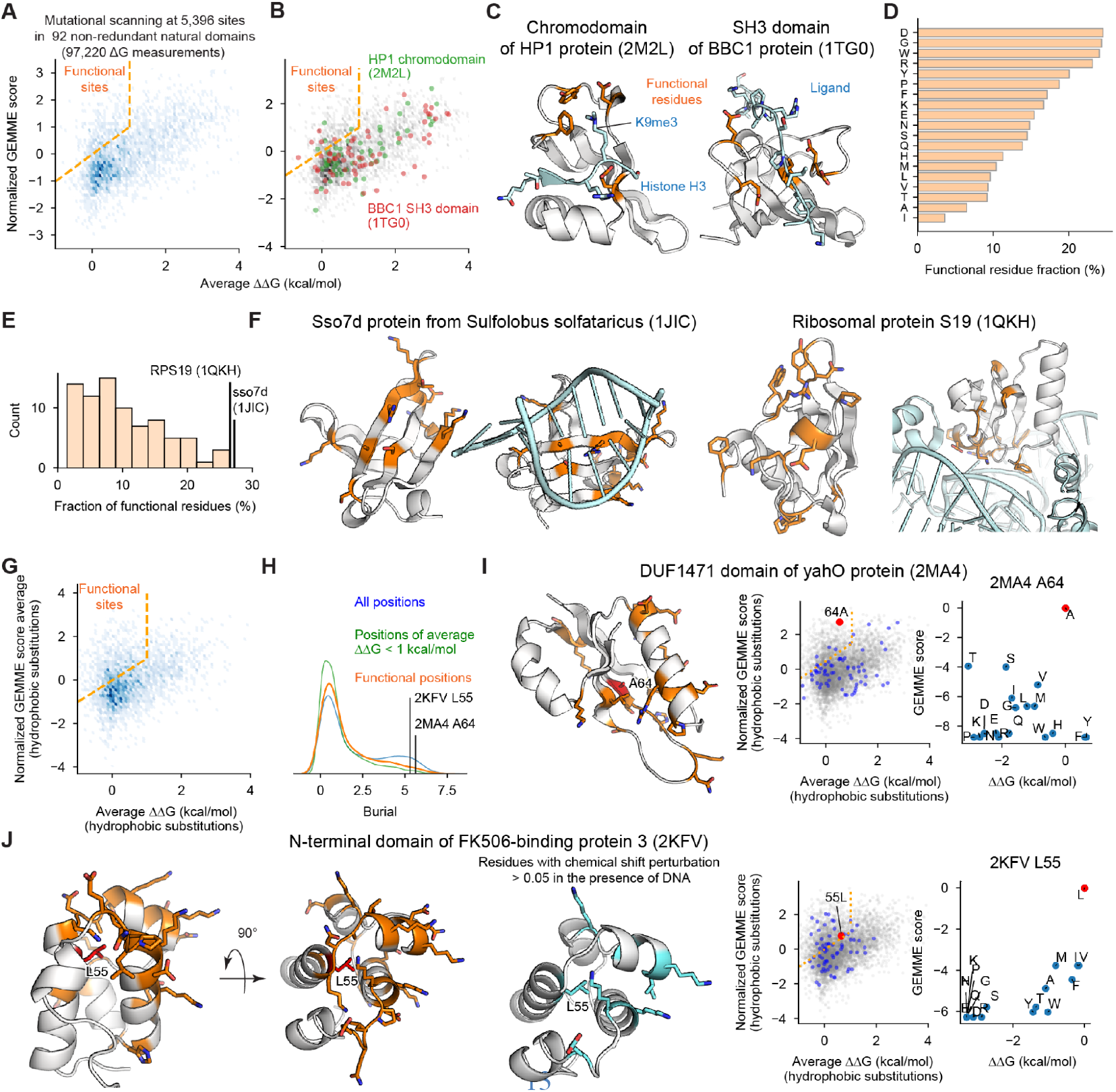
Properties of functional sites across diverse domains. **(A)** The relationship between wild-type stability (average ΔΔG for substitutions) and evolutionary-based sensitivity to substitutions (normalized averaged GEMME score). All sites above the orange dashed line are highly conserved but unimportant for stability; we define these as “functional sites”. **(B)** As in (A), highlighting positions in the HP1 chromo domain (2M2L; green) and the BBC1 SH3 domain (1TG0; red). **(C)** Structures of HP1 chromo domain and BBC1 SH3 domain (gray) and their ligands (light blue). Functional sites are shown in orange. Ligand positions were modeled based on PDB structures 1KNA (for HP1) and 2LCS (for the SH3 domain). **(D)** Amino acids are ranked by the percentage of positions where that wild-type amino acid is classified as functional, for 5,396 positions in 92 non-redundant natural domains. **(E)** The percentage of functional residues in each of the 92 non-redundant domains. **(F)** Structures of the two domains with the highest percentages of functional residues. Nucleic acids interacting with each of the structures are shown in light blue and functional residues are shown in orange. The Sso7d-DNA complex is the crystal structure 1BNZ; the S19-RNA complex is modeled based on the 4V5Y structure. **(G)** As in (A), except only considering nonpolar substitutions for calculating ΔΔG and normalized averaged GEMME score. **(H)** The distributions of burial (side chain contacts) for all sites (blue), sites where the wild-type amino acid is unimportant for stability (average ΔΔG < 1 kcal/mol) (green), and functional sites (orange). Functional sites are generally located on the surface of the protein. Two unusual buried functional residues are highlighted. **(I)** Structure of the DUF1471 domain of yahO (2MA4) with functional sites in orange and the unusual buried functional site A64 in red. Ala64 is highly conserved yet the domain is stabilized by substitutions to Tyr or Phe (positive ΔΔG, x-axis). However, Tyr and Phe are rarely found in evolution (low GEMME score, y-axis). **(J)** Left: Structure of the N-terminal domain of FK506-binding protein 3 (2KFV) with functional sites in orange and the unusual buried functional site L55 (L78 in PDB numbering) in red. Middle: Residues with chemical shift perturbations in response to DNA binding (*79*); L55 shows a perturbation despite not contacting DNA. Right: L55 is conserved (high GEMME score, y-axis) but relatively unimportant for stability (low average ΔΔG, x-axis). Substitution to Phe, Val, or Ile is thermodynamically neutral (ΔΔG near zero) but these amino acids are rarely found in evolution (low GEMME score).

**Fig. 7.**
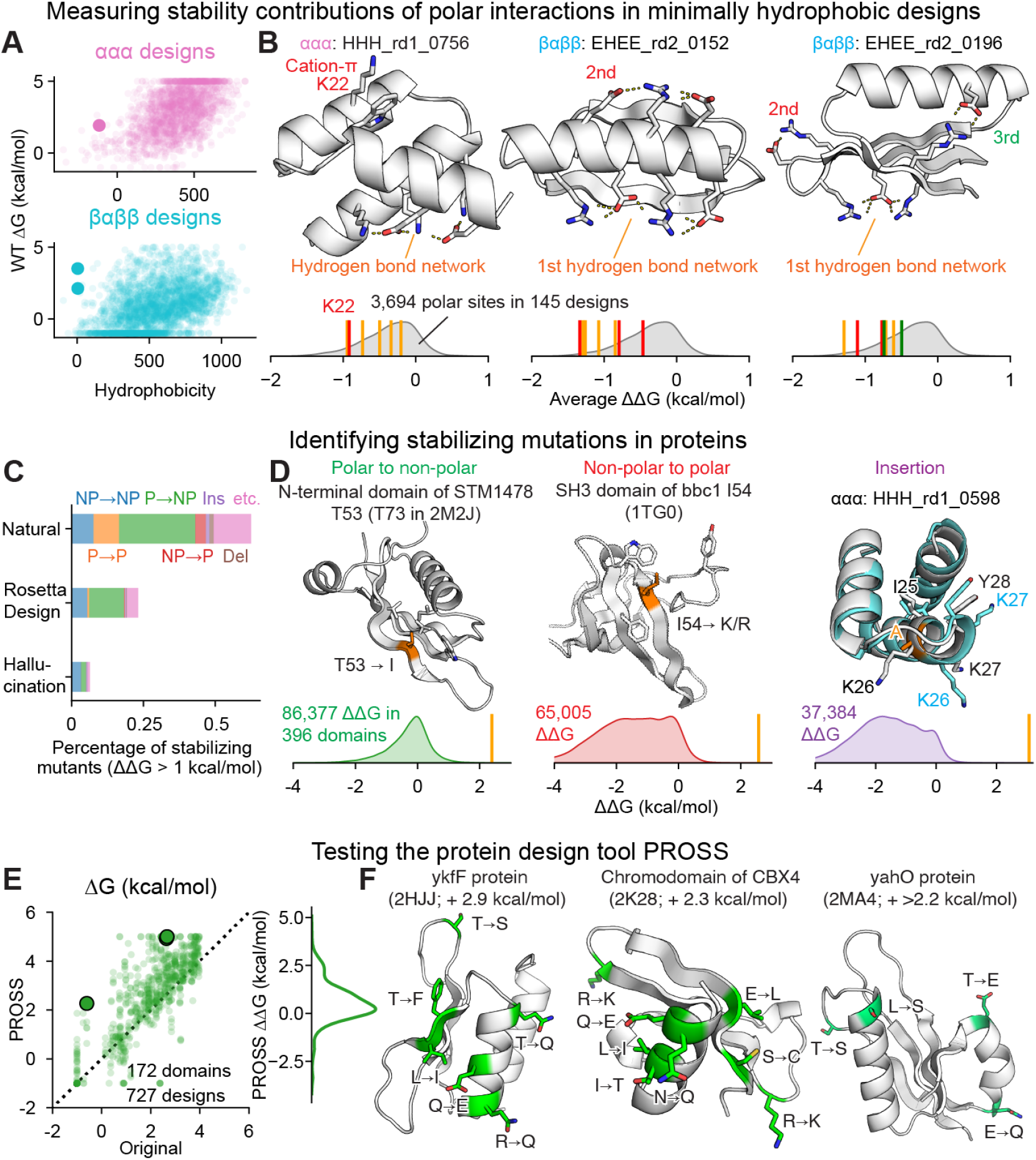
Application of screening data to protein design. **(A)** Relationship between hydrophobicity (calculated based on the previous report (Monera et al., 1995)) and folding stability (ΔG) for designed proteins (*29*). Examples from (B) are shown as large dots. **(B)** For three proteins with high folding stability and low hydrophobicity, we highlight critical hydrophilic interactions stabilizing these proteins. The gray density plots represent the average ΔΔG of substitutions at 3,694 polar sites in 145 designed domains. The colored vertical bars indicate the values for the highlighted positions. These three proteins feature polar amino acids where the average ΔΔG of substitutions is unusually destabilizing (> top 5%ile). For HHH_rd1_0756, K22 is shown as a red line; the interacting W32 is considered nonpolar and not shown. Full mutational scanning results are shown in Fig. S17. All three structures are design models reported previously (*29*), not experimental structures. **(C)** Fraction of stabilizing mutations (ΔΔG > 1 kcal/mol) found in natural domains, Rosetta designs, and hallucination designs, broken down by mutation type. NP: non-polar, P: polar, Ins: insertion, Del: deletion. **(D)** Three examples of stabilizing mutations identified by our assay, along with the distribution of ΔΔG values for these three mutation types. The highlighted mutations are indicated by vertical bars on the density plots. Full mutational scanning results are shown in Fig. S19. The structure of HHH_rd1_0598 is a design model reported previously (*29*), not an experimental structure. **(E)** Left: Testing the protein design tool PROSS (*80*). Each point shows the stability of one domain before (x-axis) and after (y-axis) redesign by PROSS. The dashed black line represents Y=X. Examples from (F) are shown as large dots. Right: Distribution of ΔG change from PROSS redesign. 40% of domains are stabilized by > 1 kcal/mol by the tool. Note that we only show 727 designs with wild-type ΔG < 4 kcal/mol. **(F)** Examples of domains stabilized by PROSS. Amino acids mutated by PROSS are shown in green on the Alphafold-generated structural model.

Mutational scanning results for nine domains are shown in Fig 2D and E. Like all mutational scans in Datasets 1 and 2, these examples show a strong consistency between independent ΔG measurements with trypsin and chymotrypsin (Pearson correlation 0.94 ± 0.04 for 542 domains in Dataset 2, median ± std.). In each structure, sites are colored according to the average effect of an amino acid substitution, with the most critical sites (where mutations are very destabilizing) colored dark blue. Most of these critical sites are in the hydrophobic core. However, our data also reveal numerous other critical interactions, such as a side chain hydrogen bond between S23 and D42 in the U-box domain of human E4B Ubiquitin ligase and a cation-π interaction between R10 and W32 in the chromodomain of human chromobox protein homolog 7 (residues have been re-numbered based on the exact sequence included in our experiments). These unique stabilizing interactions reveal the rich biophysical diversity found in our systematic exploration of stability across hundreds of domains.

### Trends in amino acid fitness at different sites and across domains

We first sought to define the major sources of variation between protein sites that determine the relative stabilities of all 20 amino acids at that site (i.e. the site’s stability landscape). To this end, we performed principal component (PC) analysis using 293,697 ΔG measurements at 15,440 sites in 337 domains from Dataset #1 after centering our data to set the average ΔG at each site to zero (Fig. 3A, B). Each principal component expresses specific properties of a site that determine which amino acids are stabilizing or destabilizing. Based on the loadings of the different amino acids onto each principal component (Fig. 3C), we interpreted the first four components to reflect amino acid hydrophobicity (PC1; 31% of the total variance explained by this PC), helical probability (PC2; 15%), aliphatic vs. aromatic favorability (PC3; 12%), and positive vs. negative charge (PC4; 7%). The fifth principal component (6%) was more complex: at one extreme were small amino acids that could be buried in dense environments, along with positively charged amino acids that can “snorkel” their charged moieties to the surface even when partially buried. At the other extreme were negatively charged amino acids that are energetically costly to bury. We interpreted this component to reflect an “ease of burial” that is orthogonal to the hydrophobic property captured by PC1. These interpretations are also consistent with the structural environments at each site, as shown in Fig. 3D. For example, the first principal component reflecting hydrophobicity is high at buried positions and low at exposed positions (Fig. 3D).

These first five principal components collectively form a coarse model of the properties of protein sites, but some sites have unique stability landscapes that cannot be accurately represented by this model. We reconstructed the stability landscapes at all sites using the first five components and examined how different sites and domains deviated from these simplified landscapes (Fig. 3E). On average, stability landscapes reconstructed using five principal components were similarly accurate (in terms of mean absolute error) for both high and low stability domains (Fig. 3F). However, as expected, these coarse reconstructions were less accurate for domains with more varied stability landscapes (domains with a higher standard deviation of ΔG for all substitutions). The coarse model was also more accurate at reconstructing the stability landscapes of de novo designed domains and less accurate at reconstructing the landscapes of natural domains (Fig. 3F). This remained true for any number of principal components and even when designed proteins were excluded from the initial PCA (Fig. S10). This indicates that the de novo design protocols examined here lead to structures with “typical” amino acid environments that can be accurately described by only five principal components, and that these proteins generally lack the more specialized environments found in natural domains. Indeed, wild-type amino acids in natural domains tend to be more stable than the fit from the coarse model (Fig. S11). This suggests the remaining components capture additional biophysical effects that contribute to the compatibility between wild-type amino acids and their environments.

Three example proteins shown in Fig. 3G illustrate how the coarse five-component model captures (or fails to capture) protein stability landscapes. At one extreme, the stability landscape of the designed protein r11_692_TrROS (from trRosetta hallucination) is accurately approximated by the coarse model (average per-residue MAE 0.13 kcal/mol). In contrast, the two natural domains (an SH3 domain (1QP2) and a unique NifT/FixU barrel domain (2JN4); Fig. S12) contain many sites with unique properties that are not accurately represented by the model (average per-residue MAE of 0.34 kcal/mol and 0.31 kcal/mol for the SH3 domain and β-barrel domains respectively). Seven of these sites are highlighted in Fig. 3H. Each stability landscape contains sharp differences between closely related amino acids that are not captured by the coarse model, such as V versus L at V8 and Q19 in 1QP2, and Q versus E at Q19 in 1QP2, M28 in 2JN4, and T60 in 2JN4. These unusual patterns are unlikely to be experimental artifacts because the patterns are consistent between independent experiments with trypsin and chymotrypsin and the same patterns are seen in both our K_50_ and ΔG analysis (Fig. 3H). Our massive dataset enabled us to identify the global trends in stability landscapes as well as specific cases that depart from these trends. These unusual cases with large reconstruction errors may provide the opportunity to study how protein flexibility and/or rare side chain interactions contribute to folding stability. These unusual sites will also serve as stringent test cases for models of protein stability.

### Quantifying thermodynamic coupling for hundreds of amino acid pairs

Next, we examined how side chain interaction between amino acid pairs affects folding stability. We constructed comprehensive substitutions (20 x 20 amino acids) of 595 amino acid pairs from 208 natural domains and designs in our ΔG dataset (Dataset #2) and measured stability for all sequences by cDNA display proteolysis. We selected pairs that were suggested to form energetically important hydrogen bonds in our mutational scanning data as well as other pairs forming close contacts (Fig. 4A; Methods). To quantify the interactions between side chains, we constructed an additive model for each amino acid pair with 40 coefficients that capture the independent stability contributions of each amino acid in each position. The deviations from these models quantify the “thermodynamic coupling” between specific amino acids. Among our curated set of wild-type pairs, thermodynamic couplings were typically 0.5-1.0 kcal/mol in magnitude, with the largest couplings stronger than 2 kcal/mol (Fig. 4B). Among all sequences tested (wild-type or mutant pairs), pairs with opposite charges and cysteine pairs tended to have positive (favorable) couplings, whereas pairs with the same charge and acidic-aromatic/aliphatic amino acid pairs tended to have negative couplings (Fig. 4C). These couplings are lower than our observed wild-type couplings because the side chain orientations and environment surrounding wild-type pairs will typically be optimized for that pair. Nonetheless, our data recapitulate expected patterns of side chain interactions, provide a wealth of data for training machine learning models, and identify a wide range of noteworthy interactions for further study.

Several notable pairs are highlighted in Fig. 4D to F. In an OB-domain from *Shewanella oneidensis*, we found strong thermodynamic coupling between two unrelated pairs of amino acids: the wild-type Tyr-Glu pair and a mutant Lys-Trp pair that may form a cation-π interaction (thermodynamic couplings of 1.6±0.2 and 1.4±0.2 kcal/mol respectively; mean±std from calculating the coupling using bootstrap resampling of the ~400 amino acid combinations; Fig. 4D, S13A). In the Alpha-spectrin SH3 domain, our comprehensive double mutant scanning of Y10 and Y52 uncovered the highly stable, tightly coupled double mutant Y10H/Y52K (coupling of 2.5±0.4 kcal/mol for His-Lys versus 1.0±0.2 kcal/mol for the wild-type pair) (Fig. 4E, S13B). AlphaFold modeling predicts that this double mutant introduces a new hydrogen bonding network to replace the original Tyr-Tyr interaction. We also identified an unexpected thermodynamic coupling between an amino acid pair lacking a direct side chain interaction. In the SH3 domain of Myo3, mutations at K24 are destabilizing even though the side chain makes no clear interactions. To investigate interactions of K24, we quantified thermodynamic couplings to nearby Y9 (0.0±0.1 kcal/mol) and D10 (1.0±0.2 kcal/mol) (Fig. 4F and Fig. S13C). The surprising K24-D10 coupling - between two side chains that appear not to interact - highlights the difficulty of inferring energetic interactions from structural data alone, and suggests a possible longer-ranged ionic interaction.

We also investigated thermodynamic couplings within 36 different three-residue networks. For each triplet, we comprehensively measured stability for all possible single and double substitutions in both the wild-type background and in the background where the third amino acid was replaced by alanine (400 mutants x 3 pairs x 2 backgrounds = ~2,400 mutants in total for each triplet). As before, we modeled each set of 400 mutants (i.e. one residue pair in one background) using 40 single-amino acid coefficients (we did not globally model all 2,400 mutants together). One notable triplet is found in the J domain of HSJ1a, where R60 and D64 both interact with the hydroxyl group on Y3 (Fig. 4G left). We observe strong couplings (> 1.5 kcal/mol) between each pair of two out of the three amino acids. However, when any of the three amino acids is mutated to alanine, the coupling between the remaining two amino acids becomes much weaker (< 0.5 kcal/mol, Fig. 4G middle and right, Fig. S13D). These results reveal a strong third-order thermodynamic coupling: the interaction between two amino acids is mediated by a third amino acid.

This strong three-way coupling is especially noteworthy because the interactions do not appear in the deposited NMR structural ensemble (2LGW; Fig S14A and B). The interaction network shown in Fig. 4G comes from the AlphaFold predicted structure for our wild-type sequence taken from the J domain of human HSJ1a. This network reproduces interactions seen in other J-domain crystal structures from *C. elegans* (2OCH) and *P. falciparum* (6RZY). However, in the deposited NMR ensemble for 2LGW, the backbone near Y5 (Y3 in our numbering) always positions that residue away from the helix containing R62 and D66, making the interaction network impossible. The strong couplings we identify support the AlphaFold model and suggest the deposited ensemble is missing conserved interactions that form in HSJ1a and other J domain proteins. This example illustrates how large-scale folding stability measurements can reveal the thermodynamic effects of a critical interaction even when that interaction is missing in the deposited NMR structure. Notably, AlphaFold itself does not always predict this network either: when we include disordered linkers from the NMR construct or used for cDNA proteolysis, AlphaFold also predicts alternative structures lacking the interaction network (Fig. S14D and E).

The scale of our cDNA display proteolysis experiments makes it straightforward to characterize unique cases like these, and again these cases will serve as stringent tests for models of folding stability. Strong third-order couplings like this example also present a special challenge for computational models that calculate stabilities by summing interaction energies between pairs of residues using a single reference structure. Deep learning models that implicitly represent entire conformational landscapes (*42*) may be more promising, but training these models using large-scale thermodynamic measurements will be essential to achieve their potential.

### Natural sequences systematically deviate from their highest stability variants

How does selection for stability influence protein sequence evolution in concert with other evolutionary mechanisms? It is well known that proteins contain specific functional residues that are commonly deleterious to stability (*68, 69*). However, the challenge of measuring stability has made it difficult to experimentally distinguish selection for stability from other selective pressures on a global level (*70–72*). To examine the strength of selection for stability, we created a simple classification model to predict the wild-type amino acid at any site in a natural protein based on the folding stabilities of all substitution variants at that site (excluding Cys) (Fig. 5A). The model contains two parts: (1) a shared weight function that converts absolute stabilities of protein variants into relative probabilities of those sequences, and (2) amino-acid specific offsets that shift amino acid probabilities by a constant amount at all sites. We fit the shared weight function parameters (a flexible monotonically increasing function) and the offsets together using absolute stability data for wild-type sequences and substitution variants at 4,718 sites in 80 non-redundant natural proteins (85,004 ΔG measurements in all, Fig. 5A). Our simple model fits the data well by three criteria: (1) it correctly produces the overall frequencies of the 19 (non-Cys) amino acids (Fig. 5B), (2) the output amino acid probabilities are correctly calibrated across the full range of probability (Fig. S15), and (3) the model performs similarly well on the training set and on a held-out testing set consisting of 621 sites in 11 domains with no similarity to the training set (Fig. 5E).

The model parameters reveal the strength of selection for stability across this heterogeneous set of domains from many organisms. Within the main range of our data (folding stabilities from 1.5 to 4 kcal/mol), amino acid probabilities increase approximately linearly with increased stability, with a 1 kcal/mol stability difference between protein variants indicating a ~5.7-fold difference in sequence likelihood (Fig. 5C). The global offsets to each amino acid’s probability (Fig. 5D) are different from the empirical amino acid frequencies (Fig. 5B) and indicate the probability of each amino acid under conditions where all sequence variants are equally stable. The offsets span a 23-fold range: the most likely amino acid (Glu) is 23-fold more likely to occur (2^1.5^/2^-3.0^) than the least likely amino acid (Trp) under the conditions that sequence variants containing these amino acids at the same site are equally stable (Fig. 5D). This probability difference corresponds to a stability difference of ~1.8 kcal/mol (Fig. 5C); i.e. Trp and Glu would be equally likely at a site if the Trp variant were 1.8 kcal/mol more stable than the Glu variant. Overall, the most likely amino acids are the charged amino acids Glu, Asp, and Lys, suggesting selection for solubility, whereas the least likely amino acids are the nonpolar aromatic amino acids Trp, Phe, and Tyr, along with Met. These offsets provide a quantitative “favorability” metric incorporating all non-stability evolutionary influences on amino acid composition, including selection for amino acid synthesis cost (*73, 74*), codon usage (*75, 76*), avoiding oxidation-prone amino acid(s), net charge, and function. These offsets also highlight that biophysical models and protein design methods trained to reproduce native protein sequences will not consistently optimize folding stability; Fig. 5D quantifies how much specific amino acids are over- or underrepresented in small, naturally occurring domains compared to their effects on stability. Notably,these offsets are similar to findings from an independent analysis of global discrepancies between variant effect data and sequence likelihood modeling (*77*)

### Properties of functional residues across diverse domains

Selection for function also causes protein sequences to diverge from the highest stability sequence variants. Previous studies (*70, 71*) have applied this strategy to identify functional sites based on the difference between evolutionary conservation and predicted effects on stability. We expanded this strategy to employ experimental stability measurements and examined the properties of functional sites on a large scale. We identified functional sites in 92 diverse protein domains by comparing each site’s average ΔΔG of substitutions with its normalized GEMME (*78*) score, an evolutionary-based measure of sensitivity to mutations (Fig. 6A, see Methods for the details). High sensitivity generally indicates high evolutionary conservation. Sites where wild-type amino acids are critical for stability (higher average ΔΔG, rightward) tend to be predicted as more sensitive to mutation by GEMME (upward) and vice versa. We defined all sites in the upper left region (where the wild-type amino acid is conserved yet unimportant for stability, 9.3% in total) to be “functional” sites. This classification correctly identifies key binding residues in the chromodomain of HP1 and the SH3 domain of BBC1 (Fig. 6B and C, see Fig S16 for mutational scanning and conservation data on these examples). We found that Gly, Asp, and the bulky amino acids Trp Arg, and Tyr were frequently classified as functional (Fig. 6D). However, like previous studies, our classification method has the notable weakness that any site that is important for folding stability will not be considered functional.

Across all 92 domains, the fraction of functional sites ranged from 0 to ~25% (Fig. 6E). The domains with the highest fraction of functional sites (the Sso7d protein (1JIC) and Ribosomal protein S19 (1QKH)) are both nucleic acid binding proteins, with the functional sites located on the surface primarily at the binding interface (Fig. 6F). To identify buried functional sites, we compared each site’s evolutionary-based sensitivity to non-polar mutations (normalized GEMME score for hydrophobic substitutions) to the average ΔΔG of nonpolar substitutions (Fig. 6G), a more permissive metric. With this approach, most functional sites are still located at the protein surface, but a small fraction are located in the core (Fig. 6H). One example is A64 in the DUF1471 domain of yahO. A64 is highly sensitive to non-polar mutations and buried in the core of the domain, but substitutions to Tyr or Phe increase folding stability (Fig. 6I). This indicates that A64 modulates the function of the domain even without interacting with external partner molecules, perhaps by maintaining the overall protein shape. Similarly, in the N-terminal domain of FK506-binding protein 3, L55 is buried in the core and highly conserved even though substitutions to Ile, Val, or Phe have no effect on stability (Fig. 6J). This domain binds DNA and the other functional residues are mainly located at the binding interface. Although L55 does not directly interact with DNA, substitutions to other hydrophobic amino acids may change the orientations of the surface side chains and prevent proper DNA binding. Notably, chemical shift perturbations in this domain indicate which residues change their magnetic environment in response to DNA binding (Fig. 6J) (*79*). Chemical shift perturbations are found mainly in the functional residues on the protein surface, but L55 experiences a chemical shift perturbation as well, indicating allosteric communication between the functional surface residues and L55. These results highlight unusual cases where buried sites are conserved due to specific functional requirements rather than to maintain overall stability.

### Large-scale stability analysis to characterize unique designs, identify stabilizing mutations, and evaluate design methods

The unique scale of cDNA display proteolysis creates new opportunities for improving protein design. Here, we examine three applications of our method and massive dataset: (1) characterizing the stability determinants of rare, highly polar proteins, (2) identifying stabilizing mutations, and (3) benchmarking the protein design tool PROSS (*80*). The hydrophobic effect is considered the dominant force in protein folding (*1*), and measuring stability for thousands of our previously-designed domains (*29*) by cDNA display proteolysis revealed a general trend of increasing stability with increasing hydrophobicity (Fig. 7A). However, increased hydrophobicity can promote protein aggregation, non-specific interactions, and low expression yield. To study the properties of high stability, low hydrophobicity proteins, we examined hundreds of designed proteins by deep mutational scanning across a wide range of hydrophobicity and stability. Although the mutational scanning patterns for low hydrophobicity proteins were not obviously different from other designs, we identified several designs that possessed exceptionally strong polar interactions (large dots in Fig. 7A). In Fig. 7B, we highlight stabilizing polar networks and a cation-π interaction in these unusual designs (see Fig. S17 for full mutational scanning results). The average ΔΔG for substitutions at these polar sites ranges from −0.20 to −1.33 kcal/mol, corresponding to the top 63 to 1.5%ile for all 3,694 polar sites in 145 designs. Our unusually massive dataset made it possible to identify these rare highly stabilizing interactions. Notably, the second hydrogen bond network in EHEE_rd2_0152 is also found in two other more hydrophobic designs. However, the network is less sensitive to substitution in those designs, highlighting how the overall protein environment mediates the effects of substitutions even on the protein surface (Fig. S18).

We next examined how our approach could be used to identify stabilizing mutations. Predicting and designing stabilizing mutations is a major goal of protein modeling, but prediction accuracy remains low (*22*). In part, this is because stabilizing mutants are rare in current databases (*14, 15*) (outside of reverting a destabilizing mutant), limiting the data available for improving modeling. In contrast, our large-scale approach revealed 2,600 stabilizing mutations, defined as mutations that increase folding stability by at least 1 kcal/mol. The overall fraction of stabilizing mutations was 0.06% to 0.6% for different protein types (Fig. 7C). Stabilizing mutations were enriched at functional sites (23% of the stabilizing mutations from 7.5% of sites classified as functional), but these were still a small fraction of the total. Notably, our set includes 112 examples of stabilizing insertions and deletions which are nearly absent from current databases. In Fig. 7D, we show three examples of different classes of stabilizing mutations found in our dataset with effects ranging from +1.2 to +3.1 kcal/mol (Fig. S19).

Finally, we applied our method to evaluate PROSS (*80*), an automated method for enhancing folding stability within sequence constraints inferred from a multiple sequence alignment. We tested 1,156 PROSS designs for 266 protein domains (a 10-100x increase over previous benchmarking study (*81*)). Unlike previous studies, our mutational scanning data for all 266 wild-type domains enabled us to examine the isolated effect of every individual substitution in every PROSS design. The average increase in stability from PROSS was 0.6±1.0 (mean±std) kcal/mol, and 40% of 727 domains (with wild-type ΔG < 4 kcal/mol) had at least one design with a 1 kcal/mol increase in stability (Fig. 7E). As expected, PROSS avoided mutations at functional positions: only 1.9% of PROSS-designed mutations were found at functional positions compared to 8.7% of sites classified as functional (defined in Fig. 6A). Three examples of domains successfully stabilized by PROSS are shown in Fig. 7F. Although the median number of designed mutations was only 4, more mutations typically led to a larger increase in stability (Fig. S20A), as theorized previously (*22*). Based on our mutational scanning data, the average effect of an individual PROSS mutation was 0.22±0.47 kcal/mol (Green line Fig. S20B). On average, the added stabilization from PROSS is comparable in size to the effect of the best single mutant designed by PROSS, and smaller than the additive effect of the two best designed mutations (Fig. S20C). Evaluating individual mutations recommended by PROSS (or other design tools) by direct comparisons to mutational scanning data provides a novel approach for systematically improving these design methods.

## Discussion

The cDNA display proteolysis method massively expands the scale of folding stability experiments. Still, the method currently has notable limitations. Because we digest proteins under native conditions, our inferred thermodynamic stabilities are only accurate when (1) folding is fully cooperative (no unfolded segments get cleaved without global unfolding (*82*)), (2) folding is at equilibrium during the assay (no kinetic stability or spurious stability due to aggregation), (3)K_50,U_ is accurately inferred (Fig. 1C), and (4) cleavage rates fall within the measurable range of the assay, which currently limits the dynamic range to ~5 kcal/mol (Fig. 1C). Many domains - particularly larger protein structures - will not satisfy these conditions, and issues such as non-cooperativity, kinetic stability, or aggregation are invisible in a single measurement. Combining cDNA display proteolysis with chemical denaturation (pulse proteolysis, (*37*)) may overcome these obstacles and enable mega-scale analysis of less cooperative and/or higher stability proteins, while also avoiding the need to infer K_50,U_. Advances in DNA synthesis (including methods like DropSynth (*83, 84*)) will also make it possible to expand cDNA display proteolysis to analyze diverse libraries of larger domains. Lastly, multiplexed measurements and automated data processing always have the potential to introduce inaccuracies, although we worked to exclude unreliable data. For notable individual results, examining the raw data can be helpful, and we included all data and code to regenerate all fits.

Despite these limitations, the unique scale of cDNA display proteolysis opens completely new possibilities for studying protein stability. By comprehensively measuring single mutants across nearly all small structures in the Protein Data Bank, we quantified several global trends: trends in amino acid fitness at different sites, trends in the effects of single and double mutants, and trends in how stability influences sequence evolution. Along with these global trends, our large-scale analysis also uncovered hundreds of exceptional cases that would be challenging to identify by smaller-scale methods. These include mutations with extreme effects, sites with unusual stability landscapes, and pair interactions with unusually strong thermodynamic couplings. The strong thermodynamic couplings we identified in the J domain of human HSJ1a (Fig. 4G) - missing in the deposited NMR structure - highlight how large-scale stability assays can complement other methods for revealing structural details in solution. The 2,400 double mutants examined in that domain made up only 0.3% of the experimental library. Beyond studying the origins of stability, cDNA display proteolysis will have a range of other applications, including assaying designed proteins on a massive scale to systematically improve design methods (*29, 43, 85*), identifying folded domains in metagenomic sequences, and dissecting the relationships between folding stability and function (*41*).

Achieving an accurate, quantitative understanding of protein stability and its sequence dependence has been a central goal in biophysics for decades. We envision millions of cDNA display proteolysis measurements forming the foundation for a new generation of deep learning models predicting absolute folding stabilities and effects of mutations. Breakthroughs in deep learning-powered structure prediction have proven the power of these models in protein science, but collecting sufficient thermodynamic data has always been a major obstacle. Due to the scale and efficiency of cDNA display proteolysis, the main limit to measuring stability for millions of small domains is the cost of DNA synthesis (*86–88*) and sequencing (*89, 90*) - both of which are rapidly decreasing (*91–94*). With the flexibility of DNA oligo synthesis, cDNA display proteolysis can assay massive mutational libraries (as shown here) as well as massive libraries of unrelated sequences and structures, which will add essential diversity in training datasets. The size and diversity of protein sequence space creates enormous challenges for biology and protein design. The cDNA display proteolysis method offers a powerful approach to map folding stability across this space on an unprecedented scale.

## Acknowledgments

We thank Epsilon Molecular Engineering (EME) Corp for providing us with cnvK linker for cDNA display, Rush University and Genome Research Core at University of Illinois Chicago for performing next-generation sequencing, and David Minh, Timothy Whitehead, Kresten Lindorff-Larsen, David M. McCandlish, Jack Maguire, John Chodera, Parisa Hosseinzadeh, and the members of the Rocklin lab for discussions and comments on the manuscript.

## Author contributions

KT designed and performed all experiments, and analyzed the data with help from GJR. JD designed and analyzed stabilities of hallucination-based proteins with help from SO. JC designed ββαα proteins using Rosetta with help from GJR. EL computed GEMME scores with help from YBM, and also assisted with interpretation of GEMME. JW generated the PROSS designs. NMM provided assistance with mathematical derivation and review of enzyme kinetic interpretation. GJR and KT conceived the project. GJR supervised the project and acquired funding. KT and GJR wrote and revised the manuscript, with input from all authors.

## Data and materials availability

All data and codes are available in the main text, the supplementary materials, or available for download at https://doi.org/10.5281/zenodo.7401275 or https://github.com/Rocklin-Lab/cdna-display-proteolysis-pipeline

## Supplementary Materials

### Materials and Methods

#### DNA oligo library construction

All sequences were reverse-translated and codon-optimized using DNAworks2.0 (*95*). Sequences were optimized using *E. coli* codon frequencies because we used an *in vitro* translation kit derived from *E. coli*. Oligo libraries encoding amino acid sequences of Library 1 were purchased from Agilent Technologies. Oligo libraries for Libraries 2-4 were purchased from Twist Bioscience.

##### Library 1

We selected ~250 designed proteins and ~50 natural proteins that are shorter than 45 amino acids. Then, we created amino acid sequences for deep mutational scanning followed by padding by Gly, Ala, Ser amino acids so that all sequences have 44 amino acids. The total number of sequences is ~244,000 sequences.

##### Library 2

We selected ~350 natural proteins that have PDB structures that are in a monomer state and have 72 or less amino acids after removing N and C-terminal linkers. Then, we created amino acid sequences for deep mutational scanning followed by padding by Gly, Ala, Ser amino acids so that all sequences have 72 amino acids. The total number of sequences is ~650,000 sequences. This library also includes scramble sequences to construct unfolded state model.

##### Library 3

We selected ~150 designed proteins and created amino acid sequences for deep mutational scanning of the proteins. We also included comprehensive deletion and Gly/Ala insertion of all wild-type proteins inlcuded in Library1 and Libary2. Additionally, amino acid sequences for comprehensive double mutant analysis on polar amino acid pairs were also included.

##### Library 4

Amino acid sequences for exhaustive double mutant analysis on amino acid pairs located in close proximity were included. We also include overlapped sequences to calibrate effective protease concentration and to check consistency between libraries.

#### EEHH design method

EEHH protein design was performed in three steps: (1) backbone construction, (2) sequence design, (3) selection of designs for deep mutational analysis. Backbone construction (the de novo creation of a compact, three-dimensional backbone with a pre-specified secondary structure) was performed using a blueprint-based approach described previously (*96, 97*). All blueprints are included as Blueprints_for_EEHH.zip in Supplementary Materials.

#### Hallucination design method

We used a TrRosetta hallucination protocol described previously in (*42, 67*) and available at https://github.com/gjoni/trDesign/tree/master/02-GD to unconditionally generate protein backbones and sequences with lengths ranging from 46 to 69 amino acids by maximizing the Kullback–Leibler divergence between the predicted and background distance/angle distributions. Predicted distograms and anglegrams were used to obtain 3D structures of these models as described in the TrRosetta paper (*98*). We selected the best designs according to the predicted distogram and 3D structure match.

#### DNA and mRNA preparation for cDNA display proteolysis method

Oligo libraries were amplified by PCR using KOD PCR Master Mix (TOYOBO) to add T7 promoter, PA tag to an N-terminal, and His tag to an C-terminal of the proteins. The number of cycles was chosen based on a test qPCR run to avoid overamplification using SsoAdvanced Universal SYBR Green Supermix (BIORAD). The PCR product was gel extracted to isolate the expected length product. Then we used T7-Scribe Standard RNA IVT Kit (CELLSCRIPT) to synthesize mRNA using the DNA fragment as a template.

#### Preparation of protein-cDNA complex

We basically follow the protocol described in the previous literature (*47, 99*) with some modifications.

##### Photocross-linking between mRNA and the puromycin linker

We prepared the photocrosslinking reaction solution including 200 mM NaCl, 40 mM Tris-HCl (pH 7.5), 20 μM cnvK linker (EME corporation), 20 μM mRNA. The solution was incubated at 95°C for 5 min, then slowly cooled down to 45°C (0.1°C / 1 second) using a thermal cycler. Then the solution including the duplex was irradiated with UV light at 365 nm using a 6W Handheld lamp (Thermofisher).

##### In vitro translation and reverse transcription

We prepared PUREfrex 2.0 (GeneFrontier) translation system with mRNA-cnvK linker duplex and RiboLock RNase Inhibitor (Thermofisher) and incubate the sample at 37°C for 2 hrs. After the incubation, 100 mM EDTA was added to the sample to dissociate ribosomes. Then, an equal amount of binding/washing buffer (30 mM Tris pH 7.5, 500 mM NaCl, 0.05% Tween 20) was added. The solution was added to Dynabeads MyOne Streptavidin C1 (Thermofisher) to pull down the protein-mRNA complex and incubated at room temperature for 20 min. Then, the beads were washed by binding/washing buffer once and rinsed twice by TBS (10 mM Tris-HCl pH7.5, 100 mM NaCl), and we added reverse transcription solution (PrimeScript RT Reagent Kit; Takara) onto the beads with protein mRNA complex, and incubated the beads at 37°C for 30 mins.

##### Purification of protein-cDNA complex

After the reverse transcription, the protein-cDNA complex was eluted by binding/washing buffer with RNase T1 (Thermofisher). The eluent was added His Mag Sepharose Ni (Cytiva) and incubated at room temperature for 30 min. Then the complex was eluted by binding/washing buffer with 400 mM imidazole then the eluent was buffer-exchaged by Zeba Spin Desalting Column (Thermofisher). Then the complex was snap-frozen by liquid nitrogen and stored at −80°C until the following protease assay.

##### Protease assay on protein-cDNA complex

We prepared 40 μL of 11 protease three-fold dilution series from 25 μM for replicate1 and 43.3 (= 25 x 3^0.5^) μM for replicate2, then added them to 12 of 20 μL the protein-cDNA complex. After 5 min protease digestion in room temperature, we added 200 μL chilled 2% BSA in PBS to quench the reaction, then the solution was added to 10 μL Dynabeads Protein G (Thermofisher) with anti-PA tag antibody (Wako; Clone number: NZ-1; 1μg antibody per 30 μL beads), and incubated at 4°C for 1 hr. Then the beads were washed by washing buffer (PBS including 800 mM NaCl and 1% Triton) three times and rinsed by PBS three times, then the complex was eluted with 50 μL PBS including 250 μg/mL PA peptide (Wako) and 200 μg/mL BSA (Thermofisher).

#### qPCR analysis of cDNA display proteolysis results on individual proteins (for Fig. S1)

The cDNA amount for each specific sequence in the eluents was quantified by qPCR using SsoAdvanced Universal SYBR Green Supermix and specific primers for each sequence. The qPCR was performed using CFX96 Touch Real-Time PCR Detection System (BIORAD), and the qPCR cycles were determined by the CFX Maestro Software (BIORAD).

#### Next-generation sequencing sample preparation

For DNA library analysis, one-half volume (25 μL) of the eluted cDNA of the complex was amplified by PCR using SsoAdvanced Universal SYBR Green Supermix (BioRad) to add P5 and P7 NGS adapter sequence. The number of cycles was chosen based on a test qPCR run using the same PCR reagents to avoid overamplification. The DNA fragment length and concentration were confirmed by 4200 TapeStation System (Agilent), then the samples were analyzed by NovaSeq 6000 System (Illumina).

#### Processing of next-generation sequencing data

Each library in a sequencing run was identified via a unique 6 or 8 bp barcode. Following sequencing, reads were paired using the PEAR program (*100*) then the adapter sequences were moved by Cutadapt (*101*). Reads were considered counts for a sequence if the read perfectly matched the ordered sequences at the nucleotide level.

#### Overall strategy for inferring K_50_ and ΔG from sequencing data

We used Bayesian inference to infer K_50_ and ΔG values for all sequences in our library. This analysis uses two main models. The first model is called the “K_50_ model” and infers each sequence’s K_50_ values based on the sequencing count data. The second model is called the “unfolded state model” and predicts each sequence’s unfolded state K_50_ value (K_50,U_) based on its sequence. Both models are implemented in Python 3.9 using the Numpyro package (*102*) version 0.80. Here, we first describe the structure of each model, and then we describe the practical process of fitting the parameters of each model. Our scripts to reproduce the complete fitting process are provided in the Supplementary Materials.

#### Structure of the K_50_ model to infer K_50_ values from next-generation sequencing data

We modeled our selection results using the single turnover kinetics model described in Fig. 1B. We chose this model because we expect that the total concentration of protein-cDNA complex is low compared to the amount of added enzyme and because the model captures the saturation behavior observed by qPCR at high enzyme concentration (Fig. S1). Instead of attempting to capture the microscopic complexity of our system (millions of different substrates and potential inhibitors), the purpose of the model is to treat each substrate in a consistent, simplified manner and infer reasonable parameters.

Our model makes two main assumptions. First, we assume that each sequence is cleaved independently, with no competition or product inhibition. As described by Fig. 1 eqs. 2 and 3, cleavage is described by four parameters: enzyme concentration (*E*), time (*t*), and the kinetic parameters *K_50_* and *k*_max_. All experiments used a fixed five minute reaction time. Based on qPCR analysis of individual sequences (Fig. S1), we fixed the quantity *k_max_* * *t* at 10^0.65^ for all sequences. Each sequence’s unique stability is defined by the *K_50_* parameter that represents the enzyme concentration producing the half maximal cleavage rate (Fig. 1 eq. 3). Our second main assumption is that we can interpret our *K_50_* values as representing the dissociation constants (*K_D_*) between each protein sequence and the enzyme (*K_50_* ≈ *K_D_*, Fig. 1 eq. 6). From this assumption, we can determine the folding stability of each sequence (ΔG) based on the relationship between the observed *K_50_* value and theoretical *K_50_* values for the fully folded and fully unfolded states (*K_50,F_* and *K_50,U_*, Fig. 1 eqs. 5-7). Although we can directly fit *K_50_* values without making any assumptions about the microscopic basis for *K_50_* (see Supplementary Text for the detail), assuming that *K_50_* ≈ *K_D_* aids our interpretation and enables us to directly fit ΔG values to our data using the *Coupled* approach described below.

To fit our model to our sequencing counts data, we first assume that the cDNA display process produces an unknown initial distribution of full-length protein-cDNA complexes (the *cDNA_0_* distribution). The distribution of sequences at enzyme concentration *E* (the *cDNA_E_* distribution) is the product of the initial sequence distribution *cDNA_0_* and the surviving fraction of each sequence according to Fig. 1 eqs. 2 and 3, after re-normalizing the total surviving fraction of all sequences to 1.

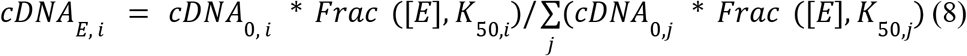

Finally, we assume that our deep sequencing counts result from *n_sel_* independent selections from the *cDNA_E_* distribution, where *n_sel_* is the number of sequencing reads that exactly matched our specified DNA sequences.

We apply the K_50_ model in two different ways based on whether K_50_ values for trypsin and chymotrypsin are *Independent* or *Coupled*. The “Independent” procedure is used in Steps 1, 2 and 5 in the section “Procedure for fitting all data”. In the independent procedure, the inputs to the model are the sequencing counts data from experiments with one protease, the enzyme concentrations, the reaction time, and the *k_max_* constant. We fit the model by sampling two parameters per sequence from normal prior distributions: (1) *K_50_*, and (2) the initial fraction of each sequence in the *cDNA_0_* distribution. The “Coupled” procedure is used in Step 5 in the section “Procedure for fitting all data”. In the coupled procedure, the inputs to the model are the sequencing counts data from experiments with both proteases, the enzyme concentrations, the reaction time, the *k_max_* constant, the *K_50,F_* constants representing the universal K_50_ value for sequences in the folded state (one for each protease), and the predicted *K_50,U_* values for all sequences for both proteases from the unfolded state model. We then assume that each sequence has a specific ΔG value that is shared across both proteases. We use this shared ΔG value along with *K_50,F_* and *K_50,U_* (for each protease) to determine *K_50_* for each protease according to Fig. 1 Eqs. 5 and 7. Finally, we fit the coupled model by sampling two parameters per sequence from normal prior distributions: (1) ΔG, and (2) the initial fractions of each sequence in *cDNA_0_*.

Full results from both the independent and coupled fitting procedure are provided in K50_dG_Dataset1_Dataset2.csv and K50_Dataset3.csv. For our stability parameters (protease-specific *K_50_* in the independent procedure and ΔG in the coupled procedure) we report the median of the posterior distribution as well as the upper and lower limits of the 95% confidence interval (the 2.5%ile and 97.5%ile values of the posterior distribution). We also used the protease-specific *K_50_* values from the independent procedure to compute protease-specific ΔG values. We do this using the same *K_50,F_* and *K_50,U_* values used in the coupled procedure according to Fig. 1 Eqs. 5 and 7. These protease-specific ΔG estimates are also reported in K50_dG_Dataset1_Dataset2.csv and are only used to examine the consistency between different proteases (e.g. Fig. 1F and Fig. 2D). In some cases, the independently fit K_50_ values can lead to impossible values for ΔG. This can occur if *K_50_* is higher than *K_50,F_* (observed cleavage is slower than our limit for cleavage in the folded state) or if *K_50_* is lower than *K_50,U_* (observed cleavage is faster than predicted cleavage in the unfolded state). If the median protease-specific *K_50_* or the confidence interval limits for a particular sequence lead to impossible ΔG values for that sequence, we report dummy values for the corresponding protease-specific ΔG estimates.

#### Structure of the unfolded state model to infer unfolded K_50_ (K_50,U_) from scrambled sequence data

Our unfolded state model is similar to the model employed previously (*29*) with two notable differences. First, instead of assuming that all scrambled sequences are fully unfolded, we assume that each scrambled sequence has its own unknown folding stability, with a prior distribution biased toward low stability (normal prior centered at ΔG = −1, sigma = 4). Second, instead of fitting an unfolded state model for each protease independently, we assume that each scrambled sequence’s stability (ΔG) is common across both proteases, and fit the models for each protease together. As a result, the majority of scrambled sequences are modeled as completely unfolded (Fig. S2C), but some scrambled sequences are modeled as stable when that interpretation is consistent with both the trypsin and chymotrypsin data.

Our unfolded state has three parts: (1) a position specific scoring matrix (PSSM) that describes how the amino acid sequence in a 9-mer window (the P5 to P4’ positions in protease nomenclature) determine the cleavage rate at the P1 position, (2) a local response function describing the saturation of the cleavage rate for a single P1 position, (3) a global response function that determines K_50,U_ based on the sum of the cleavage rates at all possible P1 positions in the full sequence.

To fit the PSSM, we assumed an identical normal prior distribution of scores at all positions, with several exceptions. Due to known critical importance of the P1 position, we used a wider prior distribution of scores for all amino acids in the P1 position for both proteases. We also used wider prior distributions at all positions (P5-P4’) for the amino acids Asp, Glu, and Pro, due to the established large effects of these amino acids on cutting rates.

For the local response function to saturation of the cleavage rate at P1 site *k*, we used a logistic function:

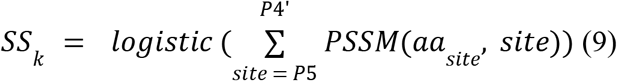

where *SS_k_* (site saturation) is the saturation of the cutting rate at site P1=*k, aa_site_* is the amino acid identity at *site*, and *logistic* is the logistic function f(x) = 1 / (1+e^x^). We fit the 21 (20 amino acids + ‘X’ representing empty sites) x 9 =189 elements of the PSSM for each protease.

For the global response function (determining K_50,U_ based on the sum of *SS_k_* across the full protein sequence), we use a sum of logistic functions with 10 different activation thresholds.

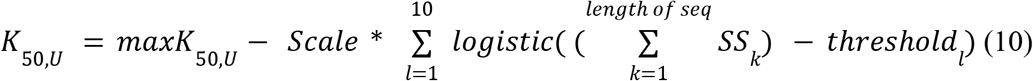

where *maxK_50,U_* is the highest possible *K_50,U_* value (*K_50,U_* assuming no cut sites), *Scale* is the range of possible *K_50,U_* values, and *threshold_l_* is the value of the *l*^th^ activation threshold for the global response function. All K_50_ values (including *maxK_50,U_*) are in log_10_ molar units.

The key parameters of the unfolded state model (for a single protease) are the 21 x 9 =189 elements of the *PSSM*, the *maxK_50,U_*, the *scale*, and the 10 *threshold* values. These parameters determine *K_50,U_* for each sequence by Eqs. 9 and 10. In addition to these parameters, we also sample the ΔG values for each scrambled sequence during fitting. These sampled parameters (as well as the universal *K_50,F_* value for all sequences) are sufficient to determine a theoretical *K_50_* value for each scrambled sequence by re-writing Fig. 1 Eq. 6:

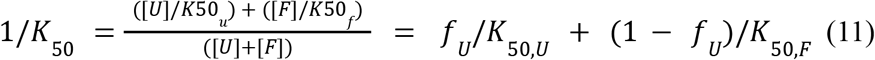

where *f_U_* is the fraction of unfolded molecules:

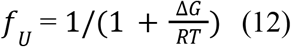

The input data for the model are the observed *K_50_* values for all scrambled sequences. The parameters of the model are fit by assuming that all observed *K_50_* values should agree (with small, normally distributed errors) with the theoretical *K_50_* values determined by the model parameters. After fitting the model, we used the median of the posterior distributions of *PSSM, maxK_50,U_, scale*, and the 10 *threshold* parameters as the final model parameters. We used these final model parameters to calculate *K_50,U_* for all sequences in our experiments without considering any uncertainty from the model posterior distribution.

#### Procedure for fitting all data

##### Step 1: Estimation of ‘effective’ protease concentrations for each library

We employed four DNA oligonucleotide libraries for this study. Although we tried to minimize the difference between assay conditions, we also fit “effective” protease concentrations to our data in order to minimize batch-to-batch differences. We used the K_50_ model to perform this fitting and fit protease concentrations for trypsin and chymotrypsin entirely independently. The main assumption of this fitting is that each sequence should have the same K_50_ when assayed in different libraries. By enforcing that each sequence had a single K_50_ value regardless of what library it appears in, we calibrated the protease concentrations in each library against each other. Although we did not use universal control sequences in all four libraries, each library contained 1000 to 2000 sequences that overlapped at least one other library in a fully connected graph. Specifically, the library pairs 1+4, 2+4, 3+4, 1+2, and 2+3 each included 1,000 to 2,000 overlapping sequences.

The overall model included 96 experimental conditions (12 protease concentrations per replicate x 2 replicates x 4 libraries; one of the 12 protease concentrations was the fixed “no protease” starting condition). However, each sequence was only present in 48 of the 96 conditions because any individual sequence was only present in two out of the four libraries. The inputs to fit the model were the sequencing counts data, the reaction time (*t*), and the *k_max_* constant. Additionally, to set the overall scale of the protease concentration series, we fixed the effective protease concentrations for Library 4 at the expected protease concentrations (i.e. three-fold serial dilutions of 25 μM protease (Replicate 1) or 43.3 μM protease (Replicate 2)). We also fixed all of the starting samples at zero protease. Using these model inputs, we sampled the K_50_ values (one per sequence), the remaining 66 protease concentrations, and the initial sequence distributions cDNA_0_ (a separate cDNA_0_ was used for each of the 8 replicates). Normal priors (with lower/upper boundaries for some parameters) covering the range of experimentally relevant values were used for the model parameters. Sampling was performed using the No U-Turn Sampler (NUTS) in Numpyro with 50 steps of equilibration and 25 steps of production. We used the medians of the protease concentrations from our 25 posterior samples as our final calibrated protease concentrations for all further analysis (discarding the uncertainties).

##### Step 2: Estimation of K_50_ values of scramble sequences

To train the unfolded state model, we need to determine K_50_ values for our scramble sequences, which were included in Library 2. We used the Independent K_50_ model for this step. The input data were the sequencing counts data from two replicates (i.e. 12 protease concentrations x 2 replicates = 24 data points per sequence), the reaction time (*t*), the *k_max_* constant, and the effective protease concentrations obtained in Step 1. We sampled the initial sequence distribution cDNA_0_ (a separate cDNA_0_ for each replicate) and K_50_ for all sequences included in Library 2. Normal priors (with lower/upper boundaries for some parameters) covering the range of experimentally relevant values were used for the model parameters. Sampling was performed using the No U-Turn Sampler (NUTS) in Numpyro with 100 steps of equilibration and 50 steps of production.

##### Step 3: Construction of unfolded state model

We trained the unfolded state model for predicting K_50,U_ using K_50_ values obtained in Step 2. The input sequences were scrambled sequences of wild-type domains selected for deep mutational screening. In addition to our set of exactly scrambled sequences (matching the wild-type amino acid composition 100%), we also included scrambled sequences containing 50%, 60%, 70%, 80%, and 90% of the number of hydrophobic amino acids in the original wild-type sequences. These sequences helped ensure the large majority of our scrambled pool was fully unfolded. Additionally, because all sequences in our experiments are padded with G/S/A linkers up to a constant length, we generated scrambled sequences using two different padding procedures. In the first approach, we designed scrambled sequences that matched the original wild-type length and were padded with G/S/A up to 72 amino acids. In the second approach, we designed 72 amino acid-length scrambles approximately matching the composition of an original wild-type domain, regardless of the length of that wild-type. These scrambled sequences required no additional padding. After measuring K_50_ for all scrambles, we only used sequences with a 95% confidence interval smaller than 0.5 log_10_ molar units for model training for model fitting (64,238 sequences in total, see Fig. S3). In addition to the exact experimental sequences, we also augmented the training dataset with dummy sequences where GS linkers were replaced by the blank ‘X’ amino acid.

The inputs for the model are amino acid sequences created as described above, and their observed K_50_ for trypsin and chymotrypsin obtained in Step 2. The parameters of the model are fit by assuming that all observed *K_50_* values should agree (with small, normally distributed errors) with the theoretical *K_50_* values. In this model, we sampled the 21 x 9 =189 elements of the *PSSM*, the *site bias*, the *maxK_50,U_*, the *scale*, and the 10 *threshold* values. These parameters determine *K_50,U_* for each sequence by Eqs. 9 and 10. In addition to these parameters, we also sample the ΔG values for each scrambled sequence during fitting.

Normal priors (with lower/upper boundaries for some parameters) covering the range of experimentally relevant values were used for the model parameters. Using NUTS model, we sampled the parameters described above, then reported the median of the 100 posteriors after removing the initial 400 steps. In Step 4, we used these final model parameters to calculate *K_50,U_* for all sequences in our experiments without considering any uncertainty from the model posterior distribution.

##### Step 4: Prediction of unfolded K_50_ values (K_50,U_) across the full dataset

Using the final model parameters obtained in Step 3, we predicted K_50,U_ values for each amino acid sequence in the libraries without considering any uncertainty. Additionally, since the model was constructed to predict unfolded K_50_ for sequences with 86 amino acids, we added a Gly linker ‘GGG’ to both ends, followed by padding by ‘X’ up to 86 amino acids.

##### Step 5: Estimation of K_50_ values and calculation of ΔG for trypsin and chymotrypsin

We applied the *Coupled* K_50_ model to each of the four libraries separately. The inputs to the model are the sequencing count data from trypsin and chymotrypsin experiments (i.e. 12 protease concentrations x 2 replicates x 2 proteases = 48 data points per sequence), the effective protease concentrations obtained in Step 1, the reaction time, the *k_max_* constant (t*k_max_ = 10^0^.^65^ based on qPCR analysis; see Fig. S1), the *K_50,F_* constants (3 for trypsin, 2 for chymotrypsin; determined based on the dynamic range of proteolysis experiment; see Fig. S5), and the *K_50,U_* values predicted by the unfolded model in Step 4. Using the inputs, we sampled ΔG shared between trypsin and chymotrypsin, and initial sequence distribution cDNA_0_ for each protease for each replicate (although our experiments utilized the same batch of the cDNA-protein complex for two replicates).

Normal priors (with lower/upper boundaries for some parameters) covering the range of experimentally relevant values were used for the model parameters. Using NUTS in Numpyro module, we sampled the posteriors of shared ΔG along with other parameters, then obtained the median of the 50 posterior samples after removing the initial 100 steps. Full results from both the independent and coupled fitting procedure are provided in K50_dG_Dataset1_Dataset2.csv and K50_Dataset3.csv. For our stability parameters (protease-specific K_50_ in the independent procedure and ΔG in the coupled procedure) we report the median of the posterior distribution as well as the upper and lower limits of the 95% confidence interval (the 2.5%ile and 97.5%ile values of the posterior distribution).

We also applied the *Independent* K_50_ model to each of the four libraries separately. The inputs to the model are the sequencing count data (i.e. 12 protease concentrations x 2 replicates = 24 data points per sequence), the effective protease concentrations obtained in Step 1, the reaction time, the *k_max_* constant (t*k_max_ = 10^0^.^65^ based on qPCR analysis; see Fig. S1). Using the inputs, we sampled K_50_ for each protease, and initial sequence distribution cDNA0 for each protease for each replicate (although we utilized the same batch of the cDNA-protein complex for two replicates).

Normal priors (with lower/upper boundaries for some parameters) covering the range of experimentally relevant values were used for the model parameters. Using NUTS in Numpyro module, we sampled the posteriors of *K_50_* for trypsin and *K_50_* for chymotrypsin along with other parameters, then obtained the median of the 50 posterior samples after removing the initial 100 steps.

Then, we computed protease-specific ΔG values using the protease-specific K_50_ values from the *Independent* model. We do this using the same *K_50,F_* and *K_50,U_* values used in the coupled procedure according to Fig. 1 Eqs. 5 and 7. These protease-specific ΔG estimates are also reported in K50_dG_Dataset1_Dataset2.csv and K50_Dataset3.csv, and are only used to examine the consistency between different proteases (e.g. Fig. 1F and Fig. 2D). In some cases, the independently fit K_50_ values can lead to impossible values for ΔG. This can occur if K_50_ is higher than K_50,F_ (observed cleavage is slower than our limit for cleavage in the folded state) or if K_50_ is lower than K_50,U_ (observed cleavage is faster than predicted cleavage in the unfolded state). If the median protease-specific K_50_ or the confidence interval limits for a particular sequence lead to impossible ΔG values for that sequence, we reported dummy values for the corresponding protease-specific ΔG estimates.

The actual number of sequencing counts, as well as the number of counts predicted for all sequences at all concentrations according to the fitted model parameters, are given in Raw_NGS_count_tables.zip and Pipeline_K50_dG.zip.

#### Data selection for Fig. 1E and F

We show all data from Library 3 within the range −2 < ΔG < 5 kcal/mol & log10_K50_trypsin < 1.75 & log10_K50_chymotrypsin < 2.25. We then overlaid the wild-type and four mutants of Protein G measured in Library 2.

#### Replicate analysis of K_50_ (Fig. 1E)

Instead of sampling K_50_ values using 24 samples per protease at one time as described in Step 5 above, we sampled K_50_ values using one experiment set (i.e. 12 samples) and obtained K_50_ for trypsin replicate 1 and 2, and chymotrypsin replicate 1 and 2. Note that we still used the calibrated protease concentrations to improve consistency between replicates. The replicates were conducted on different days using the same preparation of the protein-cDNA complex.

#### Classification of Datasets #1, #2, and #3 based on the quality of the data (For Fig. 2)

All mutational scanning data was classified into nine groups (0 through 8) according to the protocol in Fig. S8. We determined that a mutational scan was high quality (suitable for Dataset #2) if there was minimal missing data, minimal low confidence data, an appropriate slope, intercept, and correlation between the trypsin and chymotrypsin samples, sufficient wild-type stability, and the mutational scan did not include an unusual fraction of stabilizing mutations suggesting poor folding. For inclusion in the smaller Dataset #1, we additionally required that the wild-type stability was lower than 4.5 kcal/mol so that stabilizing mutations could still fall within the assay’s dynamic range. These sequences are considered “Group 0”; the remaining sequences in Dataset 2 are considered “Group 1”. Double mutant sequences were included in Datasets 1 and 2 based on whether the original wild-type mutational scan was included in that dataset.

All sequences in Dataset 1 and Dataset 2 are included in K50_dG_Dataset1_Dataset2.csv. All sequences in this file have an inferred ΔG estimate value, but only sequences in Dataset 1 have a tabulated ΔΔG estimate. Of course, one can calculate ΔΔG for the remaining sequences in Dataset 2, but these ΔΔG values will be biased toward destabilizing mutations because stabilizing mutations would typically be indistinguishable from the wild-type stability. *Note that Datasets 1 and 2 include a small number of sequences with low quality data because these sequences come from mutational scans that are high quality overall*. Although these tables include all K_50_, Δ G, and ΔΔG data (for Dataset 1), low quality data have been filtered out and replaced by a – symbol in the columns labeled “_ML” (for machine learning).

The remaining groups were defined this way:

Group 2: The wild-type protein is too unstable to see sequence-stability relationships. Group 3: Poor expression (low counts in next-generation sequencing) for the assay.

Group 4: Very few destabilizing mutations, suggesting aggregation and/or molten globule formation

Group 5: The wild-type is too stable to see consistency between trypsin and chymotrypsin Group 6 and 7: Low agreement between trypsin and chymotrypsin due to the absence of aromatic amino acids (i.e. chymotrypsin cleavage sites) or the presence of protease recognition sequences in the linker region.

Group 8: Did not fit into groups 2-7, but did not pass the quality metrics for groups 0 and 1.

Dataset #3 includes all data combined (Groups 0-8), even the data from Groups 2-8 that were excluded from Datasets 1 and 2. Although many of the K_50_ values from Groups 2-8 likely reflect factors other than folding stability (e.g. aggregation, low expression, etc.), these data can still be used to train models that directly predict K_50_. Again, a small fraction (~4%) of the K_50_ values in Dataset #3 are low confidence and have been replaced by a – symbol in the “_ML” columns.

#### Principal component analysis (related to Fig. 3)

We performed principal component analysis to determine the factors influencing stability of different amino acids. To this end, we utilized 15,440 sites in the 337 domains that are classified as G0 in the above. All folding stability data were clipped between from −1 to 5 (kcal/mol) because the folding stability outside the dynamic range is not reliable, and then the average of the stability for 20 amino acids for each site was subtracted from the data. Using the data, we performed PC analysis using the scikit-learn library implemented in Python 3.

#### Side chain contacts and burial analysis (Fig. 3D and 6H)

Burial values and contact counts were computed based on AlphaFold models (*18*) of all sequences using the included script Burial_side_chain_contact_Fig3_Fig6.ipynb based on Bio.PDB (*103*) and BioPython (*104*)). The calculation is based on the Rosetta “sidechain_neighbors” LayerDesign method previously reported (*29*). Briefly, to calculate the burial or contacts of residue X, we added up the number of residues in a cone projecting out 9 Å away from the Cβ atom on residue X in the direction of the residue X Cα-Cβ vector. “Burial” (Fig. 6H) indicates the number of Cα atoms in the cone. Contact counts (Fig. 3D) each count different atoms inside the cone: “Side chain contact count” (Fig. 3D) counts all Cβ atoms; “Aromatic side chain contact count” counts all CE2 atoms of Phe, Tyr, and Trp; “Acidic side chain contact count” counts all Glu OE1 and Asp OD1 atoms; and “Basic side chain contact count” counts all Lys NZ and Arg NE atoms.

#### Secondary structure determination (Fig. 3D)

Using the DSSP algorithm (*105, 106*), we obtained secondary structure information based on AlphaFold models.

#### Selection method of site pairs for double mutational analysis (related to Fig. 4)

Double mutants were selected for analysis in two ways. First, we manually selected polar interactions where either amino acid appeared important for stability in single mutational analysis. These pairs were mainly included in Library 3. Second, we used the program confind (*107, 108*) to identify interacting residues. All confind pairs with notable interactions such as polar interactions and cation-π interactions were selected, along with a randomly chosen subset of more common interactions such as hydrophobic interactions. These pairs were included in Library 4.

#### Thermodynamic coupling analysis (related to Fig. 4)

Thermodynamic coupling refers to the change in folding stability due to the interaction between two amino acids after removing folding stability effects from each amino acid individually. To determine this “nonadditivity”, we first modeled our double mutant data using a fully additive model (no thermodynamic coupling). The deviations from this model then reveal the thermodynamic coupling. Our additive model assumes that the absolute stability (ΔG) of each sequence is the sum of an amino acid-dependent term for site one (ΔG_1_) and an amino acid-dependent term for site two (ΔG_2_)

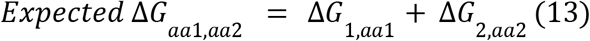

The forty site-specific terms (one ΔG_1_ term for each amino acid at site one and one ΔG_2_ term for each amino acid at site two) are not experimentally measurable; they are inferred based on minimizing the error of the additive model. We used Bayesian inference to infer the forty ΔG_1_ and ΔG_2_ terms for each set of mutants. The inputs to fit the model were the observed 400 ΔG values (20 amino acids at site one x 20 amino acids at site two) for a particular site pair. Using NUTS, we sampled ΔG_1_ and ΔG_2_ by assuming that the 400 observed ΔG values should agree (with small, normally distributed errors) with the expected ΔG values determined by eq. 13. Both expected and observed ΔG values were clipped to the range of −1 to 5 kcal/mol. We used 100 steps of burn-in and used the median of 50 posterior samples as the final values of the ΔG_1_ and ΔG_2_ terms. Using these terms, we calculated the expected (additive) ΔG for each sequence, and then the thermodynamic coupling:

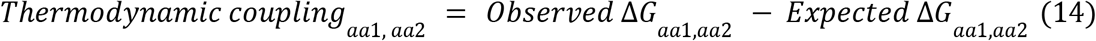

To calculate the uncertainty in the thermodynamic coupling, we re-fit the additive model 50 times by bootstrap resampling of the 400 observed ΔG values. This ensures the ΔG_1_ and ΔG_2_ terms are not overly dependent on a single experimental measurement. The model fitting code is provided in Additive_model_Fig4.ipynb.

#### Wild-type amino acid prediction model (related to Fig. 5)

The classification model in Fig. 5 used a sum of logistic functions with learned amplitudes to define the weighting function. The overall model is defined below:

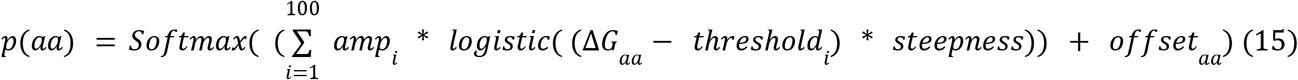

where *p(aa)* is the probability of amino acid *aa, softmax* is the softmax function 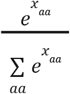, *logistic* is the logistic function f(x) = 1 / (1+e^x^), *i* indexes the 100 logistic functions defining the weighting function, *amp* is the learned vector describing the amplitudes of the logistic functions, *threshold* is the vector describing the centers of the logistic functions, *steepness* defines the steepness of the logistic functions, and *offset* the learned vector (length 19 for the 19 non-Cys amino acids) describing the absolute probability offset for each amino acid.

We used Bayesian inference to infer the *amp* vector (length 100) and *offset* vector (length 19 for the 19 non-Cys amino acids). The logistic *threshold* vector was fixed at 100 evenly spaced points between −2 and 7 kcal/mol. The *steepness* term was fixed at 5. The inputs to fit the model were the observed ΔG values and the wild-type amino acid identities for each site within the natural protein domains. Using NUTS, we sampled *amp* and *offset* by assuming that the observed wild-type amino acids were randomly chosen at each site according to the predicted probability distribution for that site, calculated according to eq. 15. We then reported the median and the standard deviation of 100 posterior samples after removing the initial 500 steps. The fitting script is included in Classification_model_Fig5.ipynb.

#### GEMME analysis (related to Fig. 6)

To calculate the “Normalized averaged GEMME score”, which represents the sensitivity of a wild-type amino acid to substitutions inferred from evolutionary information (“ΔΔE” in the previous reports (*70, 71*)), we ran GEMME (*78*) on each natural amino acid sequence using the default parameters. We computed a single score for each site by averaging the scores of the 19 amino acids (except Cys), and then standardized each domain individually (subtracted the domain’s mean and divided by the domain’s standard deviation) so that the site scores within a domain had a mean of zero and a standard deviation of one. Finally, we flip the sign of the score so that positive values imply high susceptibility to mutations (i.e. very negative raw GEMME scores for non-wild-type amino acids). We define this standardized score for each site as the “Normalized GEMME score”. To build the input multiple sequence alignments, we performed five iterations of the profile HMM homology search tool Jackhmmer (*109, 110*) against the UniRef100 database of non-redundant proteins (*111*) using the EVcouplings framework (*112*). We used the default bitscore threshold of 0.5 bit per residue.

### Supplementary Text

#### Derivation of eq.3 in Fig. 1B

We modeled the cleavage events, where Protease enzymes (E) and protein substrates (S) form an ES complex to produce cleaved protein products (P). The goal is to get a product formation equation in terms of the total product, initial enzyme and substrate concentrations and kinetic constants.

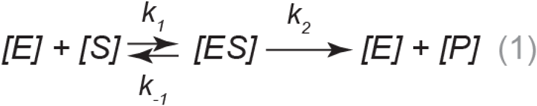

Also, we defined equilibrium constant *K_50_*:

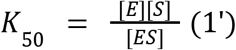

Based on the model (1), we can obtain the following dynamic formulas:

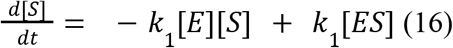

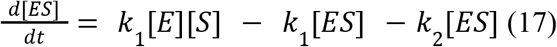

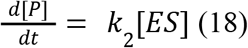

The first two of these are assumed to be at quasi-steady state. The following are additional conservation equations for substrate-product and enzyme:

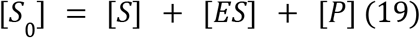

where [S_0_] is initial amount of substrates

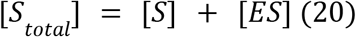

Additionally, the reaction conditions in the study were not substrate-excessive but enzyme-excessive:

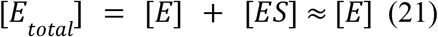

(because [E] >> [ES] or [S])

Using eqs. 1’, 19, and 20, the following can be derived to find an expression for the enzyme-substrate complex in terms of the initial substrate and enzyme concentration:

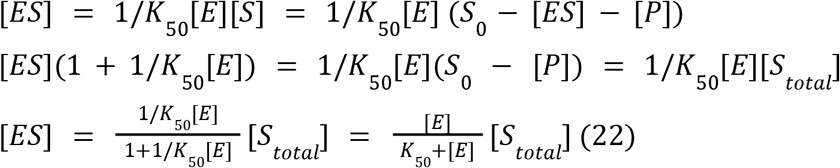

Substituting eq. 22 into eq. 18 and using the approximation[*E_total_*] ≈ [*E*], the an expression for the dynamics of the product formation in terms of enzyme concentration and substrate can be found:

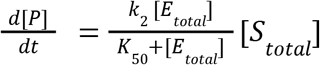

Thus, the observed kinetic rate is 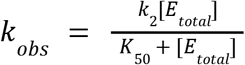 (This eq. 3)

#### Derivation of eq.6 and eq.7 in Fig1B

We modeled the cleavage events, where Protease enzymes (E) and folded substrates (F) or unfolded substrates (U) form a FE or UE complex to produce cleaved protein products (PF or PU). The goal is to get a product formation equation in terms of the total product, initial enzyme and substrate concentrations and kinetic constants. We follow a similar derivation to that above for a single enzyme/substrate:

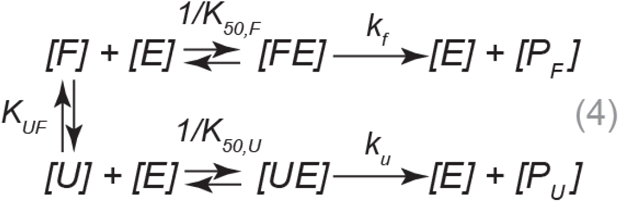

where

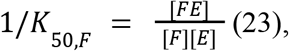

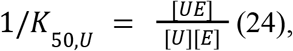

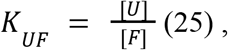

and *k_f_* and *k_u_* are rate arerate constant for cleavage of the bound folded substrates and unfolded substrates. Assuming binding and unbinding equations and the folding and unfolding transition rates are in a quasi-equilibrium then eq 23, 24, and 25 hold throughout the time-course.

We write an equation for the overall product formation:

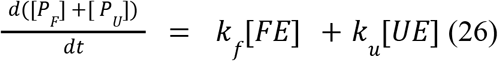

Conservation equations for substrate-product and enzyme in this case are:

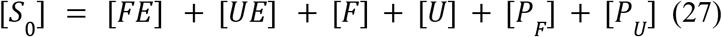

where [S_0_] is initial and total concentration of substrate

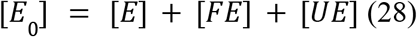

where [E_0_] is the initial concentration of enzyme.

#### Step 1: Write product formation eq. 26 in terms of [FE] and constants only, by substituting for [UE] complex

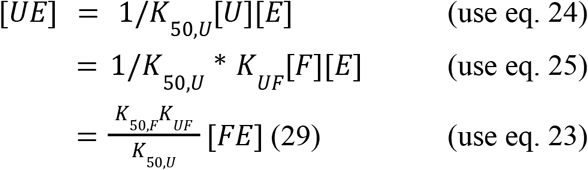

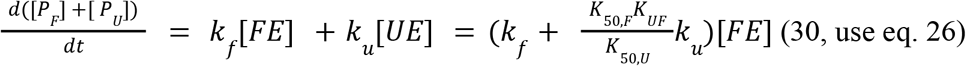

#### Step 2: Replace [FE] dependence with ([S_0_] - [P_F_] - [P_u_]) dependence using conservation laws

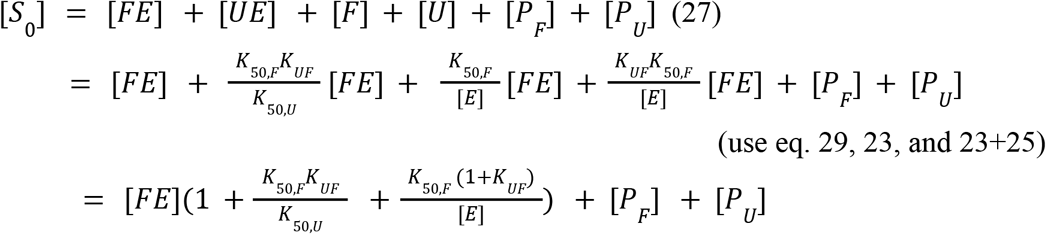

Thus, we get an equation which describes the dependence of [FE] on initial substrates and products, with terms in the denominator that capture sequestration in intermediate bound states.

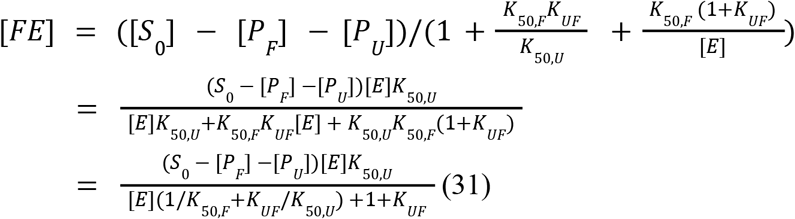

Substituting this into the product formation equation:

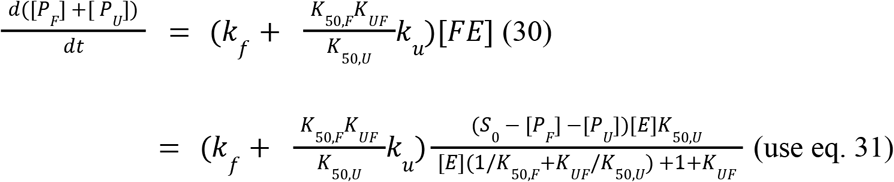

Then, we defined [*P_total_*] = [*P_F_*] + [*P_U_*]

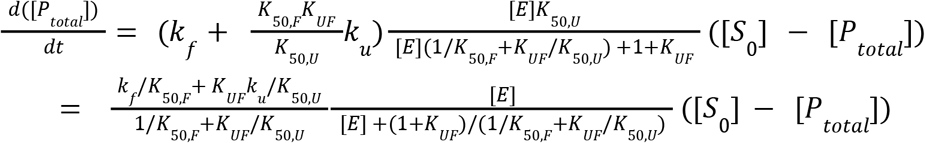

Because the reaction conditions in the study were not substrate-excessive but enzyme-excessive (i.e. [E] >> [S] or [ES]), [E]≈[E0]:

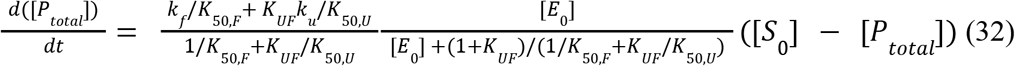

Finally, We can rewrite the product formation eq. 3 in terms of initial substrate concentration, total product, and an observed kinetic rate, which is a function of kinetic rates and initial enzyme concentration,:

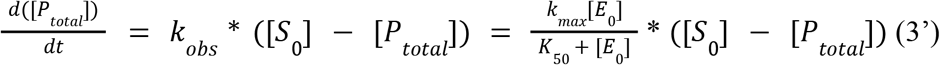

#### Step 3, Derivation eq.6 and eq.7 in Fig. 1B

By comparing eq. 32 with eq. 3’, we can derive the following equations (including eq. 6 in Fig. 1B):

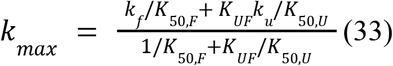

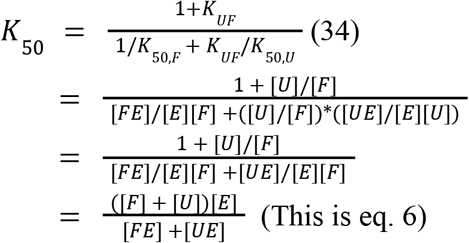

Using eq. 34 to rewriting a formula for *K_UF_* in terms of the half-max reaction rates:

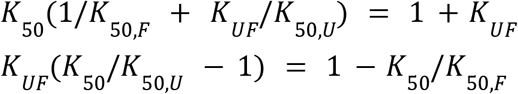

Thus, eq.7 which gives the ratio of unfolded to folded substrate is derived:

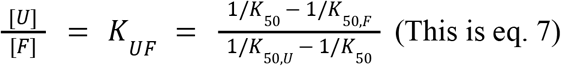

**Fig. S1.**
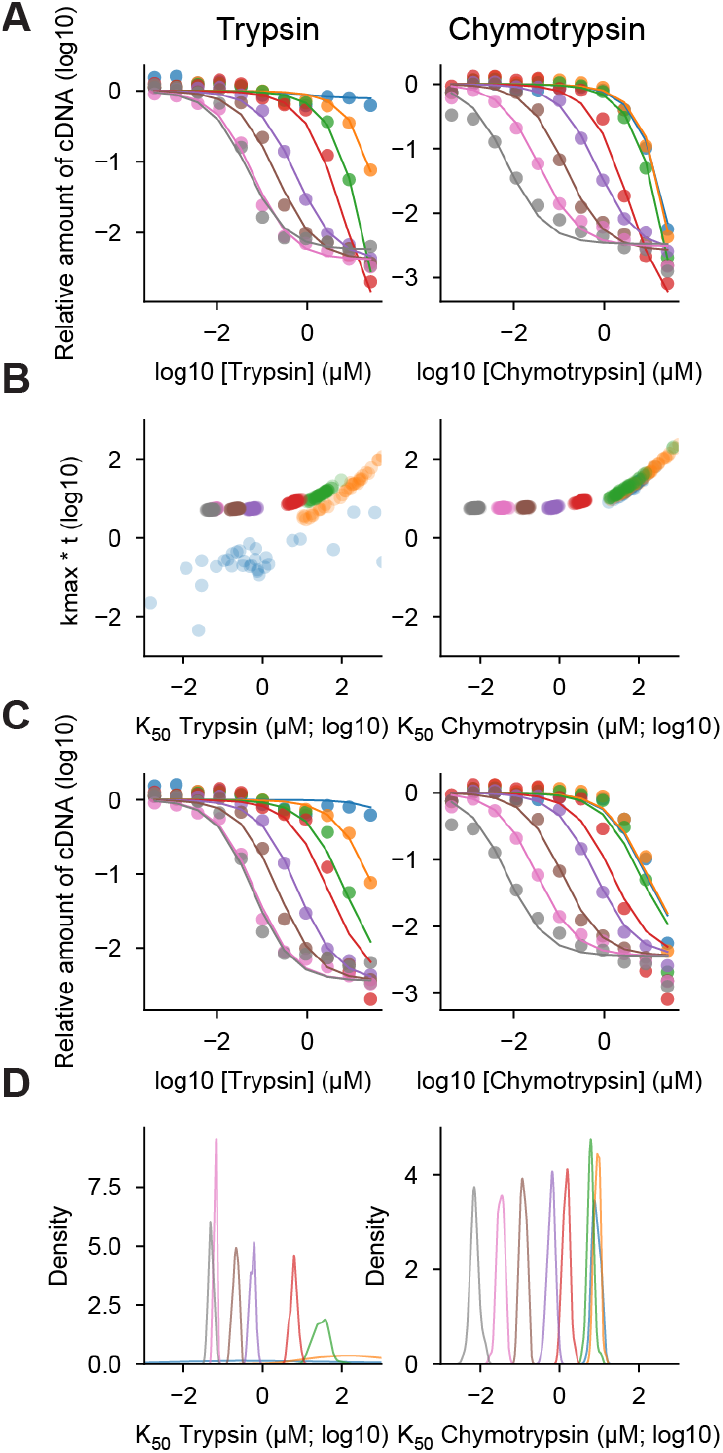
Single turnover model fitting on qPCR data. **(A)** To test the single turnover model, we performed cDNA display proteolysis on a mixture of eight mini protein sequences with diverse folding stability and quantified the surviving amount of each cDNA using qPCR. We then each curve one at a time by Bayesian inference using the single turnover kinetics model in Fig. 1B. We sampled k_max_*t and K_50_ for each sequence. Dots represent the observed cDNA amount quantified by qPCR and lines show the two-parameter fits. **(B)** Posterior distributions of k_max_*t and K_50_ for eight proteins were shown. Whereas K_50_ values vary between different proteins, k_max_*t values (indicating saturation at high protease concentrations) were either constant or unconstrained by the data. **(C)** Based on the analysis (B), we fixed k_max_*t at 10^0.65^ and re-sampled K_50_ for each protein. Dots represent the observed cDNA amount quantified by qPCR (same as in (A)) lines show the one-parameter fits. **(D)** Posterior distributions of K_50_. For trypsin, the K_50_ values for the two most stable proteins (orange and blue) could not be defined because they were too stable and outside of the dynamic range of this proteolysis assay.

**Fig. S2.**
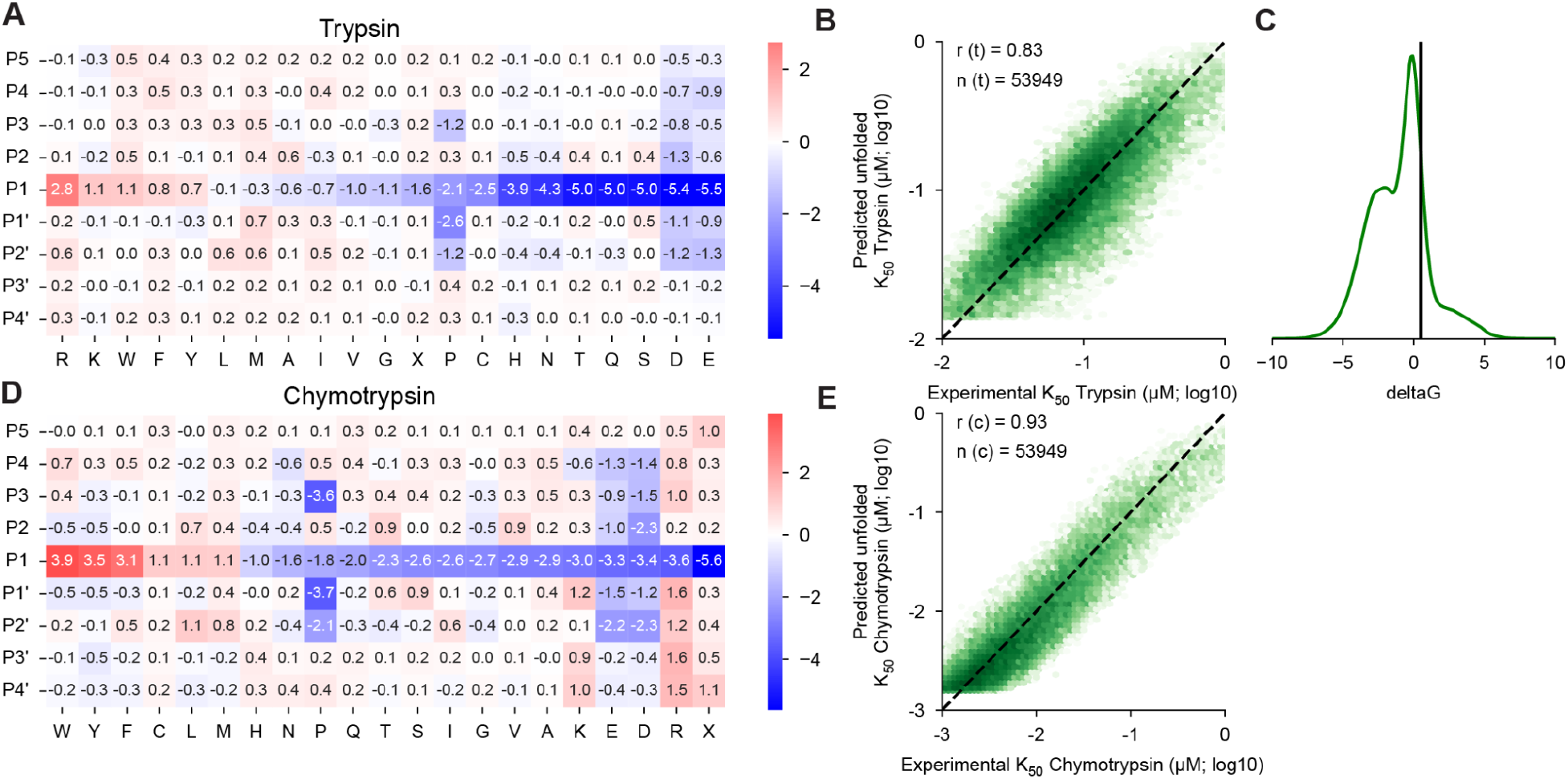
Unfolded state model parameters and goodness of fit. **(A)** Fit parameters for the unfolded state model position-specific scoring matrix (PSSM) for trypsin. The mean of all coefficients (−0.4) was subtracted from the values in the figure to aid visualization. Positive values indicate faster proteolysis and lower predicted K_50,U_ values. By using different prior distribution widths for different rows during fitting, we guided the strongest rate determinants into the center row of each matrix, which we label “P1” (the assay cannot actually identify the specific location of cutting). Overall, the heatmap resembles similar data as previously reported (*29*) and is consistent with known trypsin specificity determinants, including the preference for R/K at P1, the inhibitory effect of P, and the unfavorability of D and E (*113*). **(B)** 2D-histogram showing the overall agreement between the trypsin model (predicted K_50,U_, y-axis) and the data (experimental K_50_, x-axis). Only scrambled sequences with inferred ΔG < 0.5 kcal/mol (where we can assume K_50_ ≈ K_50,U_) are shown (53,949 out of 64,238 total sequences used in training). The Pearson r value is shown. **(C)** Overall distribution of inferred ΔG of all scramble sequences. The vertical line represents 0.5 kcal/mol, which is a threshold used in (B). **(D, E)** As above, for chymotrypsin. As in our previous report (*29*), the coefficients resemble established features of chymotrypsin specificity, including the preference for F/Y/W followed by M/L at P1, the inhibitory effect of P at P3, P1’, and P2’, and the general unfavorability of D and E (*114–116*). The mean of all coefficients (−0.5) was subtracted from the values in the figure to aid visualization.

**Fig. S3.**
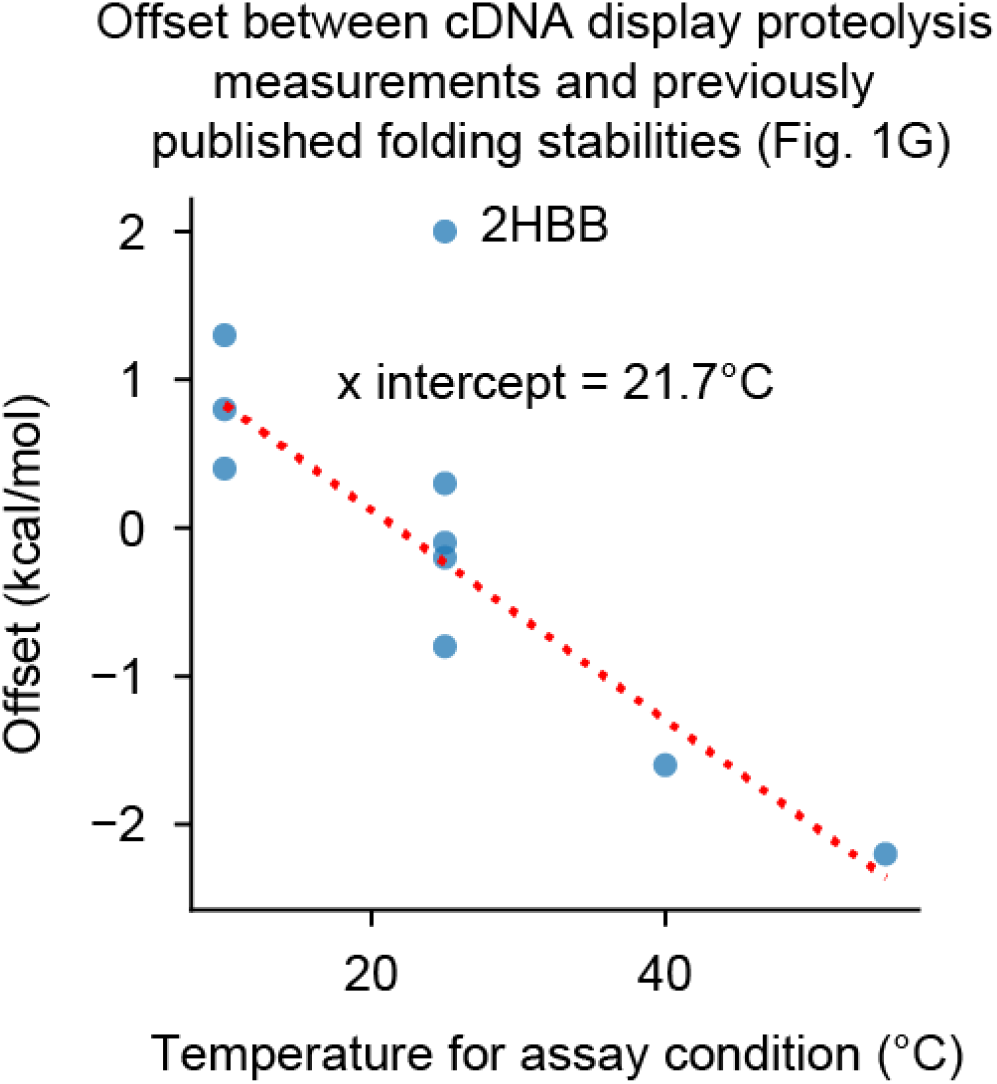
Relationship between offset in Fig. 1G and assay temperature. Previous studies shown in Fig. 1G used diverse conditions including buffer, pH, ion strength, and temperature (see Table. S2) (*52–65*). However, our measurements were all conducted in PBS at room temperature (approximately 22°C). In general, the offsets observed in Fig. 1G are correlated to the temperatures used in the previous studies, suggesting that the assay temperature is the main cause of the offsets. The red line represents a best fit line after removing the 2HBB point. The x-intercept (21.7°C) is close to our assay condition (approximately 22°C). 2HBB (the N-terminal domain of Ribosomal Protein L9) is an outlier and not included in the linear fit; the origin of the offset here is currently unknown.

**Fig. S4.**
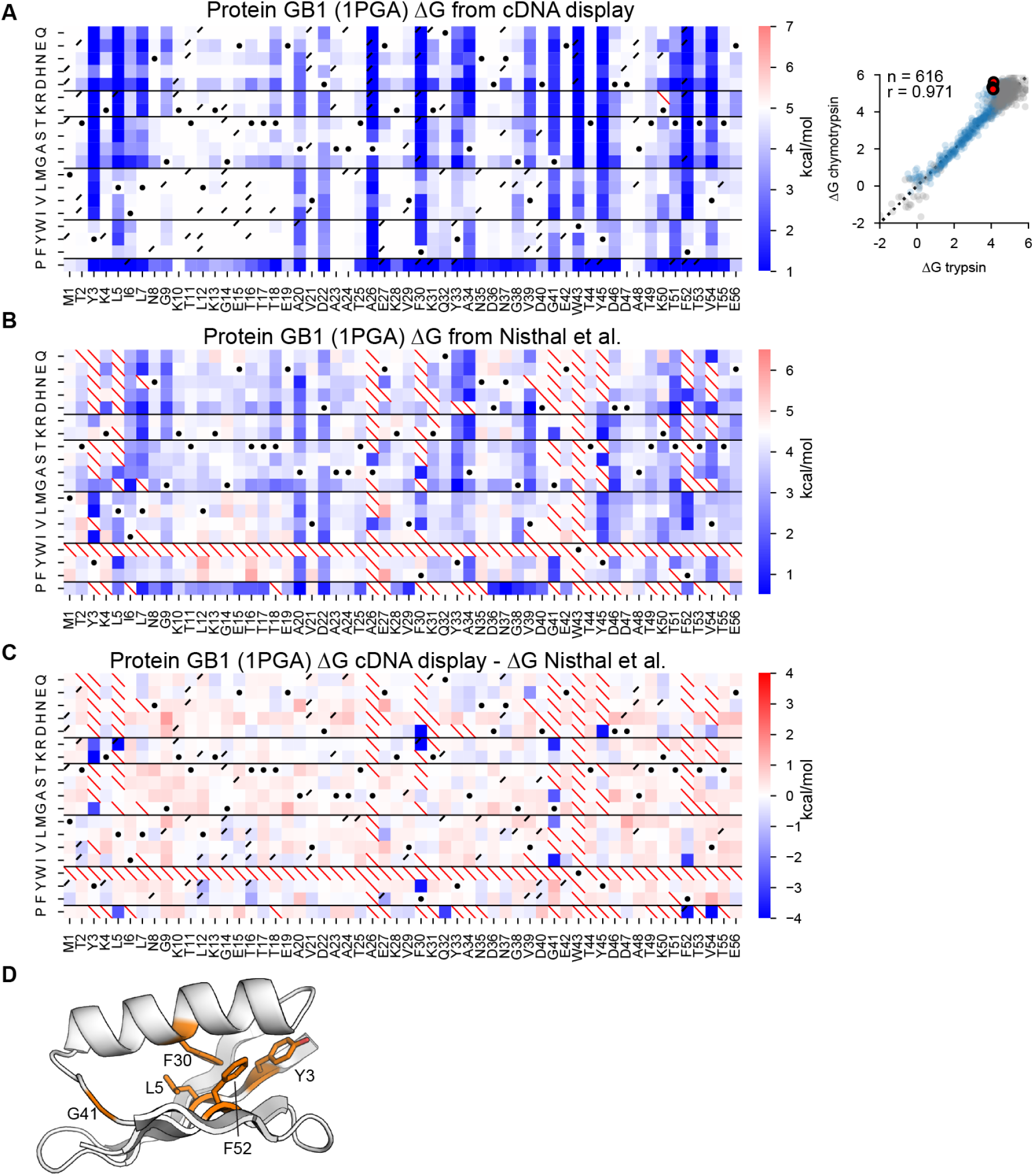
Folding stability discrepancies between cDNA display proteolysis and previous measurements on Protein GB1. **(A)** Left: Mutational scanning results from cDNA display proteolysis. As in Fig. 2, white represents the folding stability of wild-type and red/blue indicates stabilizing/destabilizing mutations. Black dots indicate the wild-type amino acid, red slashes indicate missing data, and black corner slashes indicate lower confidence ΔG estimates, (95% confidence interval > 0.5 kcal/mol), including ΔG estimates near the edges of the dynamic range. Right: Agreement between variant ΔG values independently determined using assays with trypsin (x-axis) and chymotrypsin (y-axis). Multiple codon variants of the wild-type sequence are shown in red, reliable ΔG values in blue, and less reliable ΔG estimates (same as above) in gray. The black dashed lines represent Y=X. Each plot shows the number of reliable points and the Pearson r-value. **(B)** Mutational scanning results from robotics-enabled high-throughput purification and chemical denaturation (*52*), colored as in (A). **(C & D)** Difference heat-map (C) showing the consistency between cDNA display proteolysis (A) and robotics-enabled high-throughput purification and chemical denaturation (B). Dark blue squares indicate highly inconsistent positions where cDNA display proteolysis (A) observes low stability but robotics-assisted chemical denaturation (B) observes high stability. These positions are mainly located in the protein core (shown in D). We hypothesize that many of the inconsistent variants are actually very unstable (as shown by cDNA display proteolysis, A), leading to poorly expressed protein samples that appeared stable in (B) due to the lack of a clear melting signal in chemical denaturation. This would also explain the inconsistencies between closely related mutants seen in (B). For example, the published chemical denaturation data show that Y3R is poorly expressed (no data) along with many other variants at Y3, yet Y3K is measured as very stable (both Y3R and Y3K are unstable in cDNA display proteolysis). The same pattern is seen at L5 in the core: the published data shows that L5K is poorly expressed (no data), yet L5R is measured as very stable (again both L5K and L5R appear unstable in cDNA display proteolysis). The same biophysically inconsistent patterns appear at F30, Q32 (can Q32P in the middle of the helix really be as stable as wild-type?), G41, Y45, F52, and V54. These biophysical inconsistencies at sites where many variants are poorly expressed in the published data suggest to us that the cDNA display proteolysis measurements are more likely to be correct.

**Fig. S5.**
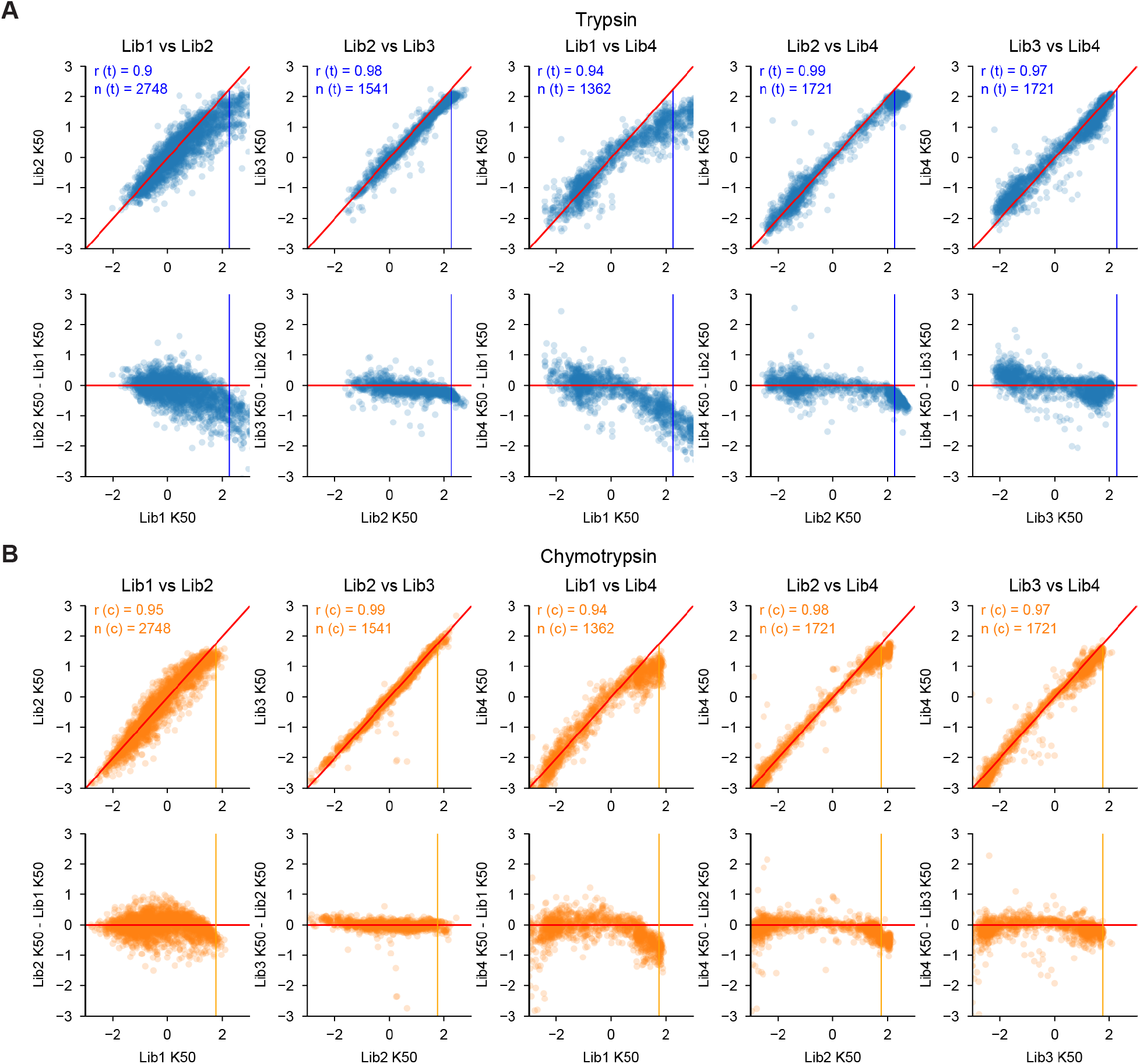
Consistency of K_50_ measurements across libraries. **(A and B)** To examine the consistency between K_50_ (μM) values measured in different libraries, we included identical sequences (potentially with different padding at the termini) in multiple libraries. For each pair of libraries with overlapping sequences, we show the K_50_ values for those sequences in both libraries for trypsin (A) and chymotrypsin (B). The top row shows raw K_50_ values for overlapping sequences in each library; the second row shows the difference in K_50_ estimates plotted against the K_50_ in one of the libraries. The red diagonal line shows Y=X in the top row and Y=0 (i.e. identical K_50_ estimates) in the bottom row. Blue/orange vertical lines show K_50,F_; all K_50_ values above K_50,F_ are treated as equivalent. Each plot is annotated at the top-left with the total number of overlapping sequences and Pearson r-value between the libraries.

**Fig. S6.**
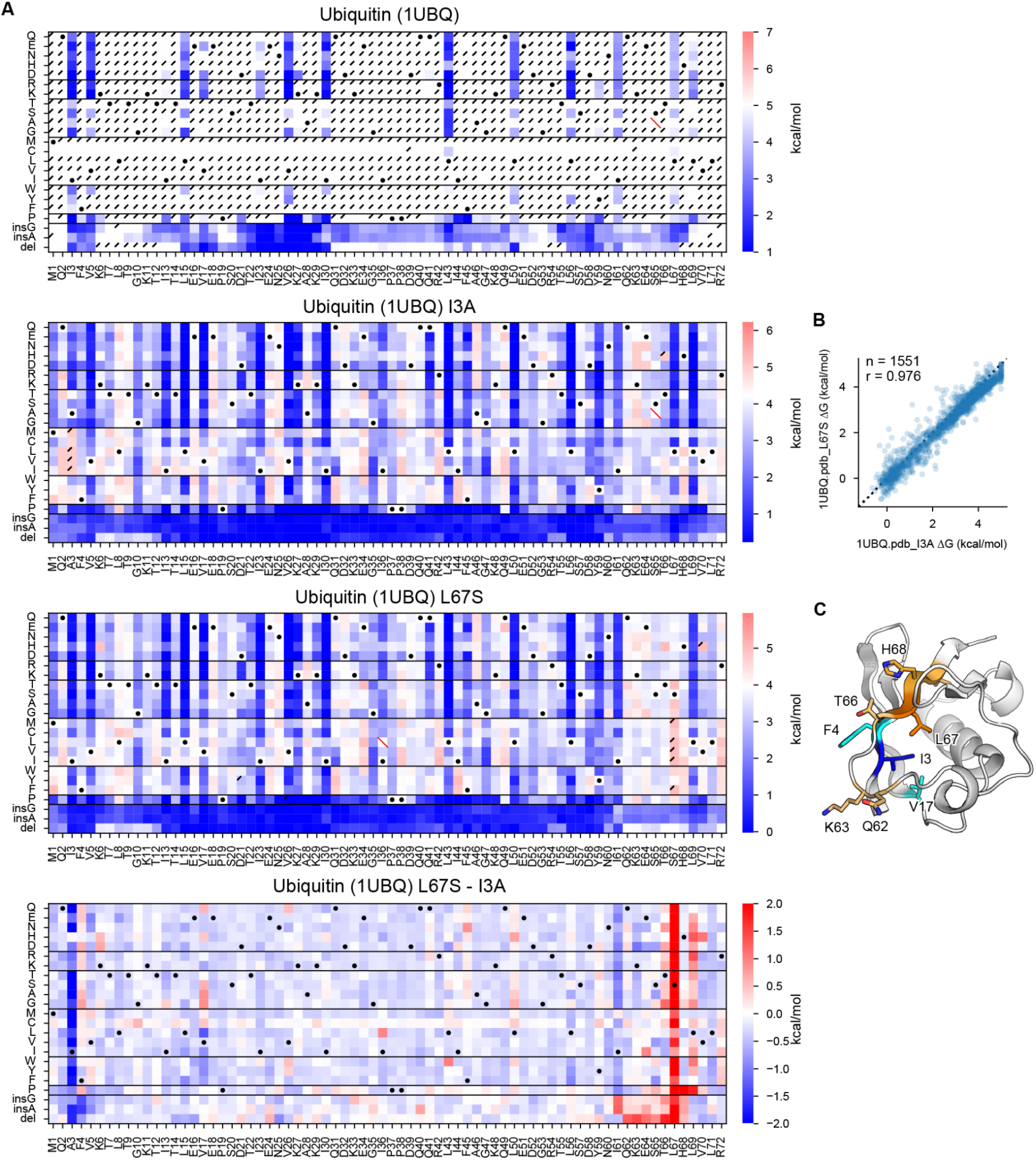
Heat maps for a stable domain (Ubiquitin; 1UBQ) and its destabilizing mutants. **(A)** Mutational scanning results for human erythrocytic ubiquitin (1UBQ) and its destabilizing mutant backgrounds (I3A and L67S). Heat maps show the ΔG of wild-type ubiquitin (top), ubiquitin I3A (middle-top), ubiquitin L67S (middle-bottom), and the difference (ΔΔG) between two mutant backgrounds (bottom) for substitutions, deletions, and Gly and Ala insertions at each residue. In the three ΔG heat maps, white represents the folding stability of the wild-type and red/blue indicates stabilizing/destabilizing mutations. Black dots indicate the background (wild-type or mutant) amino acid, red slashes indicate missing data, and black corner slashes indicate lower confidence ΔG estimates, (95% confidence interval > 0.5 kcal/mol), including ΔG estimates near the edges of the dynamic range. **(B)** Consistency between mutant stabilities measured in the I3A background (x-axis) and L67S (y-axis) background. The plot is annotated with the number of points and the Pearson r value. **(C)** Ubiquitin structure highlighting the mutant points (I3 and L67) and the residues with a different effect on stability between two mutational backgrounds.

**Fig. S7.**
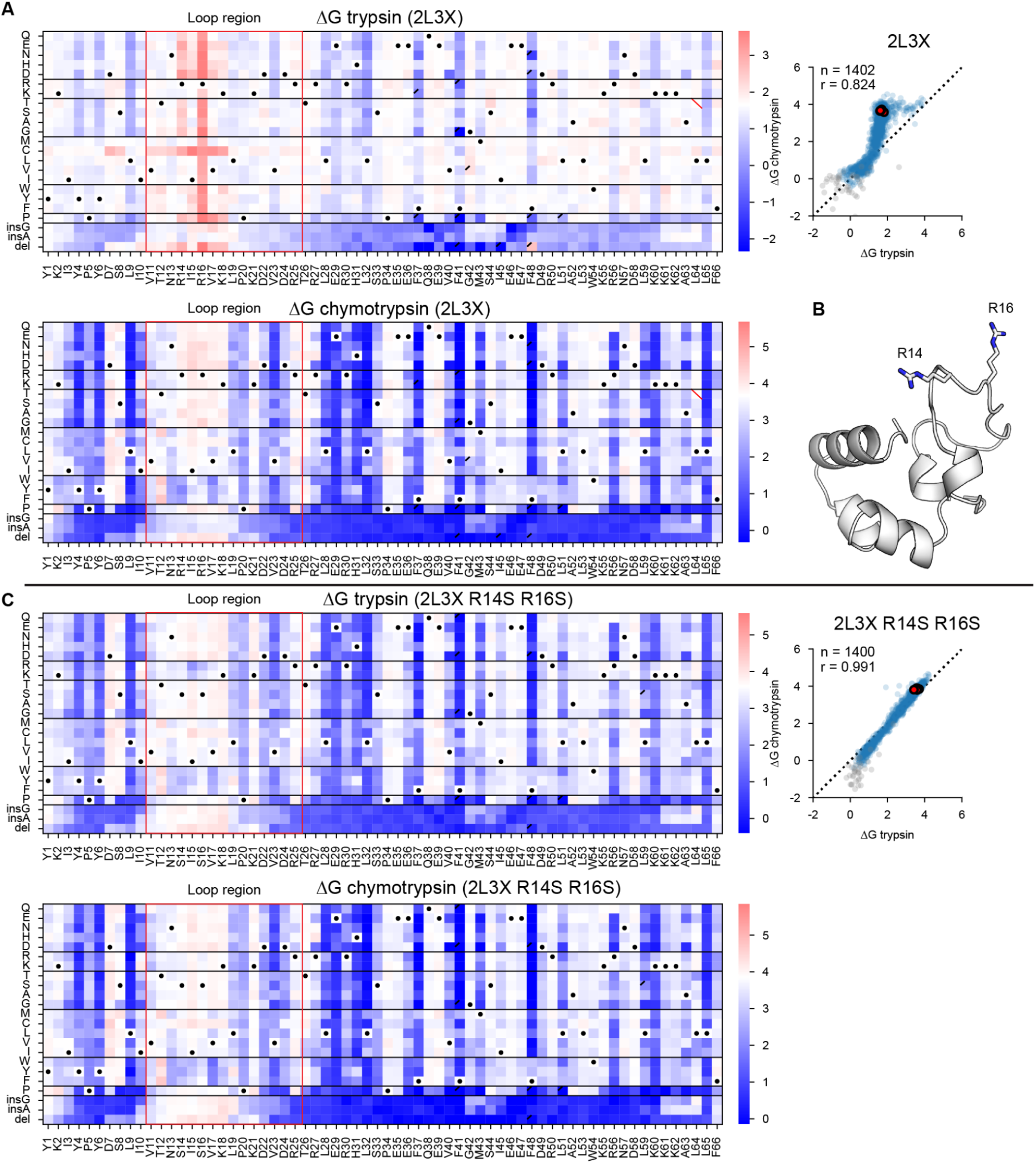
Heat maps for one domain with trypsin cleavage sites in loop region. **(A)** Mutational scanning results for 2L3X, which includes trypsin cleavage sites in the loop region. Left: Heat maps show the ΔG trypsin (top) and chymotrypsin challenge (bottom) for substitutions, deletions, and Gly and Ala insertions at each residue, with our one-indexed numbering at the bottom. Black dots indicate the wild-type amino acid, red slashes indicate missing data, and black corner slashes indicate lower confidence ΔG estimates, (95% confidence interval > 0.5 kcal/mol), including ΔG estimates near the edges of the dynamic range. The colored boxes highlight sites in the flexible loop region. Right: Replicates of the wild-type sequence are shown in red, reliable ΔG values in blue, and less reliable ΔG estimates (same as above) in gray. The black dashed lines represent Y=X. Each plot shows the number of reliable points and the Pearson r-value. The dots show a reverse ‘L’ shape due to the cleavage of the flexible loop region in the trypsin challenge. **(B)** 2L3X structure highlighting Args in the loop region (R14 and R16). **(C)** Same as (A) for 2L3X with replacement of Arg in the loop (R14 and R16) with Ser. In this deep mutational scanning, we observed higher consistency between trypsin and chymotrypsin challenges because we removed sites that could be cleaved in the folded state.

**Fig. S8.**
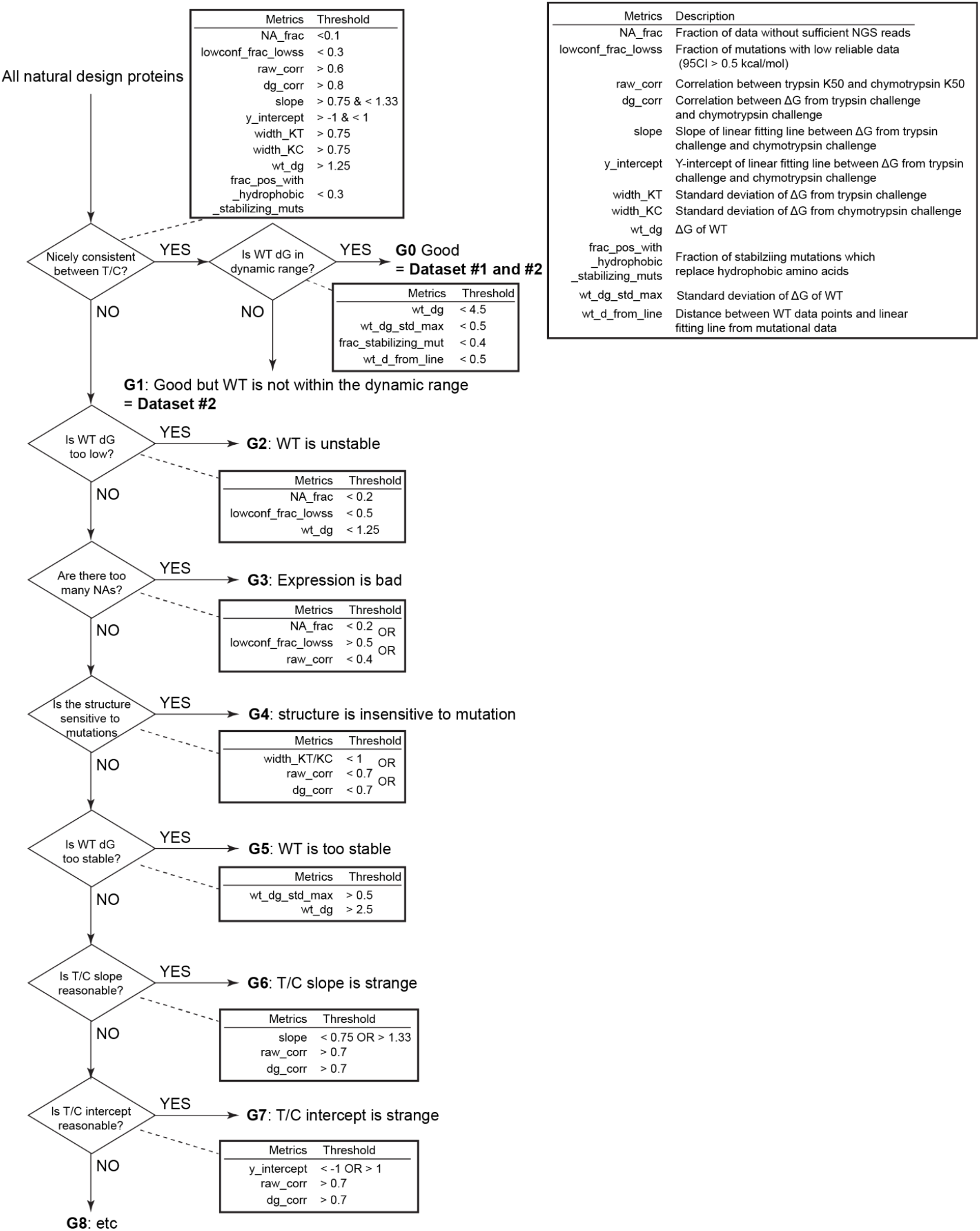
Classification of deep mutational scanning results. We classified all deep mutational scanning results into nine groups shown in Fig. 2B. Here, we show the classification criteria. The description of all metrics is also included in Table. S3, and the metrics of all domains for the classification are included in Single_DMS_list.csv.

**Fig. S9.**
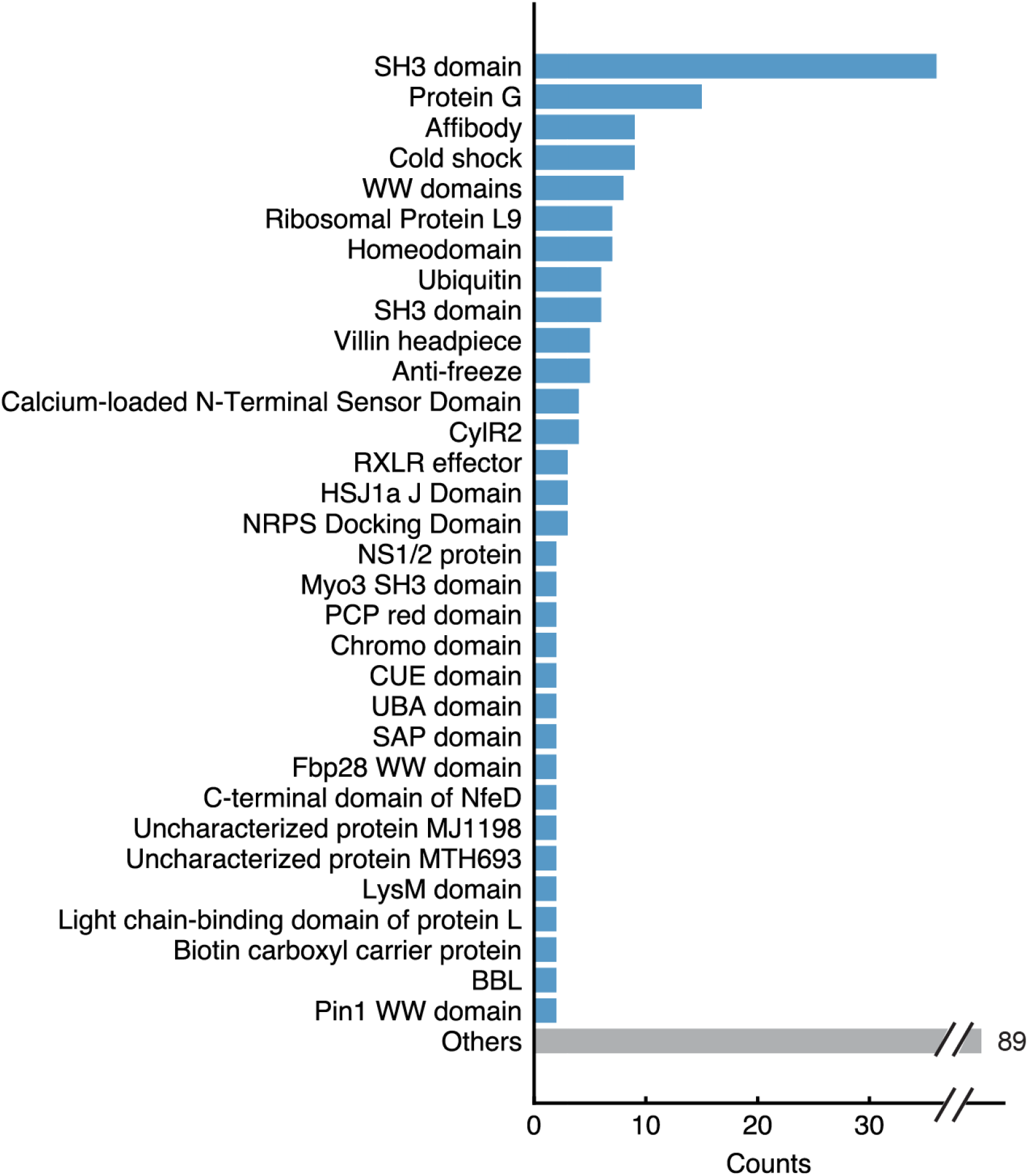
Classification of the natural protein domains investigated in cDNA display proteolysis. Comprehensive group list of wild-type structures classified as G0 in Fig. 2B grouped into domain families.

**Fig. S10.**
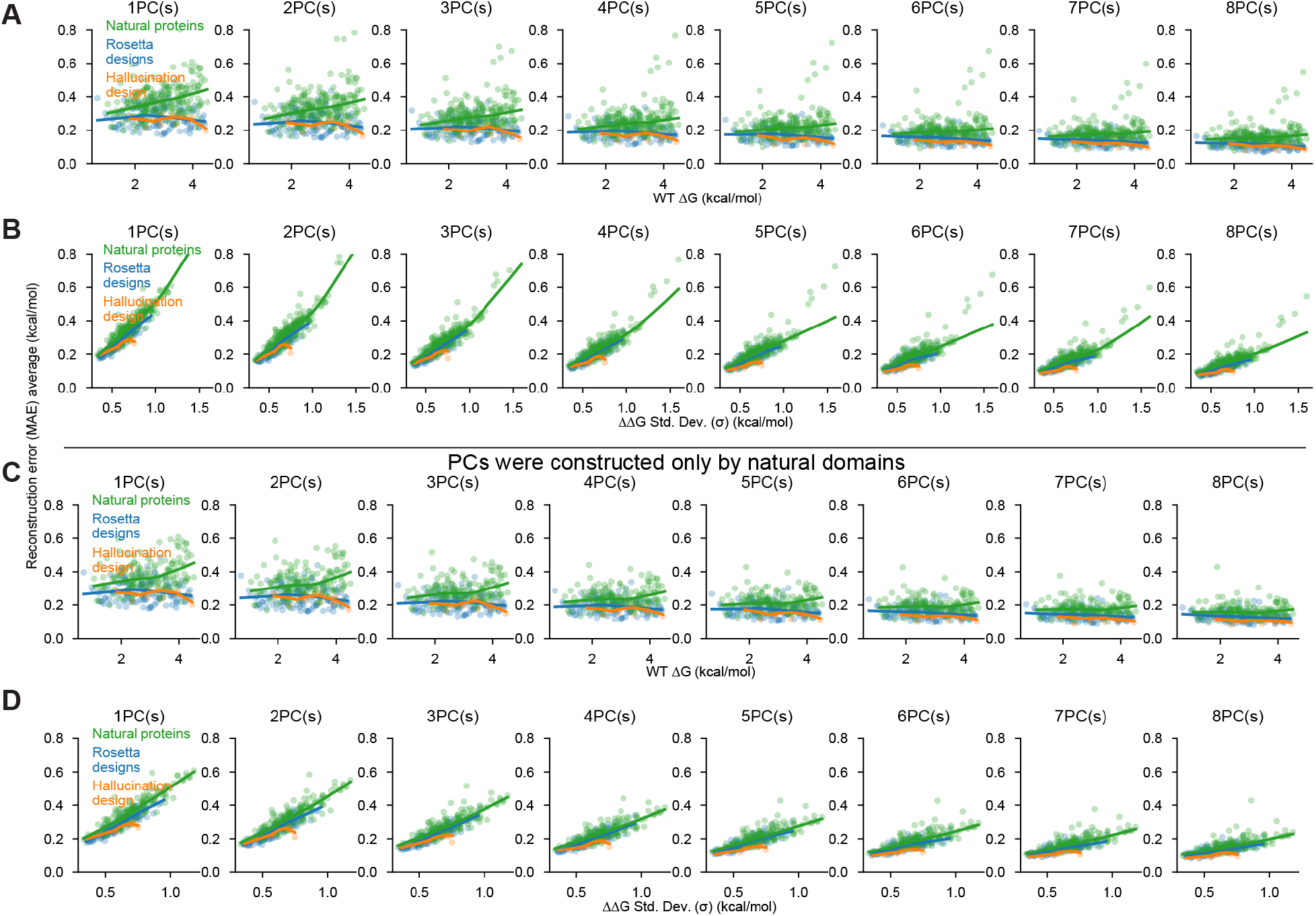
PC analysis with different numbers of PCs. **(A and B)** Relationship between reconstruction error (MAE) using 1-8 PCs (MAE, y-axis) and wild-type stability (A, x-axis) or variance in the ΔΔG data (B, x-axis). Colors represent protein structures grouped into natural proteins (green), Rosetta designs (blue), and hallucination designs (orange). Lines show LOESS fits to the data. **(C and D)** Same as (A) and (B). PCs were constructed using only natural domains. The overall tendency is the same as (A) and (B).

**Fig. S11.**
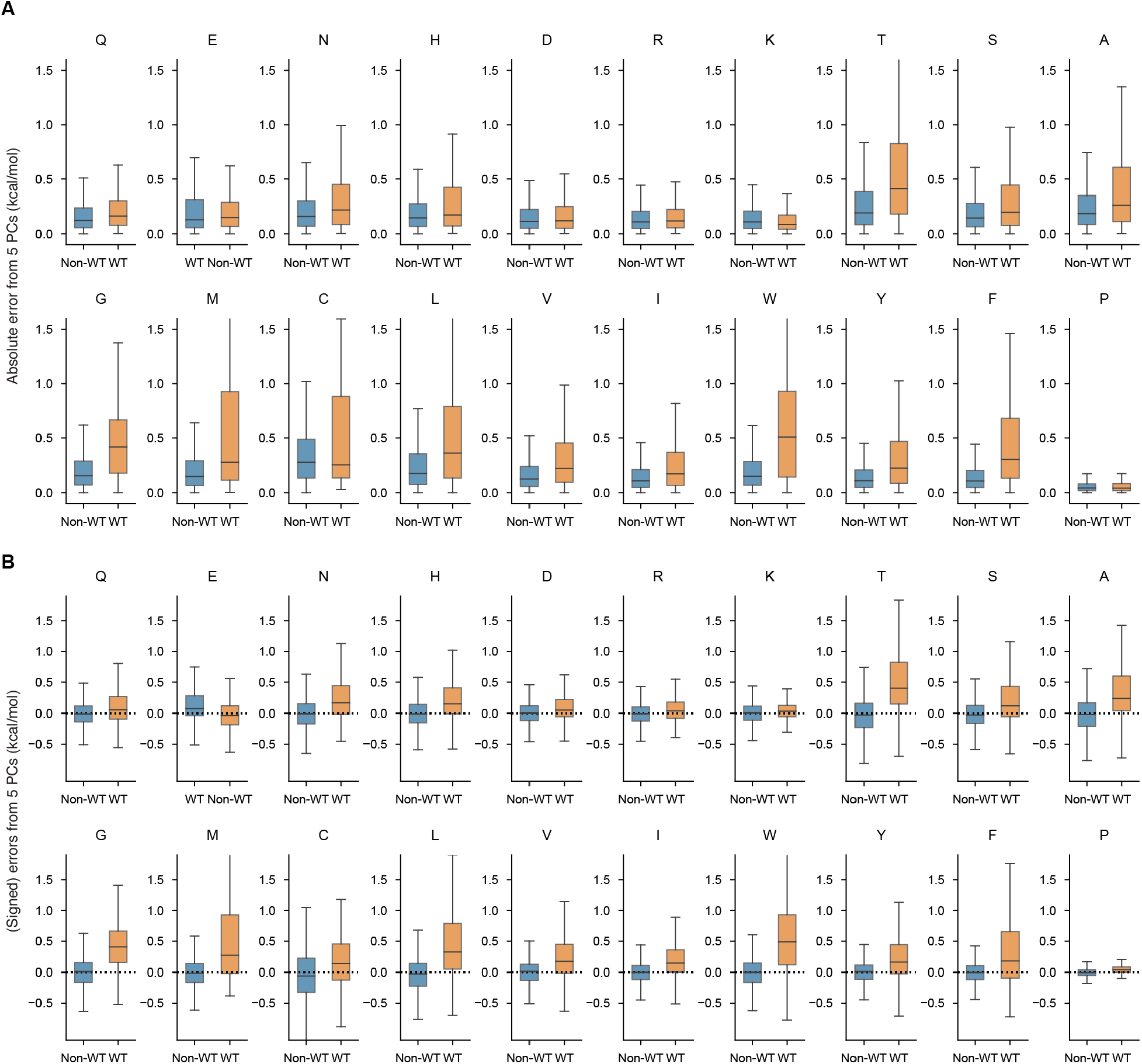
Errors between observed ΔG and reconstructed ΔG by five PCs for wild-type (WT) or non-WT residues for 20 amino acids. Absolute (A) and signed (B) errors between observed ΔG and reconstructed ΔG (from five PCs) are shown. Wild-type residues tend to show larger (A), more positive errors (B), meaning that the five-PC model underestimates wild-type stabilities.

**Fig. S12.**
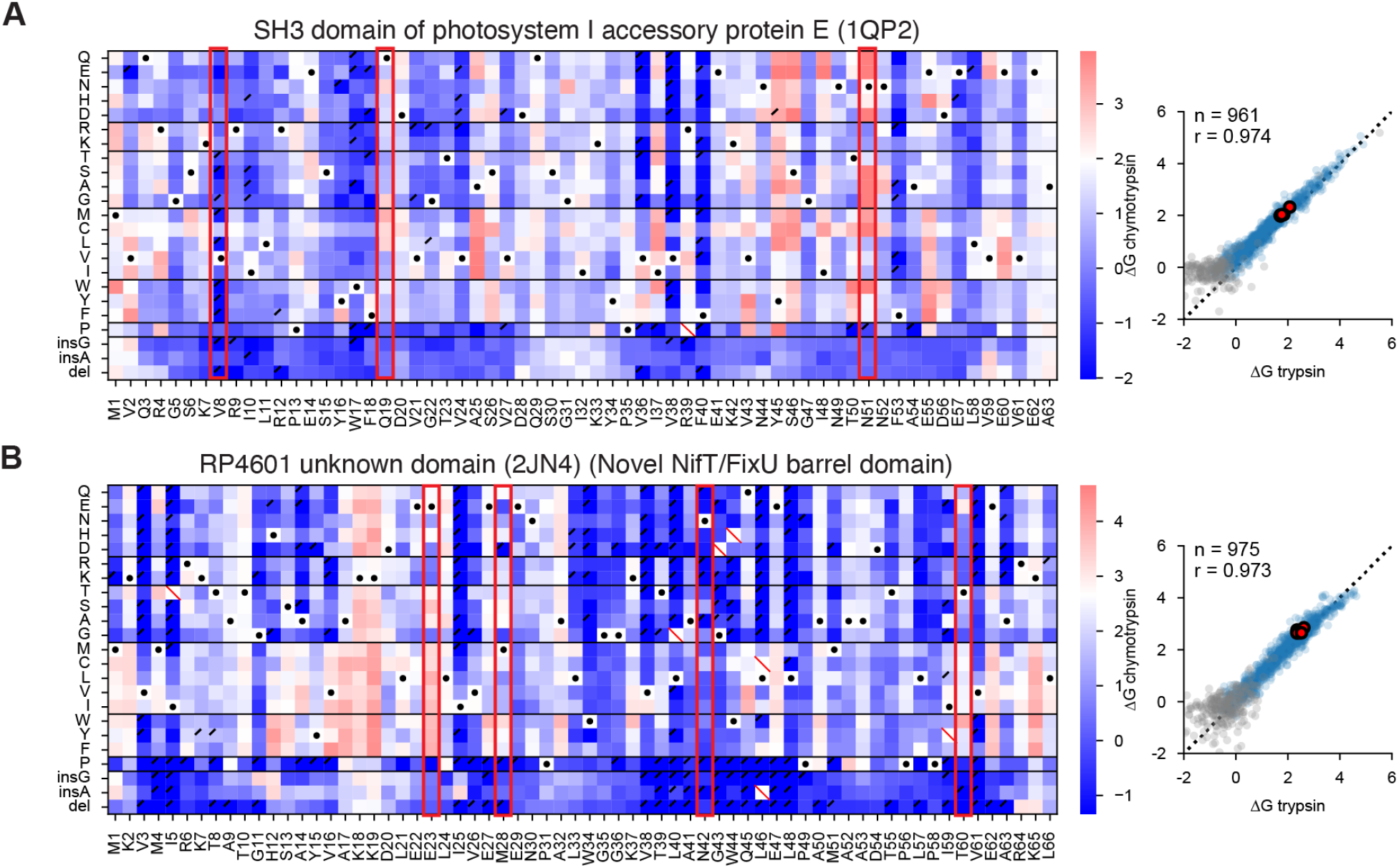
Heatmaps for the notable domains with large errors between observed ΔG and reconstructed ΔG by five PCs. Mutational scanning results for the two notable domains described in Fig. 3G. Left: Heat maps show the ΔG for substitutions, deletions, and Gly and Ala insertions at each residue, and our one-indexed numbering at the bottom. Black dots indicate the wild-type amino acid, red slashes indicate missing data, and black corner slashes indicate lower confidence ΔG estimates, (95% confidence interval > 0.5 kcal/mol), including ΔG estimates near the edges of the dynamic range. Red boxes highlight the seven notable residues with large MAE described in Fig. 3H. Right: Agreement between variant ΔG values independently determined using assays with trypsin (x-axis) and chymotrypsin (y-axis). Multiple codon variants of the wild-type sequence are shown in red, reliable ΔG values in blue, and less reliable ΔG estimates (same as above) in gray. The black dashed lines represent Y=X. Each plot shows the number of reliable points and the Pearson r-value.

**Fig. S13.**
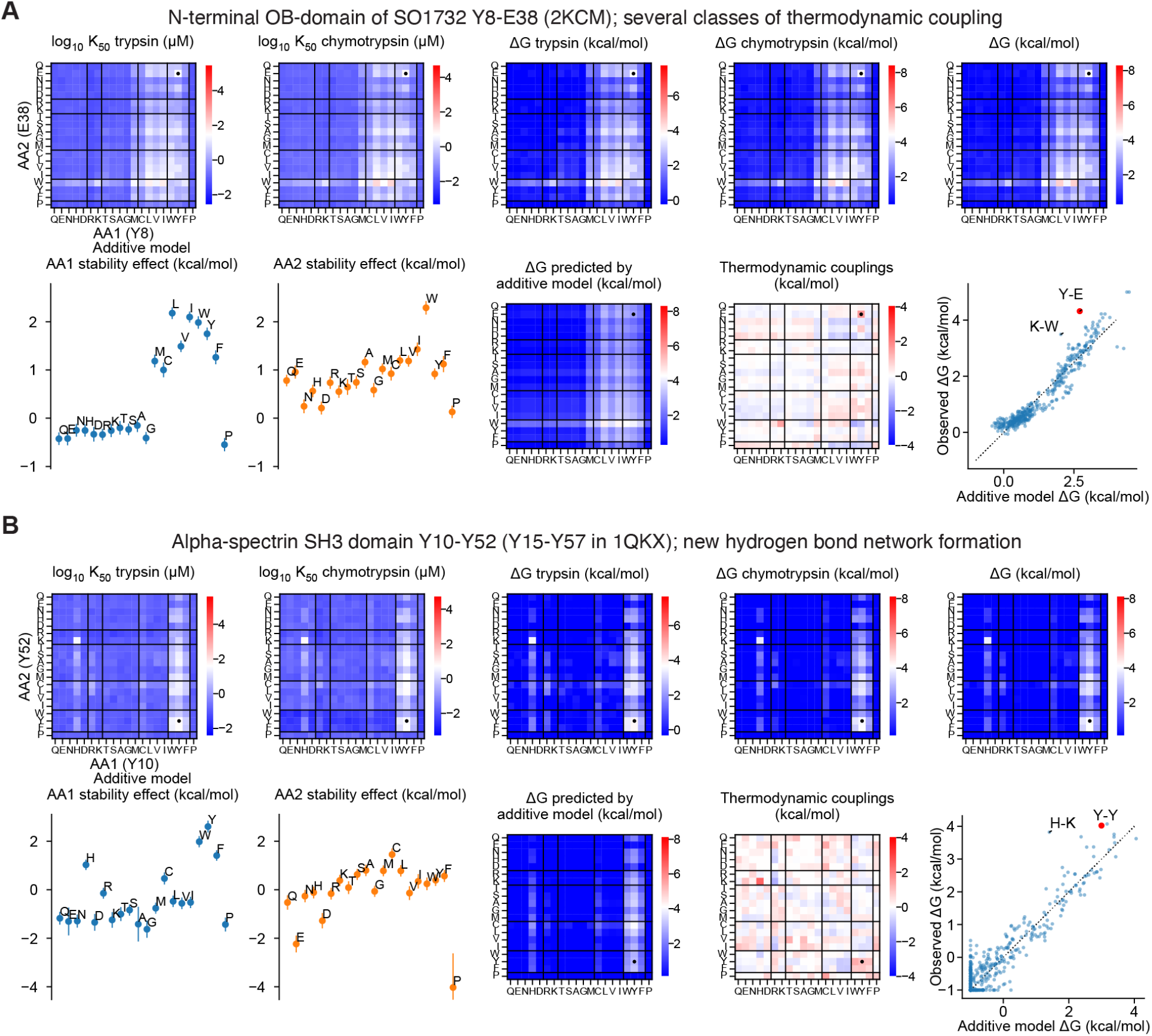
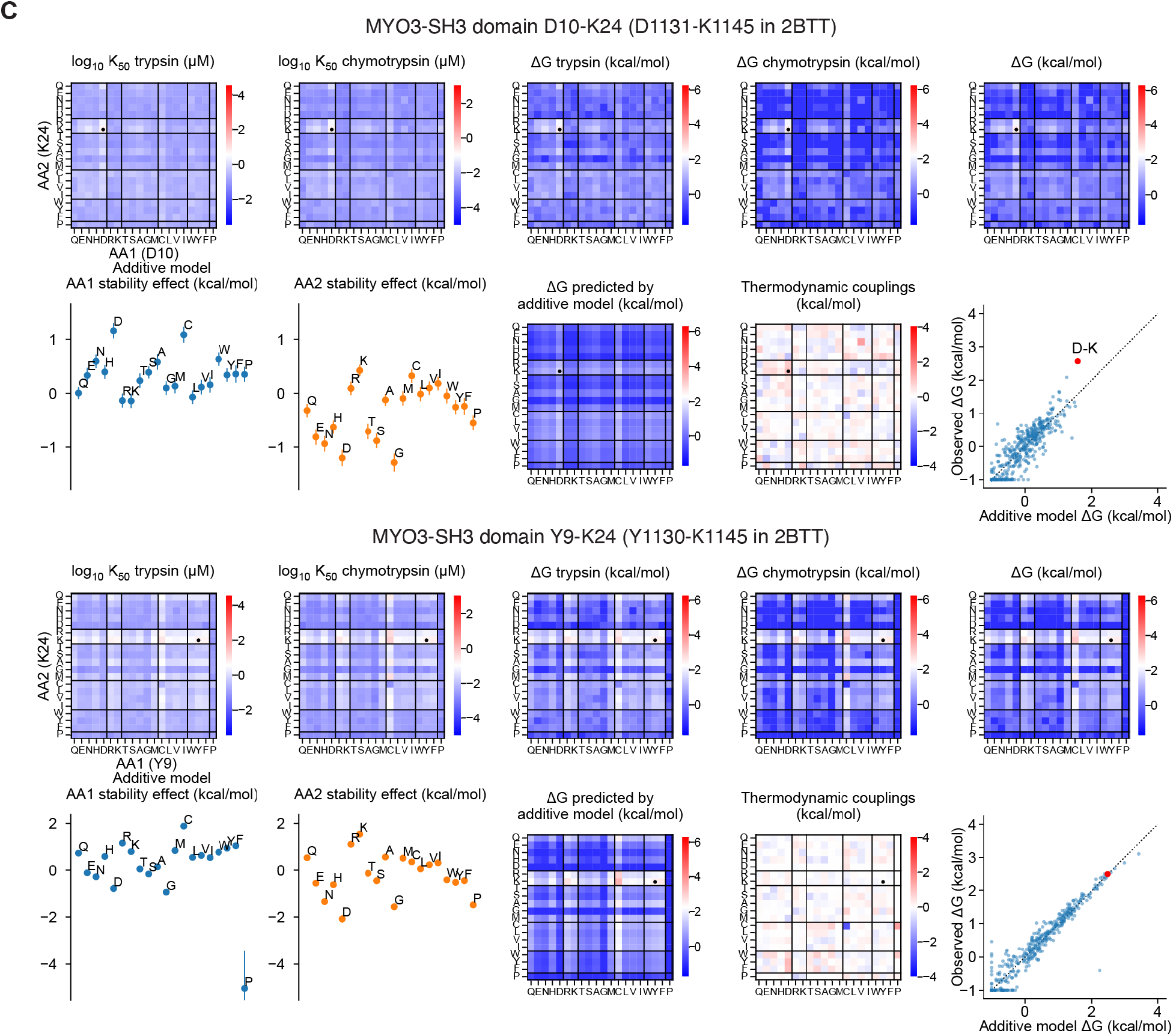
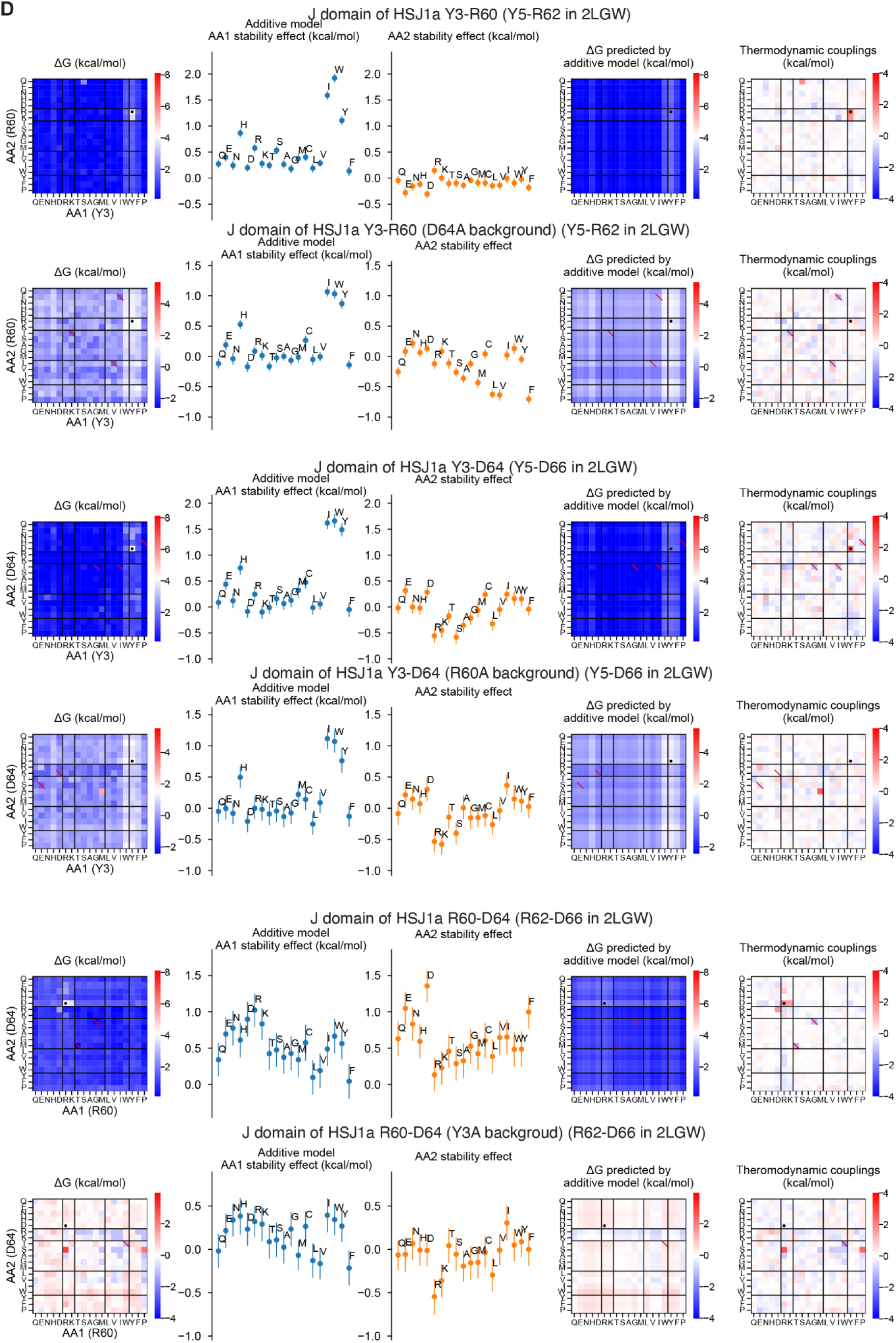
Comprehensive double mutational data for the notable amino acid pairs. **(A and B)** Analysis of thermodynamic coupling for two notable amino acid pairs. In the first row, we show stabilities for all 20×20 double mutants according to five different experimental metrics. From left to right, we show trypsin K_50_, chymotrypsin K_50_, ΔG inferred from trypsin experiments, ΔG inferred from chymotrypsin experiments, and ΔG inferred from both sets of experiments together. In the second row, we show the results of the additive model. From left to right, the first two plots show the inferred single amino acid terms for all 20 amino acids in the first and second sites of the amino acid pair. Error bars represent the standard deviation of the posterior distributions. The middle heatmap shows stability (ΔG) for all amino acid pairs according to the additive model (the sum of the two single amino acid terms). The fourth plot shows the observed thermodynamic coupling; e.g. the experimental ΔG (rightmost plot in the first row) minus the prediction from the additive model (middle plot of the second row). The final scatter plot shows experimental stabilities for all double mutants (y-axis) plotted against the results from the additive model (x-axis). **(C)** Same analysis as (A) and (B) for two site pairs in MYO3-SH3 domain (2BTT). **(D)** Analysis of thermodynamic coupling for all amino acid pairs from a notable amino acid triple. The same amino acid substitutions were also performed for the mutant background with the third amino acid replaced by Ala. From left to right, we show the stabilities (ΔG) of all pairs of amino acids, the single amino acid terms in the additive model (error bars show the standard deviation of the posterior distribution), the stabilities for all pairs according to the additive model, and the thermodynamic coupling for all pairs of amino acids.

**Fig. S14.**
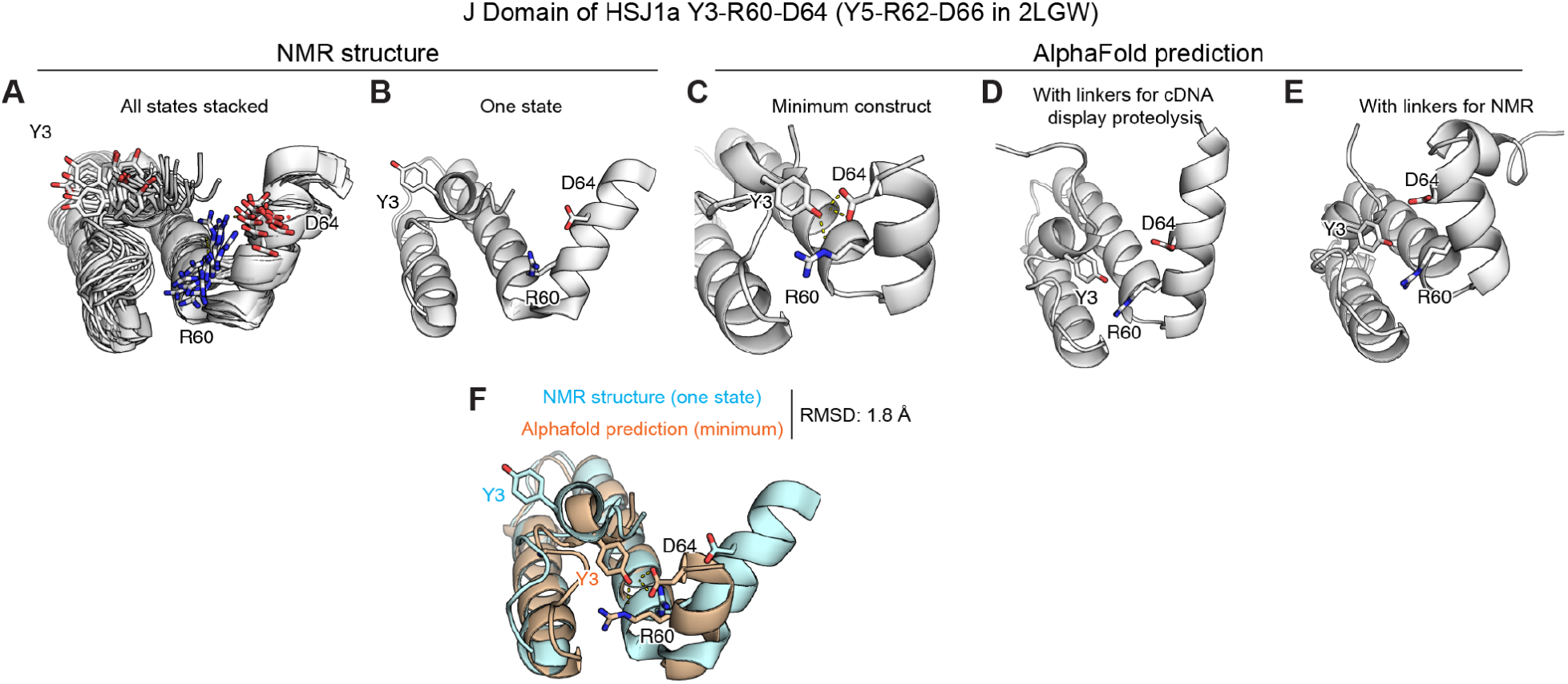
Comparison of AlphaFold model and NMR structure for J domain of HSJ1a. Structure of J domain in HSJ1a (2LGW). We show NMR structure of all states stacked (A) and the first state (B), and AlphaFold predicted structures for the minimum construct (the variable segment in cDNA display) (C), the construct with linkers for cDNA display proteolysis (D), and the exact sequence used for NMR (E). In (F), we overlay the first state of the NMR ensemble (cyan) with the AlphaFold structure (orange) of the minimal construct.

**Fig. S15.**
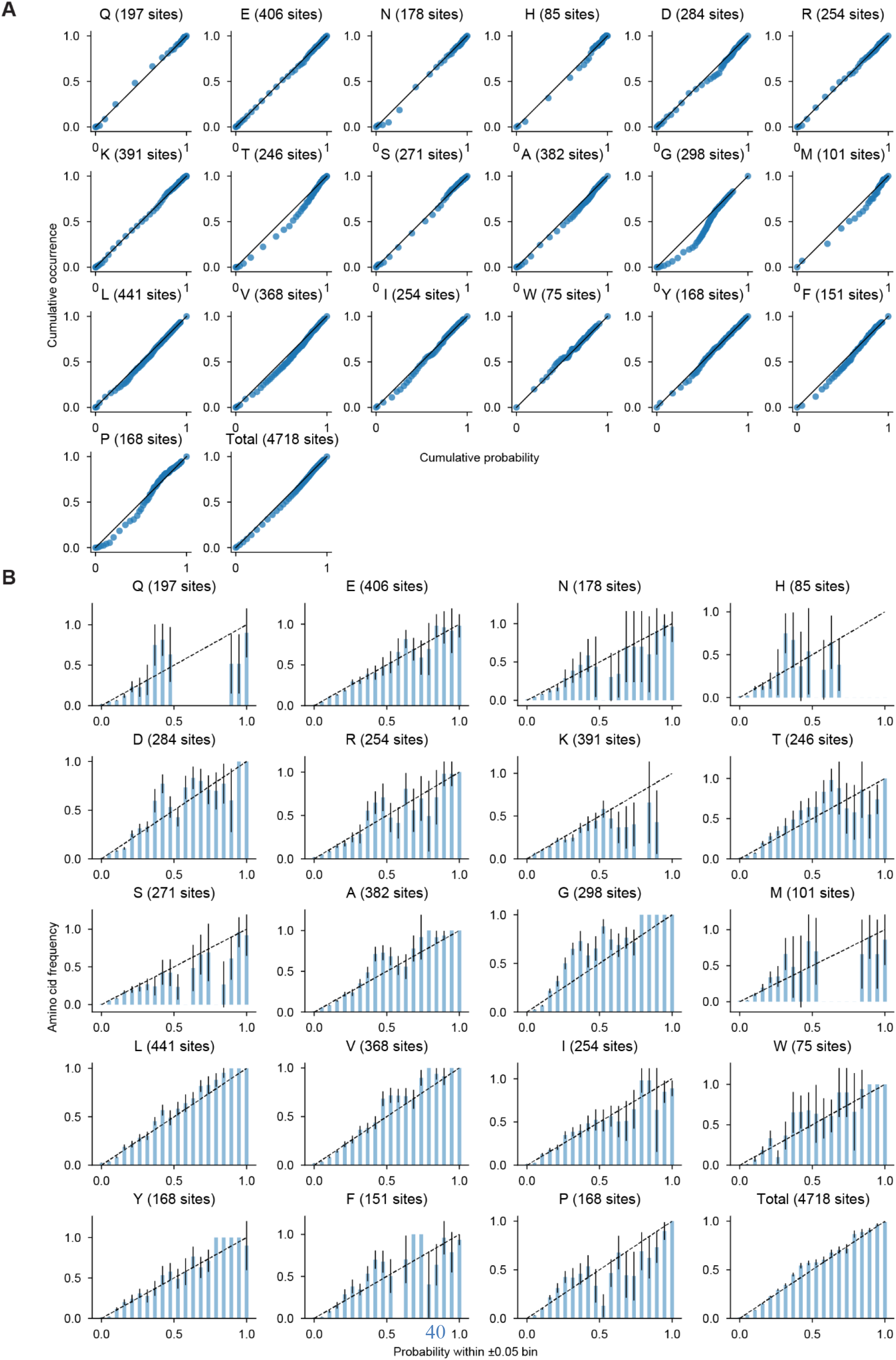
Testing calibration of classification model for predicting wild-type amino acids. **(A)** Relationship between predicted cumulative probability and observed cumulative occurrence for each of 19 amino acids and total data. For each of the 19 amino acids (excluding Cys), we order all 4,718 sites from lowest to highest probability for that amino acid, then step through the sites in that order while plotting the fraction of the total cumulative probability (x-axis) and the fraction of all occurrences of that amino acid (y-axis). For the “Total” plot, we order all 89,642 (4,718*19) amino acid possibilities at all sites from lowest probability to highest probability, then step through all amino acid possibilities in that order while plotting the fraction of the total cumulative probability (x-axis) and the fraction of all actual amino acid occurrences (y-axis). The black diagonal lines show Y=X. **(B)** Relationship between modeled amino acid probabilities and actual amino acid frequencies. For each of the 19 amino acids (excluding Cys), we binned all 4,718 sites into 20 bins according to the probability of that amino acid. Bins are spaced every 0.05 probability units and each bin has a width of 0.1, so sites can appear in two neighboring bins. For each bin (x-axis), the bar shows the true frequency of that amino acid in that bin (y-axis); error bars indicate the standard deviation of the true frequency from bootstrap resampling of all the sites. The black diagonal lines show Y=X (e.g. the predicted probability matches the true frequency). For the “Total” plot, we binned all 89,642 (4,718*19) amino acid possibilities at all sites as before, then counted the fraction of matching amino acids in each bin. Error bars represent the standard deviation of the frequencies from bootstrap resampling of all sites.

**Fig. S16.**
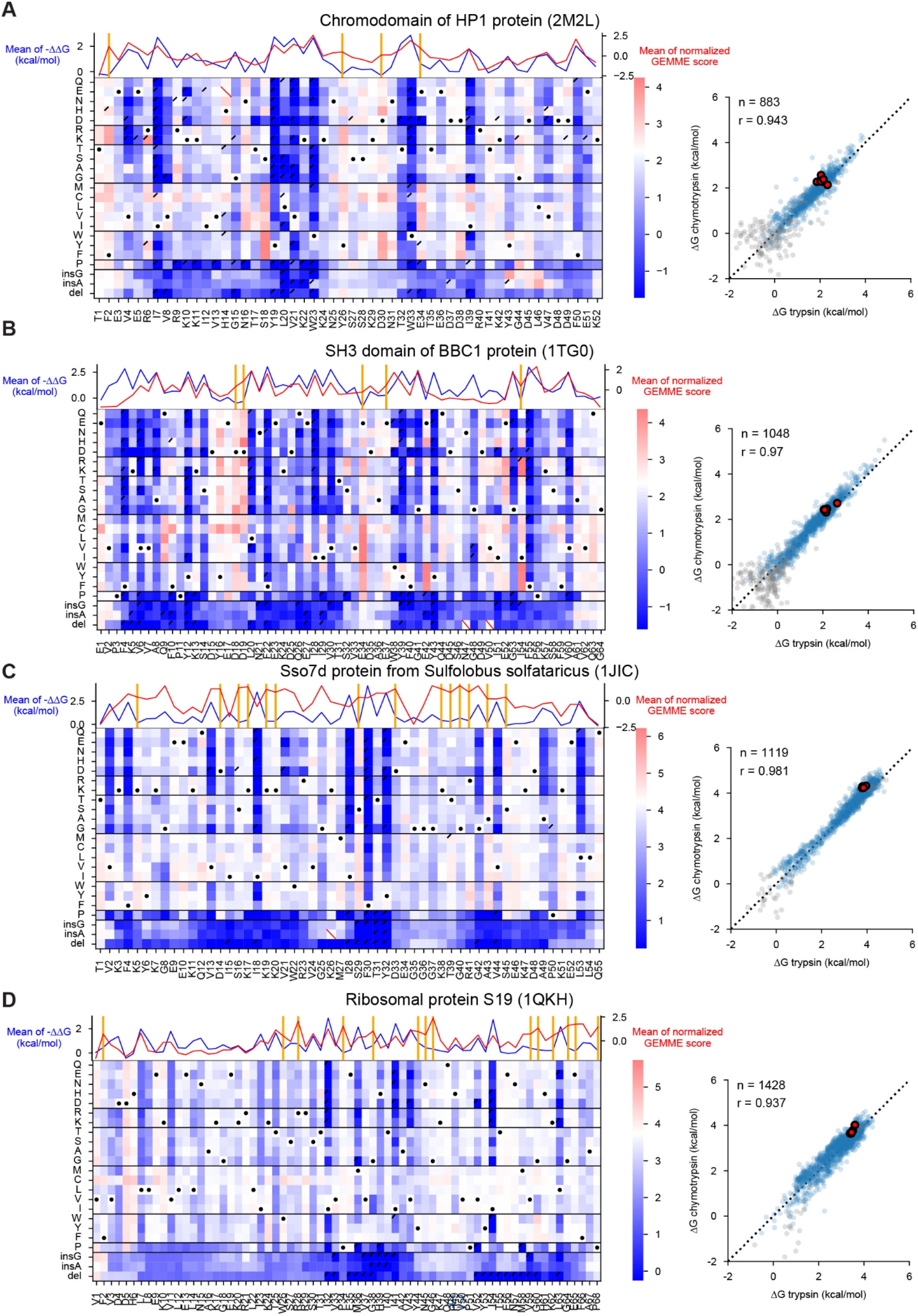
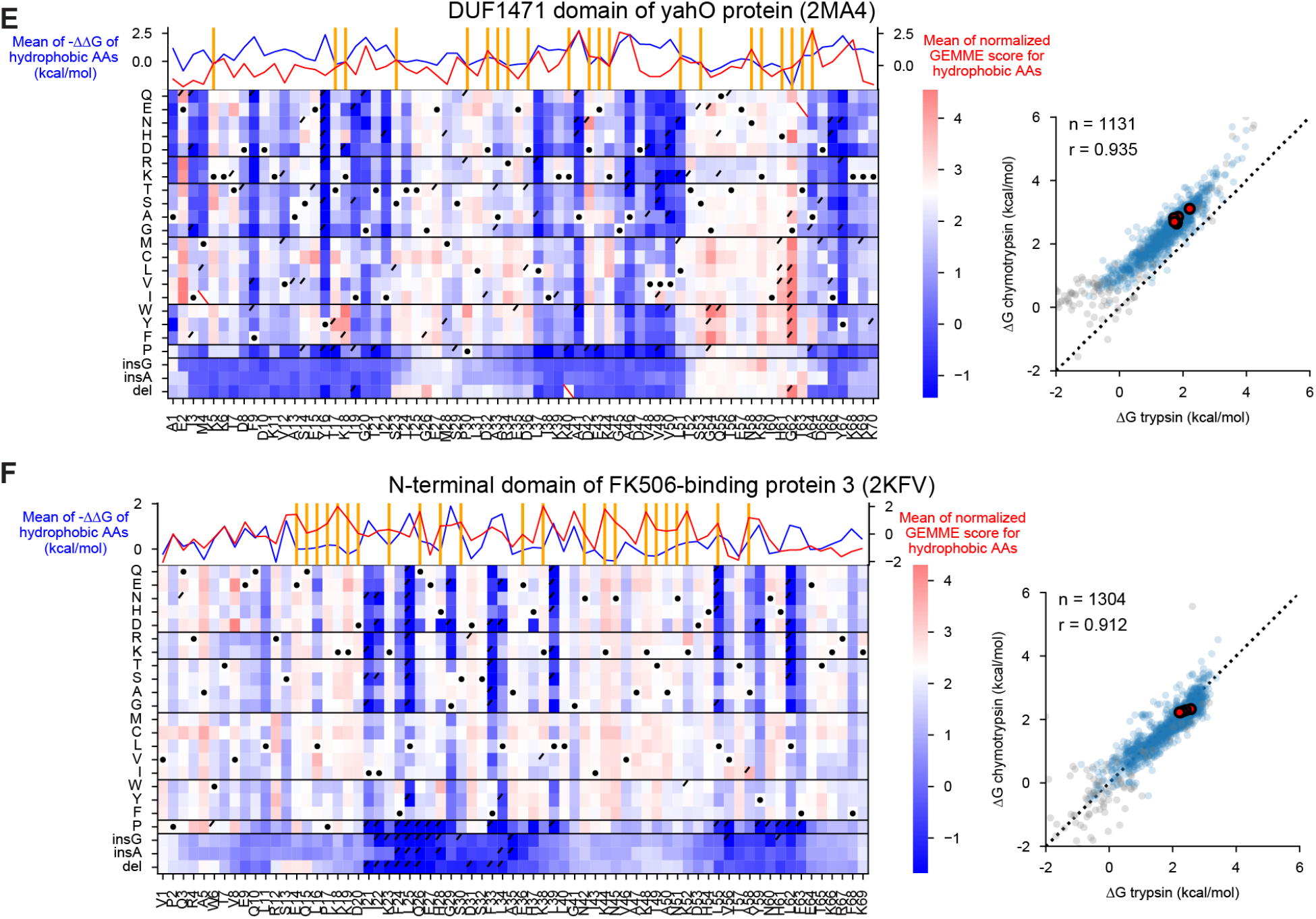
Heat maps for notable domains with functional residues. **(A- D)** Mutational scanning results for four domains. Left: Heat maps show ΔG for substitutions, deletions, and Gly and Ala insertions at each residue. White indicates the wild-type stability and red/blue indicates stabilizing/destabilizing. Black dots indicate the wild-type amino acid, red slashes indicate missing data, and black corner slashes indicate lower confidence ΔG estimates, (95% confidence interval > 0.5 kcal/mol), including ΔG estimates near the edges of the dynamic range. At top, lines show the mean ΔΔG (blue) and the mean normalized GEMME score (red), with functional sites (classified according to Fig. 6A) marked with vertical orange lines. Right: Agreement between variant ΔG values independently determined using assays with trypsin (x-axis) and chymotrypsin (y-axis). Multiple codon variants of the wild-type sequence are shown in red, reliable ΔG values in blue, and less reliable ΔG estimates (same as above) in gray. The black dashed line represents Y=X. Each plot shows the number of reliable points and the Pearson r-value for the blue (reliable) points. **(E- F)** Same as (A-D), but top lines indicate the mean of ΔΔG for hydrophobic amino acid substitutions (blue) and mean normalized GEMME score of hydrophobic amino acids (red). Functional sites are classified according to Fig. 6G.

**Fig. S17.**
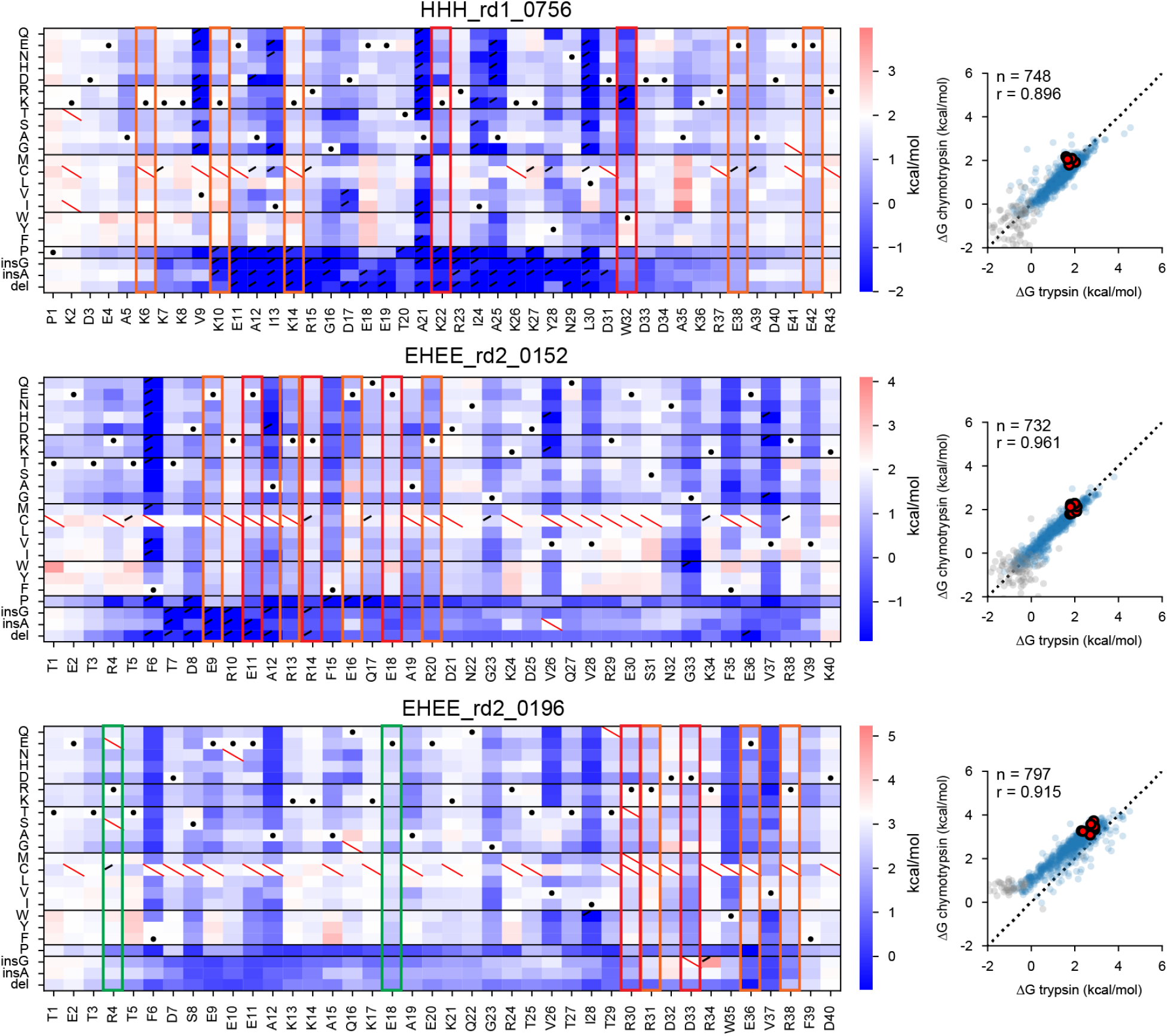
Heat maps for three designed domains with notable polar interactions. Mutational scanning results for three domains with notable polar interactions. Left: Heat maps show ΔG for substitutions, deletions, and Gly and Ala insertions at each residue. White indicates the wild-type stability and red/blue indicates stabilizing/destabilizing. Black dots indicate the wild-type amino acid, red slashes indicate missing data, and black corner slashes indicate lower confidence ΔG estimates, (95% confidence interval > 0.5 kcal/mol), including ΔG estimates near the edges of the dynamic range. The polar networks shown in Fig. 7B are highlighted in orange, red, and green. Right: Agreement between variant ΔG values independently determined using assays with trypsin (x-axis) and chymotrypsin (y-axis). Multiple codon variants of the wild-type sequence are shown in red, reliable ΔG values in blue, and less reliable ΔG estimates (same as above) in gray. The black dashed line represents Y=X. Each plot shows the number of reliable points and the Pearson r-value for the blue (reliable) points.

**Fig. S18.**
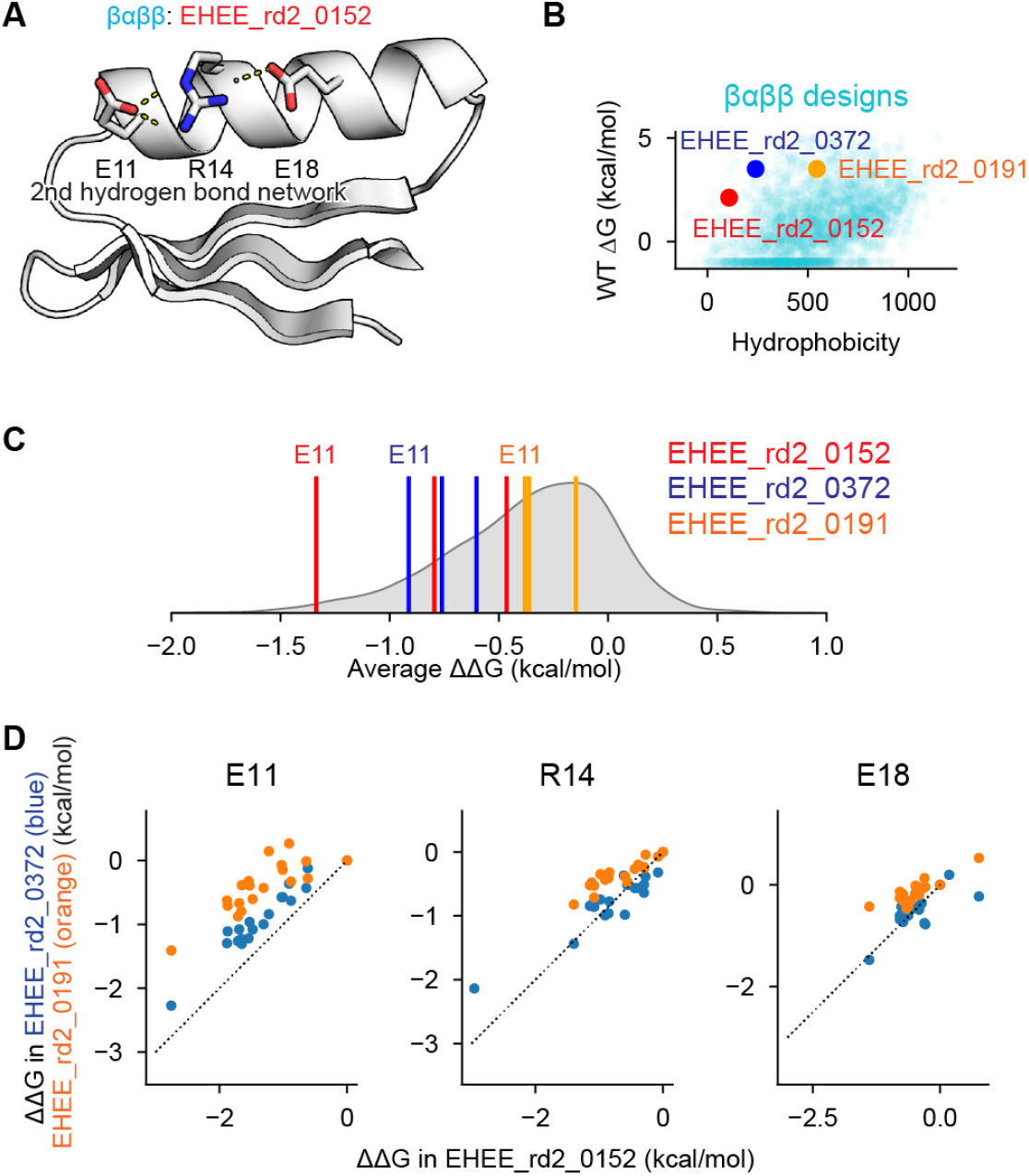
Comparison of the notable hydrogen bond networks in three designs. **(A)** Designed structure of EHEE_rd2_0152 from (*29*) highlighting residues in the “2nd hydrogen bond network” defined in Fig. 7B middle. **(B)** Relationship between hydrophobicity (calculated based on (*117*)) and folding stability (ΔG) for designed βαββ proteins. The three dots in the plot represent three designs with the same hydrogen bond network. **(C)** The gray density plots represent the average ΔΔG of substitutions at 3,715 polar sites in 144 designed domains. The colored vertical bars indicate the values for the sites related to the 2nd hydrogen bond network. **(D)** Relationship between ΔΔG in EHEE_rd2_0152 and in the other designs EHEE_rd2_0372 or EHEE_rd2_0191 for E11, R14, and E18. At E11, substitutions to the 19 other amino acids have smaller effects in EHEE_rd2_0372 (blue) and EHEE_rd2_0191 (orange) compared to in EHEE_rd2_0152 (e.g. all points are above the dashed Y=X line). However, the points are ordered similarly; i.e. the rank ordering of the 19 other amino acid variants in stability is similar between the three designs. For R14 and E18, substitutions in EHEE_rd2_372 (blue) have similar effect sizes to EHEE_rd2_0152, but substitutions in EHEE_rd2_0191 (orange) have smaller effects. Again, the rank ordering of the amino acid variants by stability is similar across the three designs.

**Fig. S19.**
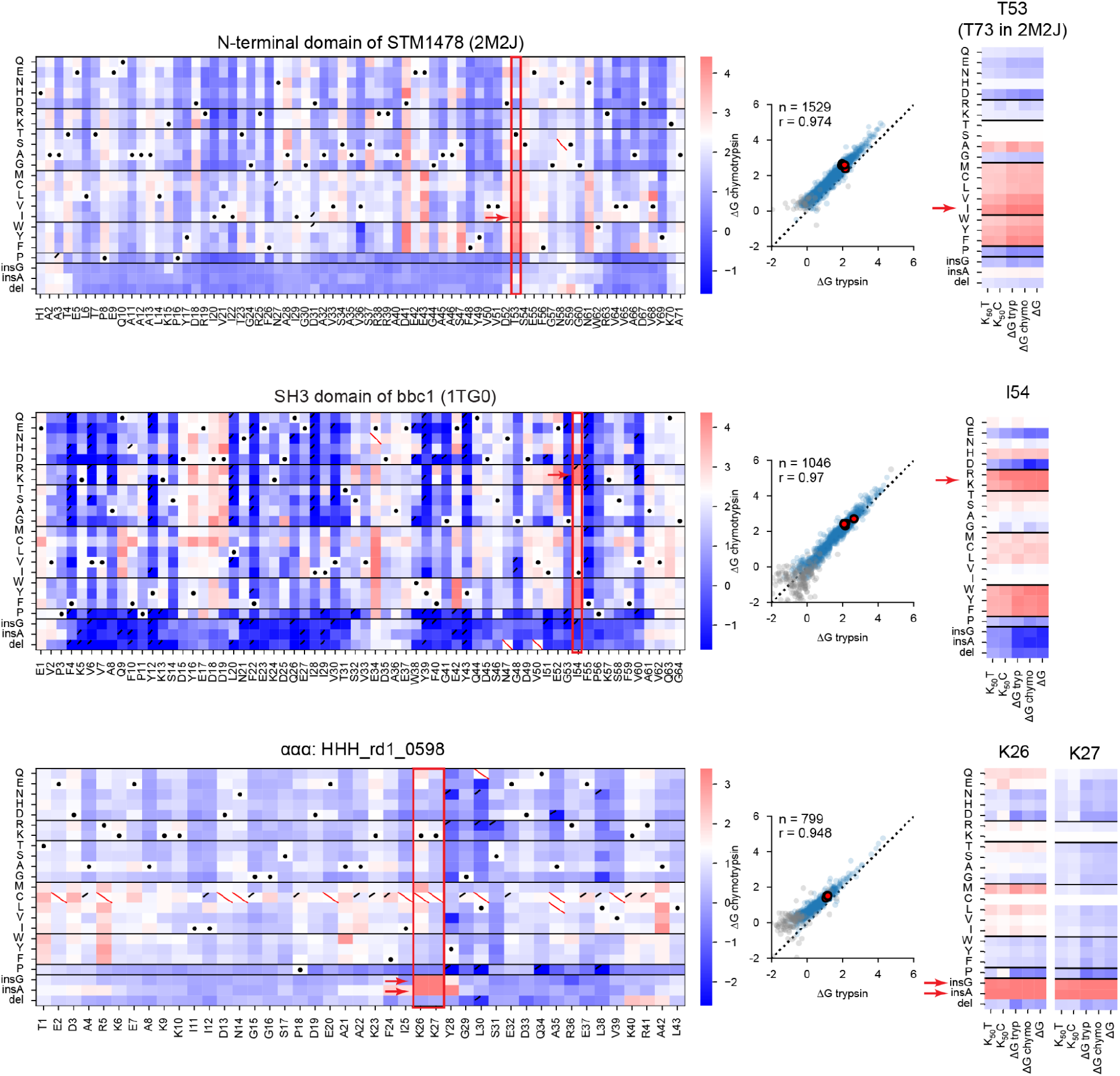
Heat maps for three domains with notable stabilizing mutations. Left: Heat maps show ΔG for substitutions, deletions, and Gly and Ala insertions at each residue. White indicates the wild-type stability and red/blue indicates stabilizing/destabilizing. Black dots indicate the wild-type amino acid, red slashes indicate missing data, and black corner slashes indicate lower confidence ΔG estimates, (95% confidence interval > 0.5 kcal/mol), including ΔG estimates near the edges of the dynamic range. The red boxes and arrows highlight sites with notable stabilizing mutations. Middle: Agreement between variant ΔG values independently determined using assays with trypsin (x-axis) and chymotrypsin (y-axis). Multiple codon variants of the wild-type sequence are shown in red, reliable ΔG values in blue, and less reliable ΔG estimates (same as above) in gray. The black dashed line represents Y=X. Each plot shows the number of reliable points and the Pearson r-value for the blue (reliable) points. Right: For four positions with stabilizing mutations, heatmaps show five experimental metrics: the trypsin (T) and chymotrypsin (C) K_50_ values, the ΔG values inferred from trypsin and chymotrypsin experiments, and the overall ΔG inferred from both trypsin and chymotrypsin experiments together.

**Fig. S20.**
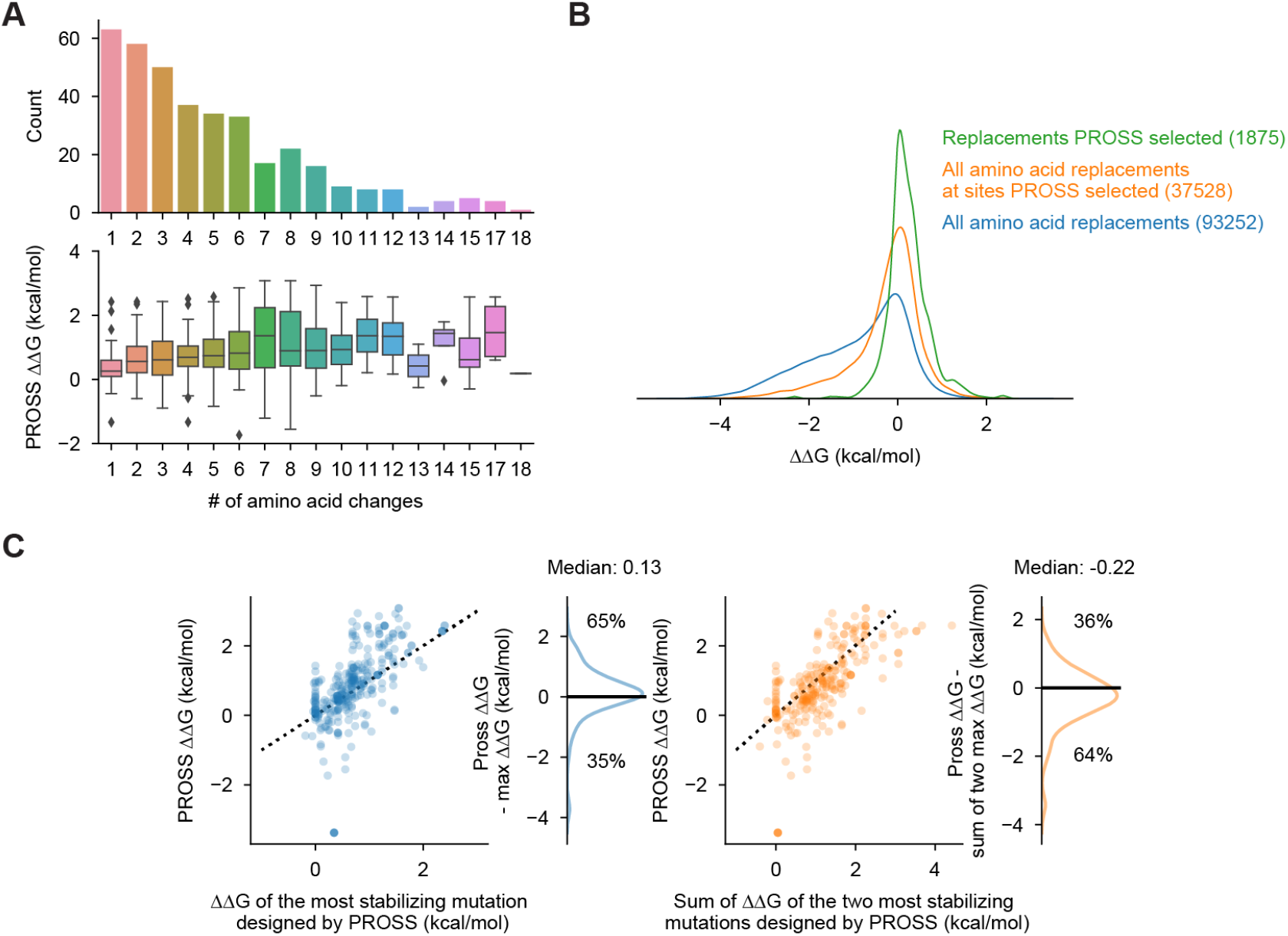
Global analysis of PROSS designs. **(A)** All 727 PROSS designs grouped according to the number of amino acid substitutions in each design. Top: the number of designs with each different number of substitutions. Bottom: the distribution of design results for each group. ΔΔG indicates the stability of the PROSS design (ΔG) minus the stability of the original wild-type sequence; positive ΔΔG indicates the design stabilized the domain. **(B)** ΔΔG distributions for all amino acid substitutions in wild-type domains used as input to PROSS (blue), all amino acid substitutions at sites modified in PROSS designs (orange), and all PROSS-designed substitutions (green). All ΔΔG measurements are in the original wild-type background; positive ΔΔG indicates stabilizing substitutions. **(C)** Relationship between ΔΔG of PROSS designs and ΔΔG of the most stabilizing mutations designed by PROSS. At left, we compare PROSS designs to the single most stabilizing mutation (in the original wild-type background) out of all the substitutions in the PROSS design. At right, we compare PROSS designs to the sum of the two most stabilizing mutations (each measured individually in the original wild-type background without considering thermodynamic coupling). The density plots show the distribution of PROSS designs that were better (positive) or worse (negative) than the single best mutation (left) or sum of the two best mutations (right). Two-thirds of designs are better than the best single designed mutation, although the difference is small. Likewise, two-thirds of designs are worse than the additive effect of the two best designed mutations (assuming no thermodynamic coupling).

**Table S1.** List of amino acid sequence number for each group.

| **Dataset name** | **# of total sequences** | **Sequence group** | **# of sequence groups** | **# of sequences** |
| --- | --- | --- | --- | --- |
| **Dataset #1** | 586,938 | Single amino acid replacement, deletion, and insertion | 396 wild-types | 434,556 |
|  |  | Double amino acid replacement | 458 pairs | 152,382 |
| **Dataset #2** | 851,552 | Single amino acid replacement, deletion, and insertion | 560 wild-types | 629,287 |
|  |  | Double amino acid replacement | 595 pairs | 222,265 |
| **Dataset #3** | 1,844,548 | Single amino acid replacement, deletion, and insertion | 983 wild-types | 1,046,752 |
|  |  | Double/Triple amino acid replacement | 725 pairs (including 36 triples) | 416,274 |
|  |  | Scrambles for unfolded model | - | 68,427 |
|  |  | Rocklin 2017 rd1-3 | - | 36,707 |
|  |  | Others | - | 276,388 |

**Table S2.** Conditions for measuring folding stability in the previous papers.

| **Protein** | **PDB ID** | **Pubmed ID** | **Reference** | **Offset in Fig.1G (kcal/mol)** | **Temperature (°C)** | **Buffer** | **pH** |
| --- | --- | --- | --- | --- | --- | --- | --- |
| Protein GB1 | 1PGA | 31371509 | (52) | -0.8 | 25 | NaPi | 6.5 |
| NTL9 | 2HBB | 28494951 | (53) | 1.9 | 25 | 20 mM NAOAc, 100 mM NaCl | 5.5 |
| SAP domain | 2WQG | 26073259 | (54) | 1.2 | 10 | 50 mM MES pH 6.0, 500 mM NaCl | 6 |
| hPin1 WW domain | 2M8I | 19565466 | (55) | -1.9 | 40 | 10 mM NaPi | 7 |
| hYap65 WW domain | 1K9Q | 11420447 | (56) | -2.3 | 60 | 20 mM KPi, 100 mM NaCl | 7 |
| hYap65 WW domain | 1K9Q | 23035249 | (57) | -2.3 | 50 | 20 mM NaPi | 7 |
| Villin HP35 | 1VII | 23798426/19354264 | (58, 59) | 0.1 | 25 | 10 mM NaOAc, 150 mM NaCl | 5 |
| BBL | 2WAV | 19445954 | (60) | 0.3 | 10 | 50 mM KPi 200 mM KCl | 7 |
| FF domain | 1UZC | 15935381 | (61) | 0.8 | 10 | 50 mM NAOAc, 100 mM NaCl | 5.7 |
| ADA2h activation domain | 1O6X | 9799641 | (62) | -0.2 | 25 | 50 mM NaPi | 7 |
| hFyn SH3 domain | 1SHF | 9819209/12079394/16142914 | (63–65) | -0.1 | 25 | 10 mM Tris, 0.2 mM EDTA and 250 mM KCl | 8 |

**Table S3.** Description of metrics in Fig. S8.

| **Metrics** | **Description** |
| --- | --- |
| dg_corr | Correlation between $\Delta G$ from trypsin challenge and chymotrypsin challenge |
| frac_pos_with_hydrophobic_stabilizing_muts | Fraction of stabilizing mutations which replace hydrophobic amino acids |
| frac_stabilizing_mut | Fraction of stabilizing mutations |
| lowconf_frac_lowss | Fraction of mutations with low reliable data (95CI > 0.5 kcal/mol) |
| NA_frac | Fraction of data without sufficient NGS reads |
| raw_corr | Correlation between trypsin K50 and chymotrypsin K50 |
| slope | Slope of linear fitting line between $\Delta G$ from trypsin challenge and chymotrypsin challenge |
| width_KC | Standard deviation of $\Delta G$ from chymotrypsin challenge |
| width_KT | Standard deviation of $\Delta G$ from trypsin challenge |
| wt_d_from_line | Distance between WT data points and linear fitting line from mutational data |
| wt_dg | $\Delta G$ of WT |
| wt_dg_std_max | Standard deviation of $\Delta G$ of WT |
| y_intercept | Y-intercept of linear fitting line between $\Delta G$ from trypsin challenge and chymotrypsin challenge |

**Table S4.** Description of columns in K50_dG_Dataset1_Dataset2.

| Column name | Description |
| --- | --- |
| name | sequence name |
| dna_seq | DNA sequence |
| log10_K50_t | Median of posteriors of K<sub>50</sub> trypsin in log10 scale (μM) |
| log10_K50_t_95CI_high | Top 2.5%ile of posterior of K<sub>50</sub> trypsin in log10 scale (μM) |
| log10_K50_t_95CI_low | Top 97.5%ile of posterior of K<sub>50</sub> trypsin in log10 scale (μM) |
| log10_K50_t_95CI | log10_K50_t_95CI_high - log10_K50_t_95CI_low |
| fitting_error_t | Absolute error between the observed counts and the expected counts for a given sequence (based on all model parameters related to trypsin data), averaged over 24 conditions and normalized by the observed counts in the no-protease samples for that sequence |
| log10_K50unfolded_t | K<sub>50</sub> unfolded trypsin in log10 scale (μM) |
| deltaG_t | ΔG calculated from log10_K50_t (kcal/mol) |
| deltaG_t_95CI_high | ΔG calculated from log10_K50_t_95CI_high (kcal/mol) |
| deltaG_t_95CI_low | ΔG calculated from log10_K50_t_95CI_low (kcal/mol) |
| deltaG_t_95CI | deltaG_t_95CI_high - deltaG_t_95CI_low (kcal/mol) |
| log10_K50_c | Median of posteriors of K<sub>50</sub> chymotrypsin in log10 scale (μM) |
| log10_K50_c_95CI_high | Top 2.5%ile of posterior of K<sub>50</sub> chymotrypsin in log10 scale (μM) |
| log10_K50_c_95CI_low | Top 97.5%ile of posterior of K<sub>50</sub> chymotrypsin in log10 scale (μM) |
| log10_K50_c_95CI | log10_K50_c_95CI_high - log10_K50_c_95CI_low |
| fitting_error_c | Absolute error between the observed counts and the expected counts for a given sequence (based on all model parameters related to chymotrypsin data), averaged over 24 conditions and normalized by the observed counts in the no-protease samples for that sequence |
| log10_K50unfolded_c | K<sub>50</sub> unfolded chymotrypsin in log10 scale (μM) |
| deltaG_c | ΔG calculated from log10_K50_c (kcal/mol) |
| deltaG_c_95CI_high | $\Delta G$ calculated from log10_K50_c_95CI_high (kcal/mol) |
| deltaG_c_95CI_low | $\Delta G$ calculated from log10_K50_c_95CI_low (kcal/mol) |
| deltaG_c_95CI | deltaG_c_95CI_high - deltaG_c_95CI_low (kcal/mol) |
| deltaG | Median of posterior of $\Delta G$ from trypsin+chymotrypsin data (kcal/mol) |
| deltaG_95CI_high | Top 2.5%ile posterior of $\Delta G$ from trypsin+chymotrypsin data (kcal/mol) |
| deltaG_95CI_low | Top 97.5%ile posterior of $\Delta G$ from trypsin+chymotrypsin data (kcal/mol) |
| deltaG_95CI | deltaG_95CI_high - deltaG_95CI_low |
| aa_seq_full | Amino acid sequence including padding linker sequence |
| aa_seq | Amino acid sequence without linker sequence |
| mut_type | Mutation type (like WT, substitution, insertion, deletion, or double mutants) |
| WT_name | Name of wild-type domain |
| WT_cluster | Cluster number of wild-type domain |
| log10_K50_trypsin_ML | K<sub>50</sub> trypsin in log10 scale for machine learning ($\mu M$) ('-' means lacking or unreliable data) |
| log10_K50_chymotrypsin_ML | K<sub>50</sub> chymotrypsin in log10 scale for machine learning ($\mu M$) ('-' means lacking or unreliable data) |
| dG_ML | $\Delta G$ for machine learning (kcal/mol) ('-' means unreliable data) |
| ddG_ML | $\Delta\Delta G$ for machine learning (kcal/mol) ('-' means unreliable data) |
| Stabilizing_mut | True if $\Delta\Delta G > 1$ (kcal/mol) and $\Delta\Delta G$ is reliable |

### Files for Supplementary Materials

Raw_NGS_count_tables.zip

NGS_count_lib1.csv
NGS_count_lib2.csv
NGS_count_lib3.csv
NGS_count_lib4.csv
K50_dG_tables.zip

K50_dG_lib1.csv
K50_dG_lib2.csv
K50_dG_lib3.csv
K50_dG_lib4.csv
Processed_K50_dG_datasets.zip

K50_dG_Dataset1_Dataset2.csv
K50_Dataset3.csv
Single_DMS_list.csv
Double_DMS_list.csv
Triple_DMS_list.csv
Heat_maps_single_DMS.pdf
Heat_maps_double_DMS.pdf
Data_tables_for_figs.zip

dG_extdG_data_Fig1.csv
dG_site_feature_Fig3.csv
dG_for_double_mutants_Fig4.csv
dG_non_redundant_natural_Fig5.csv
dG_GEMME_non_redundant_natural_Fig6.csv
Pipeline_qPCR_data.zip

Raw_qPCR_data_FigS1.csv
Process_qPCR_data.ipynb
Pipeline_K50_dG.zip

STEP1_module.ipynb
STEP1_run.ipynb
STEP2_run.ipynb
STEP3_run.ipynb
STEP4_module.ipynb
STEP4_run.ipynb
STEP5_module.ipynb
STEP5_run.ipynb
Raw_NGS_counts_overlapped_seqs_STEP1_libl_lib2.csv
Raw_NGS_counts_overlapped_seqs_STEP1_lib2_lib3.csv
Raw_NGS_counts_overlapped_seqs_STEP1_libl_lib4.csv
Raw_NGS_counts_overlapped_seqs_STEP1_lib2_lib4.csv
Raw_NGS_counts_overlapped_seqs_STEP1_lib3_lib4.csv
K50_scrambles_for_STEP3.csv
STEP1_out_protease_concentration_trypsin
STEP1_out_protease_concentration_chymotrypsin
STEP3_unfolded_model_params
Pipeline_figure_model.zip

Burial_side_chain_contact_Fig3_Fig6.ipynb
Additive_model_Fig4.ipynb
Classification_model_Fig5.ipynb
AlphaFold_model_PDBs.zip
Blueprints_for_EEHH.zip

eehh_EA_GBB_AGBB.bp
eehh_GG_GBB_AGBB.bp
eehh_XX_XXX_XXXX.bp

